# Time-resolved comparative molecular evolution of oxygenic photosynthesis

**DOI:** 10.1101/2020.02.28.969766

**Authors:** Thomas Oliver, Patricia Sánchez-Baracaldo, Anthony W. Larkum, A. William Rutherford, Tanai Cardona

## Abstract

Oxygenic photosynthesis starts with the oxidation of water to O_2_, a light-driven reaction catalysed by photosystem II. Cyanobacteria are the only prokaryotes capable of water oxidation and therefore, it is assumed that relative to the origin of life and bioenergetics, the origin of oxygenic photosynthesis is a late innovation. However, when exactly water oxidation originated remains an unanswered question. Here we use relaxed molecular clocks to compare one of the two ancestral core duplications that are unique to water-oxidizing photosystem II, that leading to CP43 and CP47, with some of the oldest well-described events in the history of life. Namely, the duplication leading to the Alpha and Beta subunits of the catalytic head of ATP synthase, and the divergence of archaeal and bacterial RNA polymerases and ribosomes. We also compare it with more recent events such as the duplication of cyanobacteria-specific FtsH metalloprotease subunits, of CP43 variants used in a variety of photoacclimation responses, and the speciation events leading to Margulisbacteria, Sericytochromatia, Vampirovibrionia, and other clades containing anoxygenic phototrophs. We demonstrate that the ancestral core duplication of photosystem II exhibits patterns in the rates of protein evolution through geological time that are nearly identical to those of the ATP synthase, RNA polymerase, or the ribosome. Furthermore, we use ancestral sequence reconstruction in combination with comparative structural biology of photosystem subunits, to provide additional evidence supporting the premise that water oxidation had originated before the ancestral core duplications. Our work suggests that photosynthetic water oxidation originated closer to the origin of life and bioenergetics than can be documented based on species trees alone.

## 1. Introduction

### 1.1. Evolution of Cyanobacteria

The origin of oxygenic photosynthesis is considered a turning point in the history of life, marking the transition from the ancient world of anaerobes into a productive aerobic world that permitted the emergence of complex life [1]. Oxygenic photosynthesis starts with photosystem II (PSII), the water-oxidising and O_2_-evolving enzyme of Cyanobacteria and photosynthetic eukaryotes. Today PSII is a highly conserved, multicomponent, membrane protein complex, which was inherited by the *most recent common ancestor* (MRCA) of Cyanobacteria in a form that is structurally and functionally quite similar to that found in all extant species [2]. Thus, the origin of oxygenic photosynthesis antedates the MRCA of Cyanobacteria by an undetermined amount of time.

Cyanobacteria’s closest living relatives are the clades known as Vampirovibrionia (formerly Melainabacteria) [3, 4], followed by Sericytochromatia [5] and Margulisbacteria [6]. Currently, no photosynthetic strains have been described in these groups of uncultured bacteria and this has led to the hypothesis that oxygenic photosynthesis arose during the time spanning the divergence of Vampirovibrionia and the MRCA of Cyanobacteria, starting from an entirely non-photosynthetic ancestral state [5, 7]. Molecular clock studies have suggested that the span of time between the divergence of Cyanobacteria and Vampirovibrionia could be of several hundred million years [8, 9]. However, it is still unclear from molecular clock analyses and the microfossil record when exactly the MRCA of Cyanobacteria occurred [10]. For example, two recent independent analysis placed the same cyanobacterial ancestor around a mean age younger than 1.5 Ga [11] and older than 3.5 Ga [12].

### 1.2. Evolution of photosystem II

The heart of PSII is made up of a heterodimeric reaction centre (RC) *core* coupled to a core *antenna.* The two subunits of the RC core of PSII are known as D1 and D2, and these are associated respectively with the antenna subunits known as CP43 and CP47. D1 and CP43 make up one monomeric half of the RC, and D2 and CP47, the other half. Water oxidation is catalysed by a Mn4CaO5 cluster coordinated by ligands from both D1 and CP43 [13, 14]. The cluster is functionally coupled to a redox active tyrosine-histidine pair (Y_Z_-H190) also located in D1, which relays electrons from Mn to the oxidised chlorophyll pigments of the RC during charge separation [15]. In a cycle of four consecutive light-driven charge separation events, O2 is released in the decomposition of two water molecules.

Photosystems evolved first as homodimers [16, 17]: therefore, the core and the antenna of PSII originated from ancestral gene duplication events that antedated the MRCA of Cyanobacteria. In this way, CP43/D1 retain sequence and structural identity with CP47/D2. The conserved structural and functional traits between CP43/D1 and CP47/D2 suggest that the ancestral PSII homodimer—prior to the duplication events—was not only structurally similar to heterodimeric PSII, but also that it split water and had evolved protective mechanisms against the formation of reactive oxygen species [17–19].

In a previous study, we attempted to measure the span of time between the duplication that led to D1 and D2 (*d*D0) and a point that approximated the MRCA of Cyanobacteria: a period of time that we called **ΔT** [18]. We observed that the magnitude of ΔT can be very large, well over a billion years. Such large ΔT suggested that the origin of a water-oxidising photosystem antedated Cyanobacteria themselves and potentially other groups of Bacteria depending on how rapidly the domain diversified. This implies that an ancestral nonphotosynthetic state at the divergence of Margulisbacteria, Sericytochromatia, and Vampirovibrionia (MSV) cannot be taken for granted despite their specialized heterotrophic lifestyles. However, our study neither provided an absolute age for the MRCA of Cyanobacteria nor the duplication event itself, as we simulated a comprehensive range of scenarios. Instead, we showed that even when ΔT is over a billion years, the rate of protein evolution at the duplication point (*d*D0) needed to be over 40 times greater than any rate ever observed for D1 and D2, decreasing exponentially during the Archean and stabilising at current rates during the Proterozoic. Thus, the shorter the ΔT, the faster the rate at *dD0,* with the rate increasing following a power law function and reaching unrealistic values even when ΔT is still in the order of several hundred million years [18]. It was still unclear if such patterns of evolution were unique to the core subunits of PSII or whether other systems have experienced similar evolutionary trajectories.

Here, to help in understanding the evolution of oxygenic photosynthesis, Cyanobacteria and MSV as a function of time, we compared the duplication leading to the RC antenna subunits, CP43 and CP47, to several well-defined ancient and more recent events: including, but not limited to, the duplication of the core catalytic subunits of ATP synthase, a very ancient event generally accepted to have occurred before the last universal common ancestor (LUCA) [20–24]; the evolution of RNA polymerase catalytic subunit β (RpoB) and ribosomal proteins, which are universally conserved and widely accepted to have originated before the LUCA [25–28]. We further constrain our analysis using *in silico* ancestral sequence reconstruction of PSII and through strict structural and functional rationales. We show that the core subunits of PSII show molecular signatures that are usually associated with some of the oldest transitions in the molecular evolution of life. We also show that all events leading from the primordial homodimeric water-splitting photosystem to Cyanobacteria’s heterodimeric PSII can be reconstructed from a comparison of available structural and genetic data.

## 2. Results

### 2.1. Phylogenetic overview

The phylogenies of CP43 and CP47 show that there is a much greater diversity of CP43 and CP43-derived subunits than CP47 (Figure 1). This difference is the result of a greater number of gene duplication in CP43 than CP47 and mirrors the evolution of D1 and D2 [2], in which D1 has undergone more duplication events than D2. CP43 can be divided into two major groups: those that are assembled into PSII and can bind the Mn4CaO5 cluster, and those which have evolved to be used only as light harvesting complexes [29, 30], usually known as chlorophyll binding proteins (CBP). The CBP are characterized by the loss of the extrinsic loop between the 5^th^ and 6^th^ transmembrane helices, where the ligands to the cluster are located (Figure 1b). This large extrinsic loop is found in both CP43 and CP47 and interacts directly with the electron donor side of PSII, within D1 and D2 respectively. The unrooted tree of CP43 is consistent with CBP having a single origin likely occurring before the MRCA of Cyanobacteria (Supplementary Figure S1) but have undergone an extensive duplication-driven diversification process. It also mirrors the evolution of D1 in that duplications appear to have occurred before the MRCA of Cyanobacteria [2]. The earliest of these D1 duplications also led to variants that have lost the capacity to bind the Mn_4_CaO_5_ cluster [2, 31], but are likely used in other supporting functions. Most notably, during chlorophyll *f* synthesis [32] in the far-red light acclimation response (farlip) [33].

**Figure 1.**
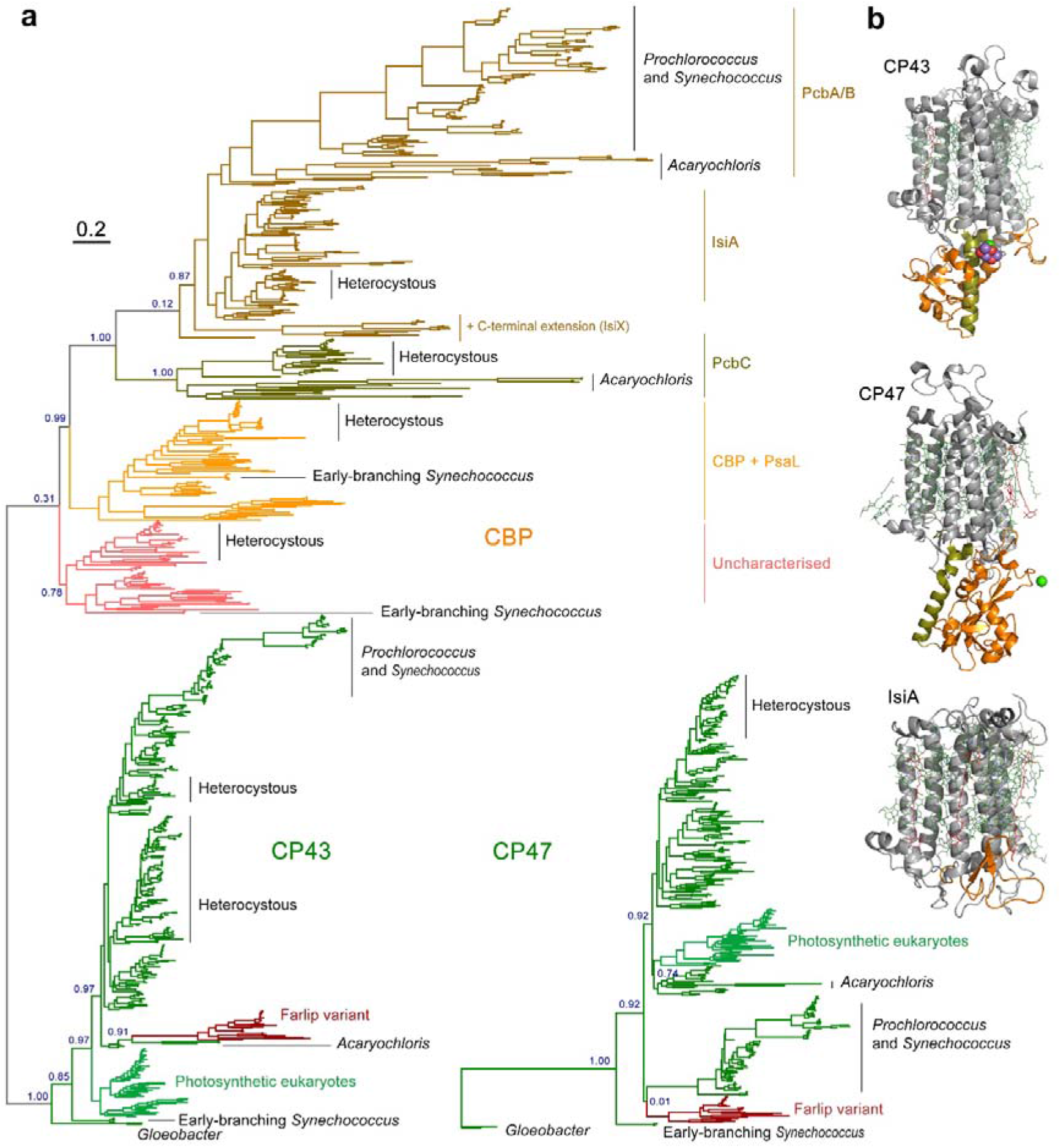
Maximum Likelihood trees of PSII core antenna subunits and derived light harvesting proteins. **a** A tree of CP43 and chlorophyll binding proteins (CBP). The unrooted tree splits into CP43 and CBP, with CP43 displaying a phylogeny roughly consistent with the evolution of Cyanobacteria, although several potential duplications of CP43 are noticeable within heterocystous Cyanobacteria and closer relatives. Farlip variant denotes the isoform used in the far-red light acclimation response. CBP shows a complex phylogeny strongly driven by gene duplication events and fast evolution. The tree of CP47 was rooted at the divergence of *Gloeobacter spp.* Scale bar represents number of substitutions per site. **b** Structural comparison of CP43, CP47, and IsiA. The latter lacks the characteristic extrinsic loop (orange) of CP43 and CP47 that links the antenna with the electron donor side of PSII.

CP43 and CP47 are also distantly related to the antenna domain of cyanobacterial PSI and that of the type I RCs of phototrophic Chlorobi, Acidobacteria and Heliobacteria (Supplementary Figure S2). We found that the level of sequence identity between any two type I RC proteins is always greater than between a type I RC protein and CP43/CP47 (Supplementary Table S1). Therefore, the distance between CP43/CP47 and other type I antenna domains is the largest distance in the molecular evolution of RC proteins after that between type I and type II RC core domains. The phylogenetic relationships between type I and type II RCs have been reviewed in detail before [19, 34, 35]. These are briefly summarized and schematized in Supplementary Figure S3.

The phylogeny of Alpha and Beta subunits of the F-type ATP synthase showed that all Cyanobacteria have a F-type ATP synthase, and a fewer number of strains have an additional Na^+^-translocating ATPase (N-ATPase) of the bacterial F-type, as had been reported before [36]. We found that MSV have a standard F-type ATP synthase (Supplementary Figure S4), but some N-ATPase Alpha and Beta sequences were also found in Vampirovibrionia and Sericytochromatia datasets, but not in Margulisbacteria. In this study we focused on the standard F-type ATP synthase of Cyanobacteria for further analysis.

The phylogeny of bacterial RNA polymerase subunit β (RpoB), with the intention of molecular clock analysis, focused on Cyanobacteria and MSV, as well as phyla with known phototrophic representatives and included Thermotogae and Aquificae as potential roots (Supplementary Figure S5). The tree was largely consistent with previous observed relationships between the selected groups [37], within Cyanobacteria and MSV, and within other phototrophs and their non-phototrophic relatives [38, 39]. The only exception was Aquificae, which branched as a sister clade to Acidobacteria, a feature that had been reported before for RpoB [27], and likely represents an ancient horizontal gene transfer event.

### 2.2. Distances and rates

To gather temporal information, we compared the phylogenetic distances between CP43 and CP47, Alpha and Beta subunits of the F-type ATP synthase, and archaeal and bacterial RpoB (visualized in Figure 2, but see also Figures S1b, S2 and S3b). We found that the distances between Alpha and Beta, and the divergence of archaeal and bacterial RpoB, are very large relative to the distance between the divergence of Vampirovibrionia and Cyanobacteria. In the case of RpoB, the distance between Vampirovibrionia and Cyanobacteria is about a fifth of the distance between Archaea and Bacteria. However, the distance between CP43 and CP47 (but also between D1 and D2 [18]) is of similar magnitude to that between Alpha and Beta, and to that between archaeal and bacterial RpoB, but substantially surpasses the distance between MVS and Cyanobacteria (Figure 2). These observations suggest that ancestral proteins to CP43/CP47 and D1/D2 existed before the divergences of MVS.

**Figure 2.**
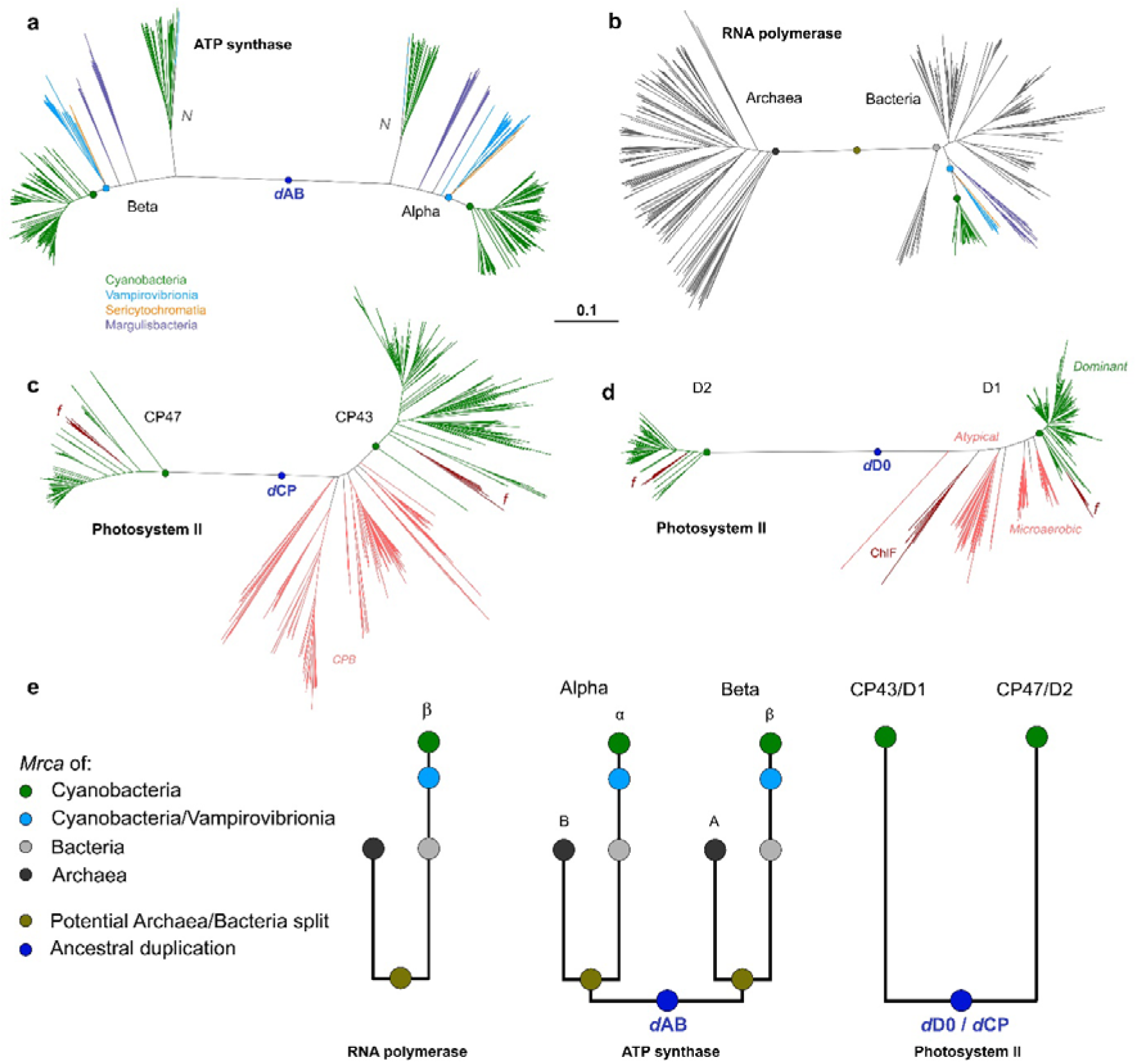
Distance comparison of the core subunits of ATP synthase, RNA polymerase, and PSII. **a** Alpha and Beta subunits of the F-type ATP synthase from Cyanobacteria, Vampirovibrionia, Sericytochromatia and Margulisbacteria. *N* denotes Na^+^ translocating N-type ATPase. *d*AB denotes the duplication event leading to Alpha and Beta. The green dot marks the MRCA of Cyanobacteria, and the light-blue dot the MRCA of the clade including Cyanobacteria and Vampirovibrionia. **b** Archaeal and Bacterial RpoB of RNA polymerase. **c** CP43 and CP47 subunits of PSII. *f* denotes farlip variants. *d*CP marks the duplication event leading to CP43 and CP47. Pink branches highlight the CPB subunits. **d** D1 and D2 of PSII showing a pattern that mimics that of CP43 and CP47. Pink branches denote the atypical D1 forms and other variants that are thought to predate the MRCA of Cyanobacteria. The sequences marked with *dominant* represent the standard D1 form of PSII, inherited by the MRCA of Cyanobacteria and found in all oxygenic phototrophs [2, 18]. ChlF marks the atypical D1 form involved in the synthesis of chlorophyll *f* during farlip [32]. *d*D0 marks the duplication leading to D1 and D2. **e** Schematic representation of distance and distribution of these enzymes in Cyanobacteria and relatives in relation to the MRCA of Bacteria, of Archaea, and the LUCA.

We compared the within-group mean distances for Alpha, Beta, RpoB, and a concatenated dataset of ribosomal proteins compiled in a previous independent study [38] (see Supplementary Table S2). We found consistently, that Vampirovibrionia and Margulisbacteria have larger within-group mean distances compared to Cyanobacteria, which suggests greater rates of evolution in the non-photosynthetic clades. These were consistently larger for Margulisbacteria relative to the other two groups. For example, RpoB in Vampirovibrionia and Margulisbacteria showed 1.6x and 4.0x larger corrected mean distances than Cyanobacteria, respectively (Supplementary Table S2). At the level of the concatenated ribosomal proteins dataset, Margulisbacteria showed an almost 2-fold larger within-group mean distance than Cyanobacteria.

We then compared the rates of evolution of CP43 and CP47 with those of Alpha and Beta, using a Bayesian relaxed molecular clock approach with identical calibrations, molecular clock parameters, and a simplified, but highly constrained sequence dataset (see Materials and Methods for an expanded rationale). The goal of these experiment is not to use the clock to estimate divergence times, but to measure and compare the rates of protein evolution that are required to simulate any chosen span of time between the ancestral duplications and the MRCA of Cyanobacteria. We used an autocorrelated log normal model of rate variation with a non-parametric CAT+Γ model of amino acid substitutions to extract rates of evolution. We will refer to the span of time between the duplication points leading to Alpha and Beta (***d*AB**), or to CP43 and CP47 (***d*CP**), and the MRCA of Cyanobacteria as ΔT (schematized in Figure 3).

**Figure 3.**
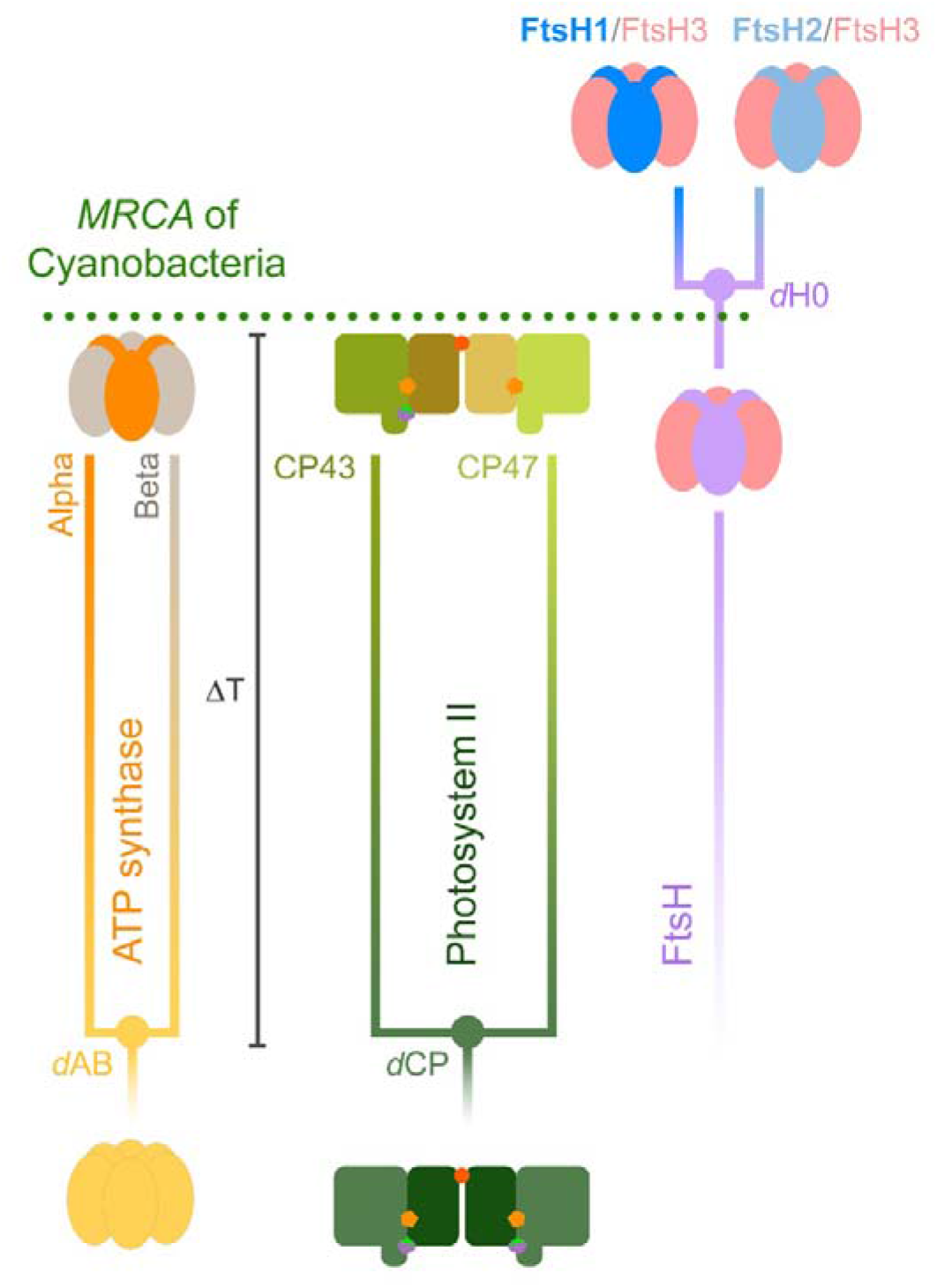
Schematic representation of ancestral duplication events. The MRCA of Cyanobacteria inherited a standard F-type ATP synthase, with a heterohexameric catalytic head (F_1_) made of alternating subunits Alpha and Beta; and a PSII with a heterodimeric core and antenna. ΔT denotes the span of time between the ancestral duplication events and the MRCA of Cyanobacteria, which in the case of ATP synthase, is suspected to antedate the divergence of Bacteria and Archaea and their further diversification. FtsH contains an N-terminal membrane-spanning domain attached to a soluble domain consisting of an AAA (ATPase associated with diverse activities) module attached to a C-terminal protease domain. FtsH is universally conserved in Bacteria, has a hexameric structure like that of ATP synthase’s catalytic head, and can be found usually as homohexamers, but also as heterohexamers. The MRCA of Cyanobacteria likely inherited three variant FtsH subunit forms, one of which appears to have duplicated after the divergence of the genus *Gloeobacter*, and possibly other early-branching Cyanobacteria [40]. This late duplication led to FtsH1 and FtsH2, which form heterohexamers with FtsH3, following the nomenclature of Shao, et al. [40] FtsH1/FtsH3 is found in the cytoplasmic membrane of Cyanobacteria, while FtsH2/FtsH3 is involved in the degradation of PSII and other thylakoid membrane proteins.

In Figure 4a to d we examine the changes in the rate of evolution under specific evolutionary scenarios. In the case of ATP synthase, we first assumed that the MRCA of Cyanobacteria occurred after the GOE to simulate scenarios similar to those presented in [8] or [11], at about 1.7 Ga; and that *d*AB occurred at 3.5 Ga (**Δ T = 1.8 Ga**). Under these conditions the average rate of evolution of Alpha and Beta is calculated to be 0.28 ± 0.06 substitutions per site per Ga (δ Ga^-1^). We will refer to the average rate through the Proterozoic as ν_min_. In this scenario, the rate of evolution at the point of duplication, which we denote ν_max_, is 7.32 ± 1.00 δ Ga^-1^, making ν_max_/ν_min_ 26. In other words, when the span of time between the ancestral pre-LUCA duplication and the MRCA of Cyanobacteria is 1.8 Ga, the rate of evolution at the point of duplication is about 26 times greater than any rate observed through the diversification of Cyanobacteria or photosynthetic eukaryotes. To place these rates in the larger context of protein evolution, we encourage the reader to refer to Supplementary Text S1.

**Figure 4.**
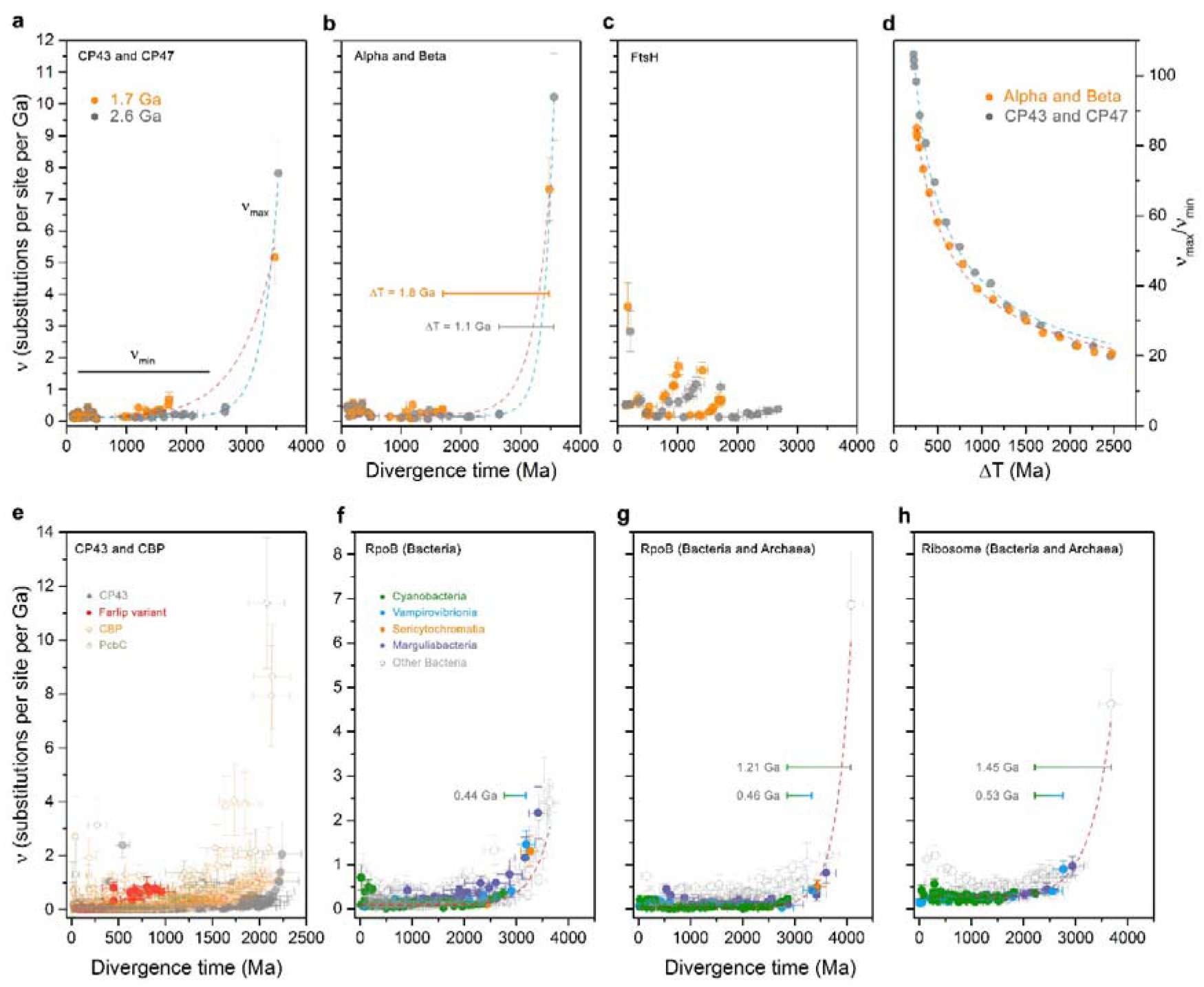
Comparison of the changes in the rates of evolution as a function of time. **a** Rates of CP43 and CP47 modelled using two specific evolutionary scenarios. Orange trace was calculated under the assumption that the MRCA of Cyanobacteria occurred after the GOE, at ~1.7 Ga, while the duplication of CP43 and CP47 occurred at ~3.5 Ga. The fastest rate seen at the point of duplication is denoted as ν_max_, and stabilizes during the Proterozoic, ν_min_. Grey dots represent a scenario in which the MRCA of Cyanobacteria is thought to have occurred before the GOE, at ~2.6 Ga instead. **b** Same calculations as **a** but performed on Alpha and Beta sequences of cyanobacterial and plastid ATP synthase. **c** Rates of evolution of cyanobacterial and plastid FtsH subunits assuming a MRCA of Cyanobacteria at 1.7 and 2.6 Ga. **d** Changes in the rates of evolution with varying ΔT for ATP synthase (orange) and PSII subunits (grey). **e** Changes in the rates of evolution as a function of time for a relaxed molecular clock computed with the full dataset of CP43 (full circles) and CBP sequences (open circles). **f** Changes in the rate of evolution of bacterial RpoB showing an exponential decrease in the rate of evolution. The bar denotes the span of time between the MRCA of the clade containing Vampirovibrionia and Cyanobacteria and the MRCA of Cyanobacteria (0.44 Ga). **g** Changes in the rate of evolution of bacterial and archaeal RpoB. The longer bar represents the span of time between the divergence of Archaea and Bacteria and the MRCA of Cyanobacteria (1.21 Ga). **h** Changes in the rate of evolution of a dataset of concatenated ribosomal proteins showing similar patterns of evolution as RpoB. Error bars on each data point are standard errors on the rate and mean divergence time.

Now, if we consider a scenario in which *d*AB is 4.0 Ga and leaving all other constraints unchanged, ν_max_ is 6.02 ± 0.9 δ Ga^-1^ resulting in a ν_max_/ν_min_ of 21. If instead we keep the duplication at 3.5 Ga but assume that the MRCA of Cyanobacteria occurred before the GOE to simulate a more conservative scenario, at 2.6 Ga (**ΔT = 1.1 Ga**), we obtain that ν_min_ is consequently slower, 0.25 ± 0.06 δ Ga^-1^, when compared to a post-GOE ancestor. This older MRCA (smaller ΔT) thus leads to a rise in ν_max_, calculated to be 10.22 ± 1.37 δ Ga^-1^ and leading to a ν_max_/ν_min_ of 40. Given that the phylogenetic distance is a constant, the rate of evolution increases with a decrease in ΔT following a power law function. We had observed nearly identical evolutionary patterns for the core RC proteins D1 and D2 of PSII [18]. The change in ν_max_/ν_min_ as a function of ΔT is shown in Figure 4d.

The core antenna of PSII, CP43 and CP47, showed patterns of divergence very similar to those of Alpha and Beta, both in terms of phylogenetic distances between paralogues and rates of evolution between orthologues (Figure 4a and b). The average rate of evolution of CP43 and CP47, assuming that the MRCA of Cyanobacteria occurred at 1.7 Ga, and the duplication (*d*CP) at 3.5 Ga (**ΔT = 1.8 Ga**), is 0.19 ± 0.05 δ Ga^-1^. Slower than for Alpha and Beta under the same condition. This slower rate is consistent with the fact that CP43 and CP47 show less sequence divergence between orthologues at all taxonomic ranks of oxygenic phototrophs when compared to Alpha and Beta (see Supplementary Table S3). Furthermore, the rate at *d*CP, ν_max_, was 5.17 ± 0.84 δ Ga^-1^, generating a ν_max_/ν_min_ of 27, similar to Alpha and Beta (Figure 4). Thus, even when ΔT is 1.8 Ga, the rate at duplication point needs to be 27 times greater than the average rates observed during the Proterozoic. If we consider instead that the MRCA of Cyanobacteria occurred at 2.6 Ga and *d*CP at 3.5 Ga **(ΔT = 1.1 Ga**), this would slowdown ν_min_ to 0.16 ± 0.04 δ Ga^-1^, while ν_max_ would increase to 7.81 ± 1.01 δ Ga^-1^ resulting in a ν_max_/ν_min_ of 49. Therefore, the molecular evolution of the core subunits of PSII parallels that of ATP synthase both in terms of rates and distances through geological time.

We then studied a relatively recent gene duplication event (Figure 4c), which occurred long after the LUCA, but also after the MRCA of Cyanobacteria: that leading to Cyanobacteria-specific FtsH1 and FtsH2 (*d*H0) [40]. This more recent duplication served as a point of comparison and control (see Figure 3 for a scheme). In marked contrast to *d*AB, the rate at the point of duplication was 0.66 ± 0.21 δ Ga^-1^. We found that FtsH1 is evolving at an average rate of 1.42 ± 0.29, while FtsH2 at a rate of 0.24 ± 0.06 δ Ga^-1^ under the assumption that MRCA of Cyanobacteria occurred at 1.7 Ga. Thus, under the assessed conditions, FtsH1 is evolving about 5.3 times faster than FtsH2, while the latter is evolving at a rate similar to that of Alpha and Beta. If the MRCA of Cyanobacteria is assumed to have occurred at 2.6 Ga, all rates slowdown respectively, but the rate of FtsH1 remains over five times faster than FtsH2. Unlike *d*AB, *d*H0 is consistent with classical neofunctionalization, in which the copy that gains new function experiences an acceleration of the rate of evolution [41, 42]. Like PSII and ATP synthase, the calculated rates of evolution match observed distances as estimated by the change in the level of sequence identity as a function of time, in which the fastest evolving FtsH1 accumulated greater sequence change than FtsH2 in the same period (Supplementary Table S3).

Given that the complex evolution of CP43 and CBP involved several major duplication events and potentially large variations in the rate of evolution (Figure 1 and Supplementary Figure S1), we carried out a molecular clock analysis of a large dataset of 392 CP43 and 465 CBP proteins using cross-calibrations across paralogues. Clocks were executed with no constraint on the MRCA of Cyanobacteria. The estimated mean divergence time for the oldest node, the duplication at the origin of CBP, is 2.23 Ga (95% confidence interval, CI: 1.90 - 2.69 Ga) using an autocorrelated log normal model (see Figure 4e, Supplementary Figure S6 for a chronogram, and Supplementary Table S4 for a comparison of estimated ages under different models). The mean divergence time for the node representing the CP43 inherited by the MRCA of Cyanobacteria was calculated to be 2.22 Ga (95% CI: 1.88 - 2.68 Ga). Thus, a span of time of only 15 Ma is measured between these two mean ages. The average rate of evolution of CP43, not including CBP sequences, was found to be 0.14 ± 0.05 δ Ga^-1^, which is in the same range as determined in the simplified, but highly constrained experiment above. We noted a 6-fold increase in the rate of evolution associated with the duplication leading to the farlip-CP43 variant (Figure 4e). This duplication led to an acceleration of the rate similar in magnitude to that of FtsH1/FtsH2 and is consistent with a neofunctionalization as the photosystems evolved to use far-red light and bind chlorophyll *f*.

CBP sequences, on average, display rates of evolution about three times faster than CP43 (Figure 4e). However, the serial duplications that led to the evolution of CP43-derived light harvesting complexes resulted in accelerations in the rate of evolution of a similar magnitude as observed for *d*AB and *d*CP. The largest of these is associated with the origin of PcbC [30], a variant commonly found in heterocystous Cyanobacteria and Cyanobacteria that use alternative pigments, such as chlorophyll *b, d* and *f*. The ancestral node of PcbC was timed at 2.07 Ga (95% CI: 1.76 - 2.50 Ga) with a rate of 11.7 ± 2.42 δ Ga^-1^, decelerating quickly, but stabilizing at about four times faster rates than the average rate of CP43. We find it noteworthy that the fast rates of evolution associated with the origin of CBP are not associated with very large spans of time between these and CP43, nor did it result in very old root node ages despite the use of very broad constraints.

### 2.3. Species divergence

To understand the evolution of MSV relative to Cyanobacteria we wished to apply a molecular clock to a system where the calculated rates could be compared to observed rates as determined by distances between species of known divergence times or at similar taxonomic ranks. We found RpoB to be suitable for this because it has been inherited vertically with few instances of horizontal gene transfer and had enough signal to resolve known phylogenetic relationships between and within clades. In collecting the RpoB sequences, we noted for the first time that Margulisbacteria and Vampirovibrionia share a comparatively greater level of divergence at similar taxonomic ranks than Cyanobacteria. For example, the level of sequence divergence of RpoB from two species of *Termititenax* (Margulisbacteria) [43] is about 40% greater than the distance between *Gloeobacter* spp. and any other cyanobacterium (not including gaps and insertions), the latter being the largest distance between oxygenic phototrophs. In the case of Gastranaerophilales (Vampirovibrionia) [5], which are specialised gut bacteria and should therefore not be much older than animals, the level of sequence identity of RpoB was found to be 70% for the two most distant strains in this group, contrasted to 84% for *Gloeobacter* spp. when compared to any other cyanobacterium. As listed in Supplementary Table S2, within-group mean distances suggest that faster rates are widespread and not just unique to RpoB.

We implemented a set of 12 calibrations across bacteria, including two calibrations on Margulisbacteria and two in Vampirovibrionia with the aim of covering both slower and faster evolving lineages. The following results are based on an autocorrelated log normal molecular clock using CAT+Γ, a root constrained with a broad interval ranging from between 4.52 and 3.41 Ga, and as described in Materials and Methods (Figure 5). We found this to perform well and provided results comparable to other independent studies that did not combine a full set of MVS sequences and other clades with phototrophs in a single tree (Table 1). Nonetheless, a pipeline of sensitivity experiments tested the dependency of these results on models and prior assumptions: these are shown and described in Supplementary Figure S7 and S8.

**Figure 5.**
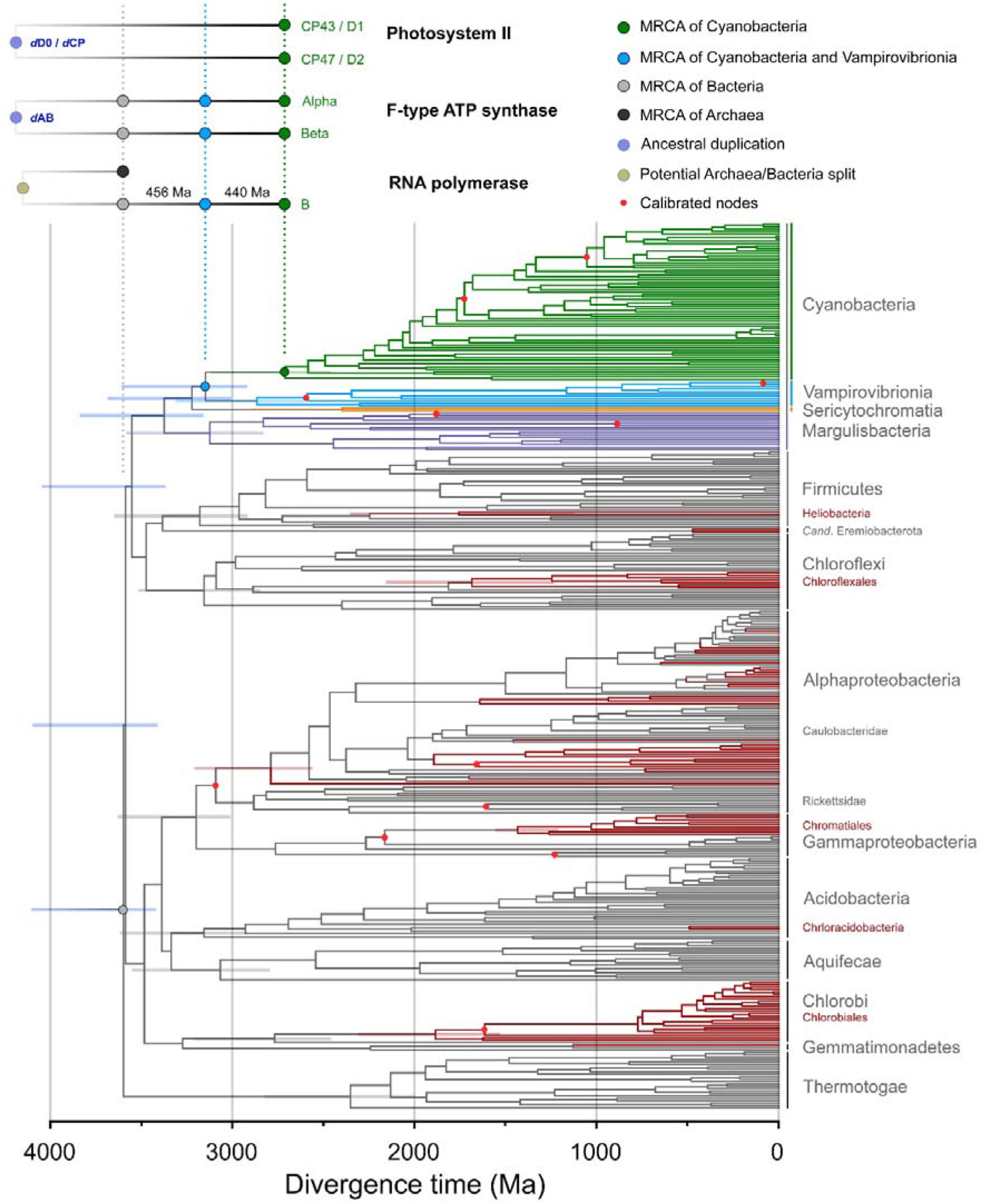
Bayesian relaxed molecular clock of bacterial RpoB. The tree highlights the spans of time between the MRCA of Cyanobacteria (green dot) and their closest relatives. Calibrated nodes are marked with red dots. Anoxygenic phototrophs are highlighted in red branches and non-phototrophic bacteria in grey, with the exception of Margulisbacteria, Sericytochromatia and Vampirovibronia, which are coloured as indicated in the figure. Superimposed at the top are the implied distribution and divergence time for ATP synthase and PSII. Horizontal bars within the tree mark 95% confidence intervals. These are shown in selected nodes of interest for clarity but see Table 1.

**Table 1.** Divergence time estimates

| Event | RpoB (Ga)<br>(95% CI) | Concatenated<br>ribosomal<br>proteins | TimeTree<br>compilation<sup>a</sup> | Shih et al.<sup>b</sup> | Magnabosco et<br>al.<sup>c</sup> |
| --- | --- | --- | --- | --- | --- |
| Divergence of<br>Thermotogae | 3.64<br>(3.43 - 4.11) | 3.01<br>(2.64 - 3.51) | 3.81<br>(3.51 - 4.16) |  |  |
| Divergence of<br>Margulisbacteria | 3.42<br>(3.17 - 3.86) | 2.93<br>(2.58 - 3.44) |  |  |  |
| Divergence of<br>Sericytochromatia | 3.26<br>(3.01 - 3.68) |  |  |  |  |
| Divergence of<br>Vampirovibrionia/Stem<br>Cyanobacteria | 3.19<br>(2.92 - 3.61) | 2.76<br>(2.40 - 3.23) | 3.19<br>(2.26 - 3.55) | 2.54<br>(2.09 - 3.06) | 2.77<br>(2.39 - 2.93) |
| MRCA of<br>Cyanobacteria | 2.75<br>(2.46 - 3.13) | 2.22<br>(1.86 - 2.64) | 2.24<br>(1.81 - 2.82) | 2.02<br>(1.72 - 2.37) | 2.24<br>(1.91 - 2.41) |
| Divergence of<br>Heliobacteria | 2.28<br>(1.74 - 2.78) |  | 2.23<br>(1.99 - 2.48) |  |  |
| MRCA of<br>Heliobacteria | 1.78<br>(1.13 - 2.36) |  |  |  |  |
| Divergence of<br>phototrophic<br>Chloroflexi | 1.83<br>(1.41 - 2.25) |  | 1076 | 1.098<br>(0.80 - 1.41) | 2.86<br>(2.49 - 3.00) |
| MRCA of phototrophic<br>Chloroflexi | 1.71<br>(1.24 - 2.16) |  |  | 0.86<br>(0.61 - 1.14) | 1.98<br>(1.68 - 2.43) |
| Divergence of<br>phototrophic Chlorobi | 2.80<br>(2.45 - 3.20) |  |  |  | 2.56<br>(2.26 - 2.85) |
| MRCA of phototrophic<br>Chlorobi | 1.91<br>(1.53 - 2.31) |  | 525 |  | 1.71<br>(1.64 - 1.95) |
| Earliest phototrophic<br>Proteobacteria | 2.82<br>(2.55 - 3.21) |  | 2.48<br>(2.04 - 2.62) |  |  |
| Divergence of<br>*Chloracidobacterium* | 2.04<br>(1.53-2.51) |  |  |  |  |

<sup>a</sup>Data from TimeTree compiling estimated divergence times from independent studies as described in ref. [44] For the divergence of Thermotogae the estimated age was taken from eight different studies (n=8). For the divergence of Melaibacteria (or divergence from Cyanobacteria's closest relatives), n=10. For stem Heliobacteria, n=2. For stem phototrophic Chloroflexi and Chlorobi, the values were reported from a single study, Marin, et al. [45] For earliest potential phototrophic Proteobacteria, n=3. This latter was taken as the node made by the clades including *Niveispirillum lacus* and *Rhodobacter sphaeroides* to match our RpoB tree. The independent studies used by TimeTree to generate the estimated ages are listed in Supplementary Table S8.
<sup>b</sup>Data taken from ref. [8] Model T68 for the cyanobacterial dates. That is, no GOE calibration with a calibration on *Bangiomorpha*. The dates on Chloroflexi were taken from ref. [46]
<sup>c</sup>Data taken from ref. [9] Model A. This used a 1.2 Ga calibration on heterocystous Cyanobacteria.

The root of the tree (divergence of Thermotogae) was timed at 3.64 Ga (95% CI: 3.42 - 4.11 Ga) and the divergence of Cyanobacteria at 2.74 Ga (95% CI: 2.46 - 3.12 Ga). Thus, the span of time between the mean age of the root and the mean age of the MRCA of Cyanobacteria was calculated to be 0.89 Ga. The span of time between Margulisbacteria and Cyanobacteria was found to be 0.67 Ga; and between Vampirovibrionia and Cyanobacteria 0.44 Ga (Figure 4f and 5). The latter is a value that is consistent with previous studies using entirely different rationales, datasets and calibrations [8, 9].

We also noted an exponential decrease in the rates of evolution of RpoB through the Archean, which stabilised at current levels in the Proterozoic (Figure 4f). The rate at the root node was calculated to be 2.37 ± 0.45 δ Ga^-1^ and the average rate of evolution of RpoB during the Proterozoic was found to be 0.19 ± 0.06 δ Ga^-1^. The average rate of cyanobacterial RpoB was 0.14 ± 0.04 δ Ga^-1^; for Margulisbacteria was 0.44 ± 0.17 δ Ga^-1^, and for Vampirovibrionia 0.19 ± 0.05 δ Ga^-1^: about 3.1x and 1.3x the mean cyanobacterial rate respectively. These rates agree reasonably well with the observed distances (Supplementary Table S2), further indicating that the calibrations used in these clades performed well and recapitulated patterns of evolution consistent with lifestyle and trophic modes. Nevertheless, we suspect that the values for MSV represent underestimations of the true rates of evolution (slower than they should be), as some of the clades that include symbionts still appear much older than anticipated from their hosts (Supplementary Text S2).

Furthermore, a more complex, but commonly used model like CAT+GTR+Γ implementing a birth-death prior with ‘soft bounds’ on the calibrations, resulted in rates that were smoothed out, which translated into spread-out divergence times with Margulisbacteria and Vampirovibrionia evolving at 1.9x and 0.7x times the cyanobacterial rates, respectively (Supplementary Figure S8, model **n** to **p**). These weird rate effects are thus translated into a Mesoproterozoic, very late, age for Cyanobacteria, and a relatively older divergence time for Vampirovibrionia: results that replicate those presented recently in ref. [11].

To investigate the effect of the age of the root on the divergence time of MVS and Cyanobacteria we also varied the root prior from 3.2 to 4.4 Ga (Supplementary Figure S7). We noted that regardless of the time of origin of Bacteria (approximated by the divergence of Thermotogae in our analysis), a substantially faster rate is required during the earliest diversification events, decreasing through the Archaean and stabilizing in the Proterozoic. This matches well the patterns of evolution of ATP synthase and PSII core subunits as shown in the previous section.

We then compared the above RpoB molecular clock with a different clock that included a set of 112 diverse sequences from Archaea, in addition to the sequences from Thermotogae, MSV, and Cyanobacteria, but removing all other bacterial phyla (Figure 6a). We found that the calculated average rate of evolution of bacterial RpoB during the Proterozoic was slower (0.09 ± 0.03 δ Ga^-1^) than in the absence of archaeal sequences (0.19 ± 0.06 δ Ga^-1^), resulting in overall older mean ages (see Figure 4g). However, the rate at the Bacteria/Archaea divergence point was 6.87 ± 1.17 δ Ga^-1^, similar to the rate for *d*AB and *d*CP, requiring therefore an exponential decrease in the rates similar to that observed for ATP synthase and the PSII core subunits. Notably, this rate and exponential decrease is associated with a span of time between the mean estimated ages of the LUCA and the MRCA of Cyanobacteria of 1.21 Ga. The span between Vampirovibrionia and Cyanobacteria was found to be 0.46 Ga (Figure 6a).

**Figure 6.**
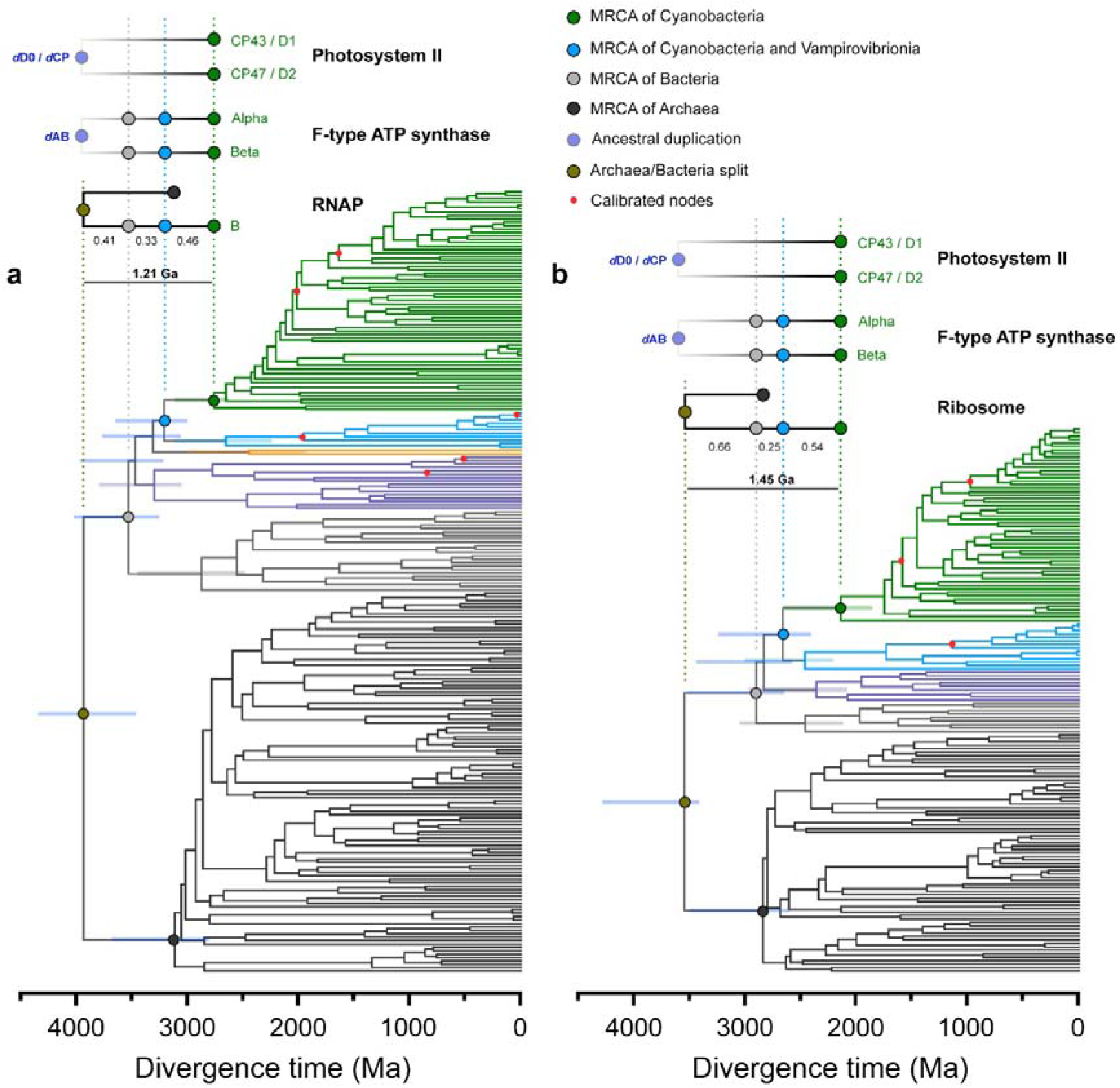
Comparison of relaxed molecular clocks of RpoB (**a**) and a concatenated dataset of ribosomal proteins (**b**) that include sequences from Archaea and as described in the main text. RNAP stands for RNA polymerase. Clocks were calculated using a calibration on the root with a maximum of 4.52 Ga and a minimum of 3.41 Ga and applying a log normal autocorrelated clock with CAT+Γ model. Bars on selected nodes denote 95% confidence intervals.

Figure 5 also highlights that the MRCA of none of the groups containing anoxygenic phototrophs nor their divergence from their closest non-phototrophic relatives appears to be older than one of the oldest and best accepted geochemical evidence for photosynthesis at 3.41 Ga [47] (Table 1). For example, the MRCA of Heliobacteria or of phototrophic Chlorobi, containing homodimeric type I RCs, which have traditionally been described as to harbour primitive forms of photosynthesis, are likely to have existed after the GOE, even allowing for large uncertainties in the calculations.

Finally, we performed a similar molecular clock analysis on a subset of concatenated ribosomal proteins published by Hug et al. [38] that included Archaea, Thermotagae, Margulisbacteria, Vampirovibrionia and Cyanobacteria. Even though this dataset generated younger ages compared to RpoB (see Table 1 and Figure 6), a similar exponential decrease in the rates was observed with a rate at the Bacteria/Archaea divergence point measured at 4.62 ± 0.71 δ Ga^-1^ and an average rate of bacterial ribosomal protein evolution of 0.30 ± 0.07 δ Ga^-1^ (Figure 4h). The span of time between the mean estimated ages of the LUCA and the MRCA of Cyanobacteria was 1.45 Ga; and between Vampirovibrionia and Cyanobacteria 0.54 Ga (Figure 6b). If one assumes therefore that the span of time between the LUCA and the diversification of Bacteria is narrower, that would imply a faster rate at the root node and a steeper deceleration of the rates.

### 2.4. Structural analysis

A fundamental premise of our investigation is that water oxidation started before the duplication of D1 and D2, and CP43 and CP47. The rationale behind this premise has been laid out before [17, 48], and more extensively recently [18]. This rationale raises the question of how the D2/CP47 side of the RC lost its capacity to carry out water-splitting catalysis. To gain further insight on the nature of the structural site around the water-oxidising complex in the ancestral photosystem, we used ancestral sequence reconstruction to predict the most probable ancestral states. We will refer to the ancestral protein to D1 and D2 as D0 (Supplementary Figure S9). We generated 14 predicted D0 sequences using a combination of three ASR methods and amino acid substitution models. On average the 14 D0 sequences had 87.12 ± 0.55% sequence identity indicating that the different algorithms provided largely consistent results. While the regions that include all transmembrane helices are aligned unambiguously, the N-terminal and C-terminal ends were aligned less confidently due to greater sequence variability at both ends. Nonetheless, we found that the predicted D0 sequences retain more identity with D1 than D2 along the entire sequence. The level of sequence identity of D0 compared to the D1 (PsbA1) of *Thermosynechococcus vulcanus* was found to be 69.58 ± 0.55%, and 36.32 ± 0.15% compared to D2.

The ligands to the Mn_4_CaO_5_ in PSII are provided from three different structural domains (Supplementary Figure S10): 1) the D1 ligands D170 and E189, located near the redox Y_Z_. These are in the lumenal loop between the 3^rd^ and 4^th^ transmembrane helices. 2) The D1 ligands H332, E333, D342, and A344 located at the C-terminus. 3) The CP43 ligands, E354 and R357, located in the extrinsic loop between the 5^th^ and 6^th^ helices, with the latter residue less than 4 Å from the Ca. Remarkably, there is structural and sequence evidence supporting the loss of ligands in these three different regions of CP47/D2.

In all the D0 sequences, at position D1-170 and 189, located in the unambiguously aligned region, the calculated most likely ancestral states were E170 and E189, respectively. The mutation D170E results in a PSII phenotype with activity similar to that of the wild-type [49]. At position D1-170, a glutamate was predicted with average posterior probabilities (PPs) of 44.2% (FastML), 67.2% (MEGA) and 77.0% (PAML). At position D1-E189, the average PPs for glutamate were 31.1% (FastML), 35.2% (MEGA) and 40.2% (PAML). The distribution of PPs across a selection of D0 sequences and key sites is presented in Supplementary Figure S11 and Table S5. In contrast, D2 has strictly conserved phenylalanine residues at these positions, but the PP of phenylalanine being found at either of these positions was less than about 5% for all predicted D0 sequences. As a comparison, the redox active tyrosine residues Y_Z_ (D1-Y161) and Y_D_ (D2-Y160), which are strictly conserved between D1 and D2, have a predicted average PPs of 68.8% (FastML), 98.8% (MEGA) and 98.6% (PAML). Therefore, the ligands to the catalytic site in the ancestral protein leading to D2 were likely lost by direct substitutions to phenylalanine residues, while retaining the redox active D2-Y160 (Y_D_) and H189 pair (Supplementary Figure S10).

Prompted by the finding of a Ca-binding site at the electron donor site of the homodimeric type I RC of Heliobacteria (Firmicutes) with several similarities to the Mn_4_CaO_5_ cluster of PSII, including a link to the antenna domain and the C-terminus [19], we revisited the sequences and structural overlaps of CP43 and CP47. We found that a previously unnoticed structural rearrangement within the extrinsic loop occurred in one subunit relative to the other (marked EF3 and EF4 in Figure 7, Supplementary Figure S12 and S13). CP43 retains the simplest domain, being about 60 residues shorter than CP47. If CP43 retains the ancestral fold, the additional sequence in CP47’s swapped domain (EF4 in Figure 7d) would have contributed to the loss a catalytic cluster as it inserted one phenylalanine residue (CP47-F360) into the electron donor site, less than 4Å from Y_D_. An equivalent residue does not exist in CP43.

**Figure 7.**
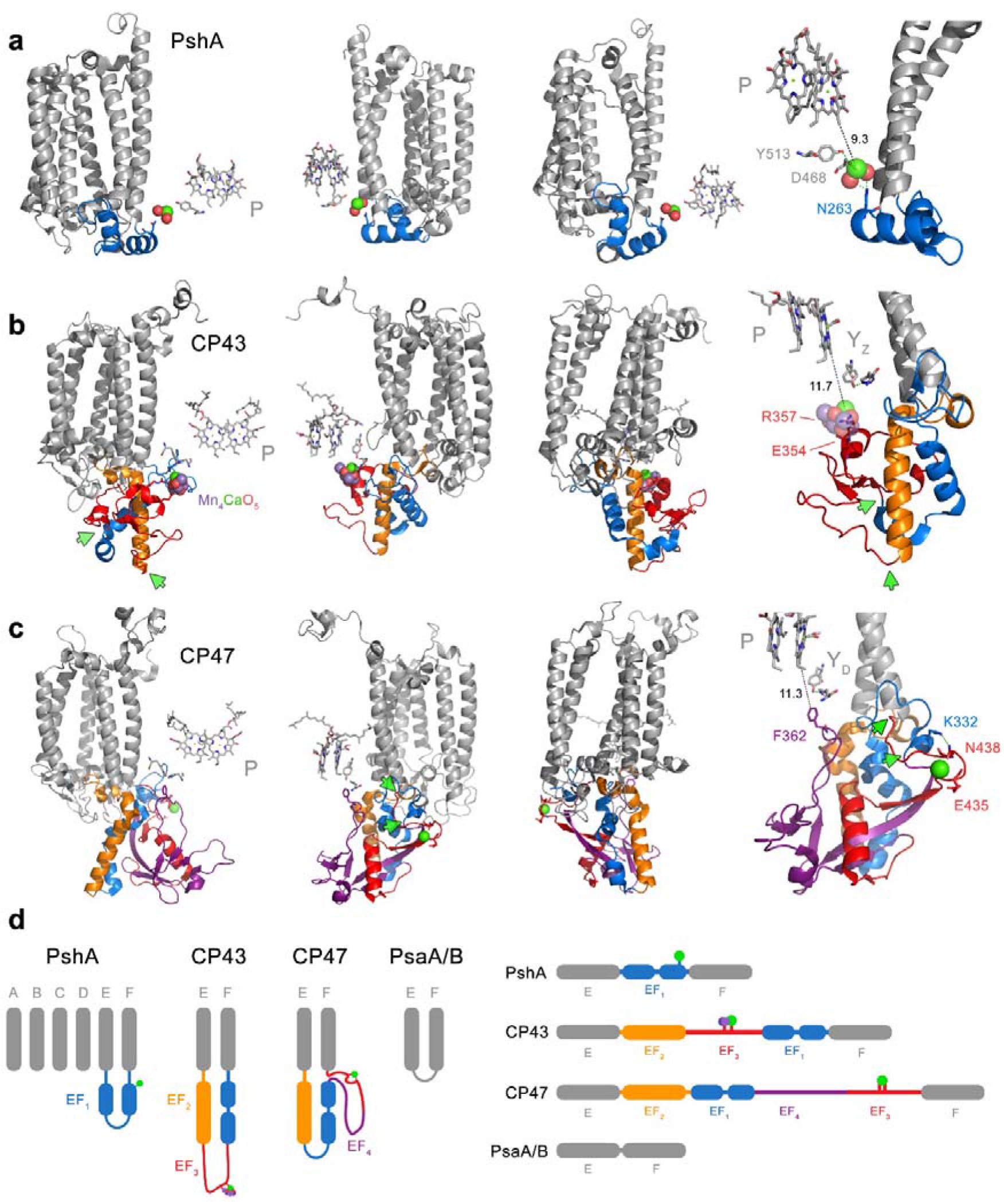
Structural rearrangements of the large extrinsic loop of CP43 and CP47. **a** The antenna domain of heliobacterial PshA is shown in three different rotated perspectives. Only the first six transmembrane helices of the antenna are shown for clarity. A Ca at the electron donor side is bound from an extrinsic loop between the 5^th^ (E) and 6^th^ (F) helices. This extrinsic loop, EF_1_ (blue), is made of two small alpha helices. The fourth molecular view furthest to the right shows the link between the electron donor site and EF_1_ in closer detail. **b** The CP43 subunit of PSII with the extrinsic loop shown in colours. **c** The CP47 subunit of PSII. Immediately after the 5^th^ helix (E), a long alpha helix protrudes outside the membrane in both CP43 and CP47 and showing structural and sequence identity (orange). We denote this helix EF_2_. After EF_2_ structural differences are noticed between CP43 and CP47 as schematised in panel **d**. In CP43, after helix EF_2_ a loop is found (shown in red ribbons), which we denote EF_3_. This contains the residues that bind the Mn_4_CaO_5_ cluster and it is followed by a domain that resembles EF_1_ in the HbRC at a structural level. In CP47, EF_3_ and EF_1_ retain sequence identity with the respective regions in CP43. CP47 has additional sequence that is not found in CP43 (EF_4_, purple). The green arrows mark the position at which the domain swap occurred in CP43 relative to CP47. We found that the CP43-E354 and R357 are found in the equivalent domain in CP47 as E436 and N438 coordinating a Ca atom. N438 (EF_3_) links to EF_1_ via K332. It is unclear if the EF_1_ region in the HbRC is strictly homologous to that in CP43 and CP47 as very little sequence identity is found between the two: however, a couple of conserved residues between all EF_1_ may suggest it emerged from structural domains present in the ancestral RC protein (see Supplementary Figure S13).

We then noted that in the swapped region (EF_3_ in Figure 7d), sequence identity is retained between CP43 and CP47 (Supplementary Figure S12). We found that CP43-E354 and R357 are equivalent to CP47-E435 and N438. An inspection of the crystal structure of cyanobacterial PSII showed that these two residues specifically bind a Ca of unknown function in (Figure 7c and d). The presence of an equivalent glutamate to CP43-E354 in CP47 is consistent with this being already present before duplication.

Finally, a peculiar but well-known trait conserved across Cyanobacteria and photosynthetic eukaryotes is that the 5’ end of the *psbC* gene (CP43) overlaps with the 3’ end of the *psbD* gene (D2) usually over 16 bp (Supplementary Table S6). The *psbD* gene contains a well-defined Shine-Dalgarno ribosomal binding site downstream of the *psbC* gene and over the coded D2 C-terminal sequence [50–52]. The evolution of this unique gene overlap has no current explanation in the literature, but its origin could have disrupted the C-terminal ligands of the ancestral protein leading to the modern D2, an event that would have contributed to and favoured heterodimerization.

## 3. Discussion

### 3.1. Origin of oxygenic photosynthesis and rates of evolution

The duplication of ATP synthase’s ancestral catalytic subunit, and the archaeal/bacterial divergence of RNA polymerase and the ribosome, are some of the oldest evolutionary events known in biology [20–28]. When taken on their own, their phylogenetic features are often interpreted as evidence of the earliest origin. We have shown in here and in our previous work [18] that PSII show patterns of molecular evolution that closely parallel those of these very ancient systems and independently of the exact time of origin of Cyanobacteria or their closest relatives. Therefore, it cannot be taken for granted that there was ever a long period of time between the origin of life and the origin of anoxygenic photosynthesis, followed by another long period of time between the origin of anoxygenic photosynthesis and the origin of oxygenic photosynthesis.

The exponential decrease in the rates of evolution observed in the studied systems, even when the span of time between the ancestral duplications (or the LUCA) and the MRCA of Cyanobacteria was well over a billion years, are consistent with our assessment that a large ΔT for the evolution of the core subunits of ATP synthase and PSII cannot be explained entirely by duplication-driven effects. Instead, we speculate that these effects are the result of the faster rates of evolution that are expected to have occurred during the earliest history of life due to unrestricted positive selection at the origin of bioenergetics, accelerated rates of mutations caused by environmental factors [53–55], lack of sophisticated DNA repair mechanisms [56], or other genetic properties attributed to early life [57, 58], yet in combination with very large spans of time.

It is worth highlighting that ATP synthase, RNA polymerase, the ribosome, and the photosystems, are all complex molecular machines, with crucial functions and under strict regulation. These features largely explain their slow rates of evolution and the high degree of sequence (structural and functional) conservation through geological time. A similar level of complexity, however, can be traced back to the last common ancestor of each one of these molecular systems regardless of how ancient they truly are. Now, given that the rates of evolution of the core of PSII have remained slower than those of ATP synthase for billions of years, even through evolutionary processes that could be associated with acceleration in the rates of evolution such as: 1) primary endosymbiosis at the origin of photosynthetic eukaryotes [59], 2) radiations within clades of flowering plants [60], or 3) the radiation of marine *Prochlorococcus* and *Synechococcus* [61, 62], just to name a few; if those rates have remained proportional since the origin of the enzymes, it could imply that the duplications of the core of PSII occurred before the duplication leading to Alpha and Beta.

In consequence, to argue that oxygenic photosynthesis is a late evolutionary innovation relative to the origin of life, one must first demonstrate that the rates of protein evolution of PSII and ATP synthase catalytic core subunits, RNA polymerase, and the ribosome, have not remained proportional to each other through geological time. In such a way that PSII—exclusively—experienced unprecedentedly faster rates of evolution. One could potentially argue that the origin of water oxidation itself could account for this hypothetical unprecedentedly faster rates. However, because of the relationships between *speed, distance,* and *time,* a decrease in ΔT of about 60% (from 1.1 to 0.46 Ga for example), would result in a 30-fold additional increase in the rates at the earliest stages of diversification, resulting in photosystems that would be evolving at rates that surpass those of the fastest evolving peptide toxins (Figure 4d and Supplementary Text S1). It should be noted, that a ΔT of over a billion years already accounts for a period of fast evolution at the origin of all these ancient systems.

The reader should also be reminded that these patterns of evolution, though they might appear somewhat unusual, do not emerge from the application of any particular molecular clock approach or computational analysis, but from the inherently long phylogenetic distance that separates Alpha and Beta, Archaea and Bacteria, or the core subunits of PSII, together with relatively slow rates of evolution throughout their entire multibillion-year diversification process.

### 3.2. Diversification of Bacteria

It has been postulated before that the diversification of the major phyla of Bacteria occurred very rapidly, starting at about 3.4 Ga and peaking at about 3.2 Ga, in what was dubbed as the Archean Genetic Expansion [63]. Another recent independent molecular clock analysis put the MRCA of Bacteria at about 3.4 Ga [11]; and Marin et al. [45] placed the major bacterial radiation, except for the divergence of Thermotogae and Aquifecae, starting at a mean divergence time of about 3.2 Ga too. Our clocks of RpoB or ribosomal proteins are also in agreement with these patterns. A recent study suggested that the major groups of Bacteria and Archaea diversified rapidly after the LUCA and hypothesized that the long distance between archaeal and bacterial ribosomal proteins could be attributed to fast rates of ribosome evolution during their divergence [64], but see also [65]. Our analyses suggest that the more explosive the diversification of prokaryotes, the greater the chance that photosynthetic water-splitting is an ancestral trait of all life. And yet, if phototrophic communities—whether oxygenic or not—already existed 3.2 to 3.4 Ga ago, as it is supported by the geochemical record [47, 66], then the earliest bacteria were likely photosynthetic. That the earliest bacteria were photosynthetic is entirely consistent with the evolution of RC proteins, which indicates that the structural and functional specialization that led to the two photosystem types, antedated the diversification processes leading to the known phyla containing photosynthetic bacteria [34]. This is also consistent with the fact that none of the groups of extant photosynthetic bacteria appear to be older than the earliest geochemical evidence for photosynthesis.

The presented data is also consistent with recent studies of the oxygenation of the planet, which suggests that even if photosynthetic O2 evolution started as early as the oldest rocks, the properties of early Earth biogeochemistry would have maintained very low concentrations of O2 over the Archean without the need to invoke any particular biological innovation or trigger to coincide with the GOE [67–69].

### 3.2. Structural constraints

We showed that the phylogenetic distance between CP43/CP47 and other type I RC proteins is the second largest distance after that between type I and type II (Supplementary Figure S2). It is conventionally considered that the first six transmembrane helices of the photosystems make up the antenna domain, while the photochemical core encompasses the last five helices. Structurally and functionally the antenna domain extends to the 8^th^ helix, both in type I RCs and in PSII; with the latter retaining one antenna chlorophyll in the equivalent 8^th^ helix (marked Z in Figure 8), as well as substantial sequence identity around this chlorophyll’s binding site as shown before [34, 70]. These conserved features indicate that there is not a moment in time during the evolution of PSII in which it was devoid of core antenna proteins. In other words, CP43/CP47 are a descendant of the ancestral core antenna of type II RCs, now lost and replaced in anoxygenic type II RCs of phototrophic Proteobacteria, Chloroflexi and others. Both structural and phylogenetic evidence are in agreement and indicate that the ancestral type II RC, at the dawn of photosynthesis, was architecturally like water-splitting PSII. The above disproves and supersedes a previous hypothesis by Cardona [35] where the antenna of PSII was claimed to have originated from a refolding of an entire type I RC protein.

**Figure 8.**
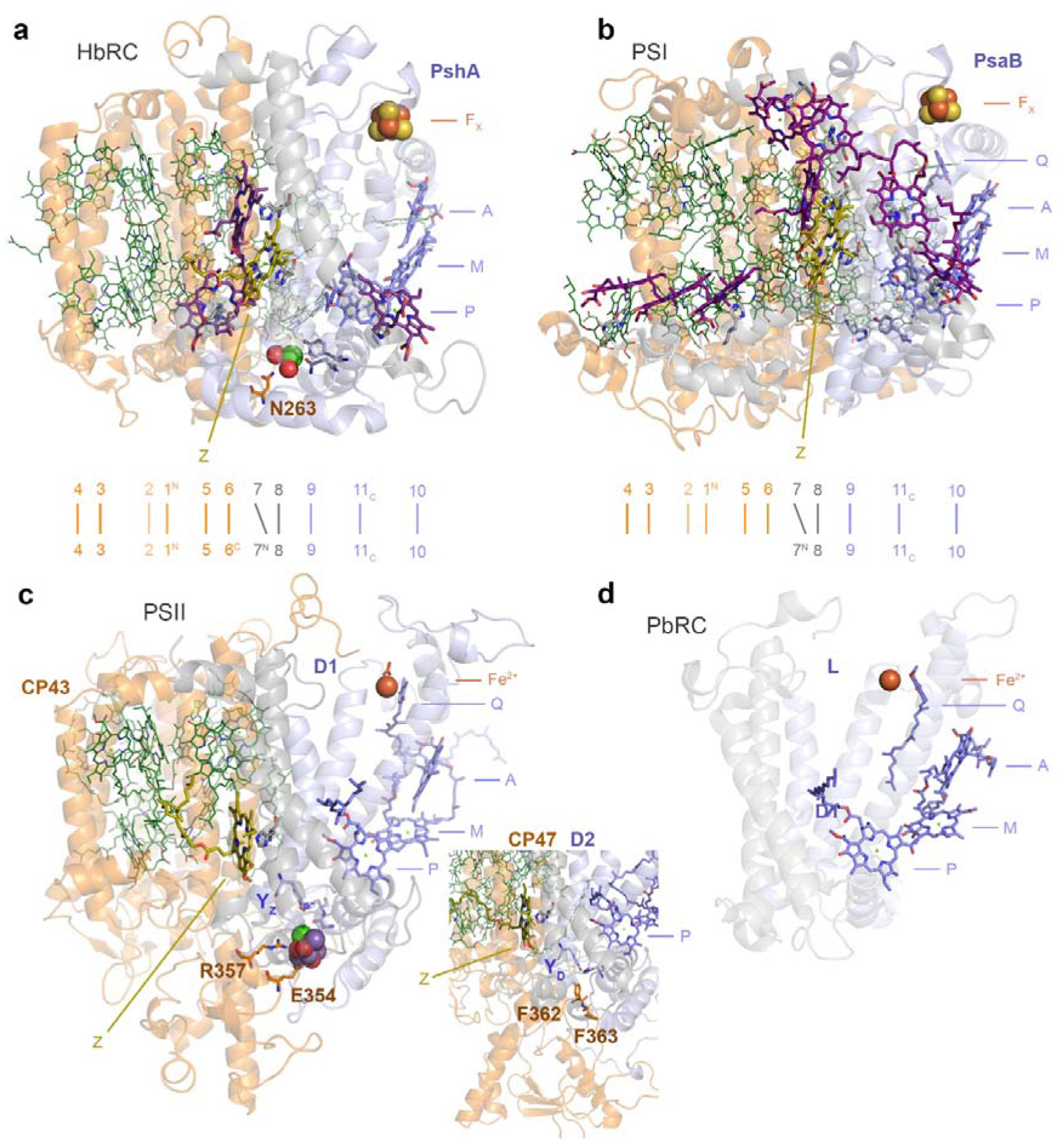
Structural comparisons of the antenna and core domains of photosynthetic RCs. **a** A monomeric unit (PshA) of the homodimeric RC of Heliobacteria (HbRC, Firmicutes). The RC protein is made up of 11 transmembrane helices. The first six N-terminal helices are traditionally considered as the antenna domain (orange ribbons), while the last five helices are the RC core domain (grey and light-blue ribbons). Below the structure, the organization of the 11 helices is laid down linearly for guidance: 1^N^ denotes the first N-terminal helix and 11_C_ the last C-terminal helix. P denotes the “special pair” pigment; M, the “monomeric” bacteriochlorophyll electron donor; and A, the primary electron acceptor. F_X_ is the Fe_4_S_4_ cluster characteristic of type I RCs. Antenna pigments bound by the antenna domain are shown in green lines. Antenna pigments bound by the 7^th^ and 8^th^ helices are shown as purple sticks, except for the bacteriochlorophyll molecule denoted as Z, in olive green and bound by the 8^th^ helix. The antenna domain connects the electron donor side of the RC via N263 and a Ca-binding site. **b** A monomeric unit of PSI (PsaB, Cyanobacteria) following a similar structural organization and nomenclature as in panel **a**. Unlike the HbRC, PSI binds quinones (Q) as intermediary electron acceptors between A and F_X_. **c** A monomeric unit of PSII (CP43/D1 and CP47/D2 in the inset). It has a structural organization similar to that of type I RCs. However, the monomeric unit is split into two proteins after the 6^th^ transmembrane helix. The Mn_4_CaO_5_ cluster is coordinated by D1, but is directly connected to the antenna domain via E354 and R357 in manner that resembles the HbRC. **d** A monomeric unit of an anoxygenic type II reaction (PbRC, Proteobacteria). Unlike type I RCs, type II RCs lack F_X_. Instead, a non-heme Fe^2+^ is found linking the RC proteins. The PbRC lacks antenna domain and the antenna pigment Z bound at the equivalent 8^th^ helix.

Sequence reconstruction of the ancestral subunit to D1 and D2 is consistent with the existence of a highly oxidizing homodimeric photosystem, though transient and short-lived [18], that could split water in either side of the RC [48]. The structural comparisons between CP43/D1 and CP47/D2 suggest a mechanism for heterodimerization and loss of catalysis on one side that accounts for all ligands. These include direct amino acid substitutions, the domain swap within the extrinsic region of the ancestral protein to CP47, and the gene overlap between *psbC* (CP43) and *psbD* (D2). It would be difficult to reconcile these unique structural and genetic features with a scenario in which PSII evolved water oxidation at a late stage, or starting as a purple anoxygenic type II RC, once the heterodimerization process was well underway or completed, or in the absence of core antenna domains, given that these interact directly with the electron donor side of PSII, in a manner strikingly similar to that of the homodimeric type I RC of the Firmicutes [19]. What is more, these structural and functional features also suggest that the atypical forms of D1, lacking ligands to the Mn4CaO5 cluster, such as the so-called chlorophyll *f* synthase [32, 71], or the so-called “rogue” or “sentinel” D1 [31, 72], diverged from forms of D1 that were able to support water oxidation. Even if they originated from early duplications that could have antedated the MRCA of Cyanobacteria [2] and as additionally supported by ASR analysis (Supplementary Figure S9 and S10).

We have calculated that (oxygenic) PSII has experienced the slowest rates of evolution between type II RCs, with the core of the anoxygenic type II RC of Proteobacteria and Chloroflexi evolving approximately five times faster than the core of PSII [18]. That considerably faster rate has led to conspicuous structural changes of the anoxygenic RC relative to PSII and type I photosystems. The consequence of these differences in the rates are visually apparent (Figure 8 and Supplementary Figure S3) as the anoxygenic RC has experienced greater sequence and structural change than PSII. That is, PSII retains a greater number of ancestral traits that can be traced to before the ancestral core duplications. For example, like type I, PSII retains: 1) substantially greater structural symmetry at the core, 2) the antenna functionally linked to the core and the electron donor side, 3) lack of histidine ligands to the monomeric photochemical chlorophylls (marked M in Figure 8), or 4) the use of a chlorophyll *a* derived pigment as the primary electron acceptor. This list is not meant to be exhaustive, but see refs. [19, 73] for additional detail. It follows then, that because these conserved traits can be traced to the common ancestor of type I and type II, then the rates of evolution of PSII should have remained slow and constrained relative to that of other photosystems, since before the core duplications. These structural constraints demonstrate that the core subunits of PSII did not experience unprecedentedly faster rates of evolution. In fact, and not surprisingly, the anoxygenic type II RC which experienced the highest rates of evolution, has accumulated greater change and greater loss of ancestral traits.

## 4. Conclusion

Both phylogenetic and structural evidence converge towards a scenario in which photosynthetic water-splitting started at an early stage during the evolution of life and long before the rise of what we understand today as Cyanobacteria. Nonetheless, greater resolution of divergence time estimation in deep-time evolutionary studies could be achieved if molecular clocks are complemented with experimentally validated rates of protein and genome evolution across taxa of interest.

We think it is plausible that there was never a discrete origin of photosynthesis, but that the process may trace back to abiogenic photochemical reactions, some of which may have resulted in the oxidation of water, and at the interface of nascent membranes, membrane proteins, photoactive tetrapyrroles and other inorganic cofactors: much in the same way that ribosomes may have originated at the interface of nascent genetics and protein synthesis [74]. A photosynthetic origin of life is not a new idea [75–77] and abiotic photosynthesis-like chemistry has been recently proposed to have occurred at Gale Crater on Mars [78], and to occur even on Earth [79].

## 5. Materials and methods

### 5.1. Sequence alignments and phylogenetic analysis

The first dataset was retrieved on the 31^st^ of October 2017 to initiate this project. It included a total of 1389 type I RC, CP43 and CP47 protein sequences from the NCBI refseq database using BLAST. From Cyanobacteria, 675 PsaA and PsaB and 685 CP43 and CP47 subunits were retrieved. From anoxygenic phototrophs, 24 PscA and 4 PshA were obtained. This dataset did not contain CBP proteins.

A second dataset was retrieved on the 10^th^ of October 2018 from the same database. This dataset focused on CP43 and CP47 subunits and consisted of a total of 1232 sequences. It included 392 CP43, 465 CBP proteins, and 375 CP47 subunits. 40 CP43 and 40 CP47 sequences from a diverse set of photosynthetic eukaryotes were selected manually and included. In addition, a selection of cyanobacterial and plastid CP43, CP47, AtpA (Alpha), AtpB (Beta), and FtsH were manually selected for an analysis of the rates of evolution as illustrated in Supplementary Figure S14 and described below.

A dataset of Alpha and Beta subunits belonging to cyanobacterial F-type ATP synthase were retrieved from the NCBI refseq on the 31^st^ of August 2019. 507 and 529 cyanobacterial Alpha and Beta sequences were obtained respectively. We retrieved Alpha and Beta homologous for Margulisbacteria (19 and 18 sequences respectively), Sericytochromatia (4 and 2), and Vampirovibrionia (66 and 49) using AnnoTree [80] and searching with the KEGG codes K02111 (F-type Alpha) and K02112 (F-type Beta). A total of 111 AtpA (subunit A) and 176 AtpB (subunit B) from archaeal V-type ATP synthase were retrieved using the sequences from *Methanocaldococcus jannaschii* as BLAST queries.

A dataset of 366 bacterial RNA polymerase subunit β (RpoB) were collected from the NCBI refseq database on the same date as above. We focused on Cyanobacteria (65 sequences) and other phyla known to contain phototrophic representatives, as well as for their potential for allocating calibration points as described below. These included sequences from Firmicutes (32), Chloroflexi (32), Proteobacteria (102), Acidobacteria (34), Chlorobi (26), Gemmatimonadetes (3), Aquifecae (12) and Thermotogae (24). Sequences for *Candidatus* Eremiobacterota (2) recently reported to include phototrophic representatives [81], Margulisbacteria (13), Sericytochromatia (2), and Vampirovibrionia (11) were obtained from AnnoTree using KEGG code K03043 as query. In addition, three extra margulisbacterial RpoB sequences were collected: from *Termititenax persephonae* and *Termititenax aidoneus* retrieved from the refseq database, and from margulisbacterium *Candidatus* Ruthmannia eludens obtained from https://www.ebi.ac.uk/ena/data/view/PRJEB30343 [82]. A second dataset containing only sequences from Cyanobacteria and MSV, in addition to sequences from Archaea was also used. To retrieve the archaeal sequences the subunit B’ (RpoB1) of *Methanocaldococcus jannaschii* was used as a BLAST query. A total 213 sequences across the entire diversity of Archaea were obtained. Two version of these were observed, those with split B (RpoB1 and RpoB2) and those with full-length B subunits: from the latter, 112 full-length B subunits were selected for molecular clock analysis.

All sequences were aligned with Clustal Omega using 5 combined guided trees and Hidden Markov Model Iterations [83]. Given that RpoB sequences are known to contain many clade specific indels [27], we further processed this particular dataset using Gblocks [84] to remove these indels and poorly aligned positions allowing smaller final blocks, gap positions within the final blocks, and less strict flanking positions. This procedure left a total of 903 well-aligned sites for the dataset with only bacterial sequences and 788 for the dataset with archaeal sequences.

Maximum Likelihood phylogenetic analysis was performed with the PhyML online service using the Smart Model Selection mode implementing the Bayesian Information Criterion for parameter selection [85, 86]. Tree searching operations were calculated with the Nearest Neighbour Interchange model. Support was computed using the average likelihood ratio test [87]. Trees were visualised with Dendroscope V3.5.9 [88].

### 5.2. Distance estimation

Distance estimation was performed as a straightforward and intuitive approach to detect variations in the rate of evolution across taxa with known divergence time of at similar taxonomic ranks. This was done in several ways. Distance trees were plotted using BioNJ [89] as implemented in Seaview V4.7 [90] using observed distances and 100 bootstrap replicates. Within-group mean distances were calculated using three different substitution models as implemented in the package MEGA-X [91]: no. of differences, a Poisson model, and a JTT model. A gamma distribution to model rates among sites was used with a parameter of 1.00 and 500 bootstrap replicates.

To compare changes in the rates of evolution as a function of time of key proteins used in this study, we compared the percentage of sequence identity between orthologues in a total of 20 pairs of sequences from representative photosynthetic eukaryotes and Cyanobacteria with known or approximate divergence times. These are listed in Supplementary Table S3. We compared CP43, CP47, Alpha, Beta and FtsH. Of these 17 pairs, the first 15 were comparisons between land plants. The approximated divergence times were based on those recommended by Clarke, et al. [92] after their extensive review of the plant fossil record. These were taken as the average of the hard minimum and soft maximum fossil ages suggested by the authors. The soft maximum age for the earliest land plants was taken as 515 Ma as discussed extensively in Morris, et al. [93]

Another sequence comparison was made between the unicellular red alga *Cyanidioschyzon merolae* and *Arabidopsis thaliana.* The earliest well-accepted fossil evidence for red algae is that of the multicellular *Bangiomorpha* [94, 95] dated to about 1.0 Ga [96]. Recently fossils of multicellular eukaryotic algae have been reported at 1.6 Ga [97–99]. The distance between heterocystous Cyanobacteria and their closest non-heterocystous relatives is represented by a pairwise comparison between *Chroococcidiopsis thermalis* and *Nostoc* sp. PCC 7120. Excellently preserved heterocystous Cyanobacteria have been reported from Tonian deposits older than 0.72 Ga [100]. The largest distance is that between *Gloeobacter* spp. and any other cyanobacterium or photosynthetic eukaryote, representing the distance since the MRCA of Cyanobacteria of uncertain divergence time.

To compare the level of sequence identity between RC proteins, two datasets of 10 random amino acid sequences were generated using the Sequence Manipulation Suit [101]. The datasets contained sequences of 350 and 750 residues. These were independently aligned as described above, resulting in 45 pairwise sequence identity comparisons for each dataset. These random sequence datasets were used as a rough minimum threshold of identity. Alignments of RC proteins were generated using three representative sequences spanning known diversity. Cyanobacterial CP43, CP47, standard D1 and D2 sequences were from *Gloeobacter violaceus, Stanieria cyanosphaera,* and *Nostoc* sp. PCC 7120; Heliobacterial PshA from *Heliobacterium modesticaldum, Heliobacillus mobilis,* and *Heliorestis convoluta;* PscA from *Chlorobium tepidum, Prosthecochloris aestuarii, Chlorobium* sp. GBchlB, *Chloracidobacterium thermophilum* and *Chloracidobacterium* sp. CP2-5A; Protoebacterial L and M from *Methylobacterium* sp. 88A, *Roseivivax halotolerans*, and *Blastochloris sulfoviridis;* and L and M from *Chloroflexus* sp. Y-400-fl, *Roseiflexus castenholzii,* and *Oscillochloris trichoides.*

### 5.3. Quantification of rates of evolution (rationale)

Molecular clocks are conventionally used to estimate divergence times. In general terms, given: 1) a tree topology, which sets the relationship between taxa; 2) a sequence alignment, which sets the phylogenetic distance between taxa; and 3) some known events (calibrations), which set the rates of evolution, the molecular clock can then estimate divergence times. This means that if the tree topology and divergence times for two sets of protein sequences are the same, any differences in phylogenetic distances between these two should only reflect differences in the rate of evolution. Thus, assuming that CP43/CP47 and Alpha/Beta have mainly been inherited vertically in Cyanobacteria and photosynthetic eukaryotes, any difference in phylogenetic distance between the two is the result of differences in the rates of evolution. For example, the level of sequence identity between CP43 in *Cyanidioschizon* and *Arabidopsis* is 78%, and the level of sequence identity between Alpha in the same species is 69%. Given that these plastid-encoded subunits have mostly been inherited vertically since the MRCA of Archaeplastida, then one can argue that Alpha is evolving somewhat faster than CP43. This is because faster rates of protein evolution should lead to faster rates of change resulting in a faster decrease in the level of sequence identity (increase in phylogenetic distance). Now, because CP43 and CP47 are paralogues, we can then use this approach to estimate the rates of evolution at the moment of duplication under a given number of specific scenarios. For example, assuming that the MRCA of Cyanobacteria occurred at 2.0 Ga, we can then model how the rates of evolution would change at the moment of duplication if this occurred at 2.5, 3.0, 3.5 Ga, or at any other time point of interest. Because these set of proteins have likely achieved mutational saturation at the largest distance, the rate of evolution at the point of duplication will be an underestimation (slower than it should be), given that saturation would hide substitution events. Similarly, keeping the time of duplication constant, we can then model how the rates of evolution would change across the tree if the MRCA of Cyanobacteria is assumed to have occurred at 2.0, 2.5, 3.0 Ga, or at any other time of interest. Therefore, even though it is difficult to determine the absolute time of origin of Cyanobacteria, or of the early duplications of the core subunits of ATP synthase, it is possible to determine what rates of evolution are required to fulfil any particular scenario.

### 5.4. Quantification of rates of evolution

To measure rates of evolution of CP43/CP47 and Alpha/Beta a total of 19 sequences from photosynthetic eukaryotes and 4 cyanobacterial sequences per subunit were selected. A standardized tree topology was constructed from consensus evolutionary relationships as illustrated in Supplementary Figure S14a: 1) Relationships between land plants was taken from ref. [92] 2) It is well established that the divergence of red algae predates the MRCA of land plants. 3) Ponce-Toledo, Deschamps [102] recently suggested that *Gloeomargarita* is the closest living cyanobacterial relative to the plastid ancestor and predated the emergence of heterocystous cyanobacteria, but see also [103]. 4) The clade containing *Chroococcidiopsis thermalis* PCC 7203 is one of heterocystous Cyanobacteria’s closest non-heterocystous relatives [104, 105]. 5) *Gloeobacter* spp. is the earliest branching and well-described genus of Cyanobacteria capable of oxygenic photosynthesis [105–109].

Calibration points were allocated as shown in Supplementary Figure S14a and listed in Table 2. Nodes 1 to 15 were applied following the justifications and best practice listed in ref. [92] and using the 515 Ma older calibration as suggested in ref. [93] Node 16 represents here the MRCA of red and green lineages of photosynthetic eukaryotes. The minimum age was set to 1.0 Ga based on the *Bangiomorpha* fossil as described above [94, 95]. The maximum age was set to 1.8 Ga, which is similar the oldest ages reported in recent molecular clock analyses for the MRCA of photosynthetic eukaryotes and it is also similar to the age of the earliest plausible fossilised unicellular eukaryotes [110, 111]. That said, compelling fossil evidence for eukaryotes could go as far back as 2.1 Ga [112]. Node 17 marks the divergence of heterocystous Cyanobacteria and it was given a minimum age of 0.72 Ga based on the recently described fossils of filaments bearing clear heterocysts from the Tonian period [100]. A maximum age of 1.65 Ga was given to this node, based on the recent report of heterocystous cyanobacteria from the Gaoyuzhuang Formation [113]. Although, it is unclear if reported akinetes that are older than 1.65 Ga are truly that [10], we considered that the 0.72-1.65 Ma range for the origin of heterocystous Cyanobacteria was a reasonably broad constraint for the purpose of this experiment, but see below.

**Table 2.** Calibration points used in this study

| Node label | Relationship | Max age (Ma) | Min age (Ma) |
| --- | --- | --- | --- |
| 1 | *Arabidopsis* v *Populus* | 127 | 82 |
| 2 | *Arabidopsis* v *Sorghum* | 248 | 124 |
| 3 | *Chloranthus* v *Liriodendron* | 248 | 98 |
| 4 | *Arabidopsis* v *Illicium* | 248 | 124 |
| 5 | *Arabidopsis* v *Nymphaea* | 248 | 124 |
| 6 | *Arabidopsis* v *Amborella* | 248 | 124 |
| 7 | *Welwitschia* v *Pinus* | 309 | 121 |
| 8 | *Welwitschia* v *Cryptomeria* | 309 | 147 |
| 9 | *Welwitschia* v *Ginkgo* | 366 | 107 |
| 10 | *Welwitschia* v *Cycas* | 366 | 306 |
| 11 | *Arabidopsis* v *Cycas* | 366 | 306 |
| 12 | *Arabidopsis* v *Psilotum* | 454 | 388 |
| 13 | *Arabidopsis* v *Anthoceros* | 515 | 420 |
| 14 | *Arabidopsis* v *Physcomitrella* | 515 | 420 |
| 15 | *Arabidopsis* v *Marchantia* | 515 | 449 |
| 16 | *Arabidopsis* v *Cyanidioschyzon* | — | 1000 |
| 17 | *Nostoc* v closest non-heterocystous strain | 1560 (or none) | 720 |
| 18 | *Arabidopsis* v *Gloeobacter* | Variable (or none) | Variable (or none) |
| 19 | CP43 v CP47 or AtpA v AtpB duplications | Variable | Variable |
| 20 | *Richelia intracellularis* v closest neighbouring sequence | 250 | 100 |
| 21 | Chroococcales cyanobacteria | — | 2016 |
| 22 | *Termititenax* | 396 | 140 |
| 23 | *Candidatus Ruthmannia eludens* v GCA 002716725.1 | 660 | 542 |
| 24a | GB GCA 001899365.1 Koala v GB GCA 001917115.1 Homo | 201 (or 55) | — |
| 24b | GB GCA 001917115.1 *Homo* v GB GCA 000980455.1 Homo | 201 | — |
| 25 | GB GCA 001917115.1 *Homo* v *Vampirovibrio chlorellavorus* | 1800 | — |
| 26 | *Polynucleobacter necessarius* | 444 | — |
| 27 | *Bradyrhizobium* | 86 | — |
| 28 | Rickettsias | — | 1800 |
| 29 | *Wolbachia* and *Anaplasma* | 395 | — |
| 30 | Phototrophic Chlorobi | — | 1640 |
| 31 | Chromatiaceae | — | 1640 |

Node 18 denotes the MRCA of Cyanobacteria. The age for this node is highly debated ranging from before to after the GOE. Node 19 represents the duplication events leading to CP43 and CP47, and to Alpha and Beta. To calculate the rates of evolution under different scenarios, node 18 and node 19 were varied. Firstly, a molecular clock was run using a scenario that assumed that the MRCA of Cyanobacteria postdated the GOE. To do this, node 18 was set to be between 1.6 and 1.8 Ga, which emulates results reported in recent studies [8, 11]. This was compared to a scenario that assumed that the MRCA of Cyanobacteria antedated the GOE, and thus node 18 was set to be between 2.6 and 2.8 Ga, which simulates other evolutionary scenarios [103, 114]. In both cases, the duplication event (node 19) was set to be 3.5 Ga old, or changed as stated in the main text, by assigning a gamma prior at the desired time fixed with a narrow standard deviation of 0.05 Ga. In a separate experiment, the age of the duplication was varied while maintaining node 18 restricted to between 1.6 and 1.8 Ga, while node 19, the root, was set with a gamma prior with an average varied from 0.8 to 4.2 and with a narrow standard deviation of 0.05 Ga.

The period of time between the duplication event (node 19), which led to the divergence of CP43 and CP47, and the MRCA of Cyanobacteria (node 18), we define as ΔT. ΔT is calculated as the subtraction of the mean age of node 19 and node 18. For PSII, we used node 18 from the CP43 subunit and for ATP synthase we used node 18 from the Alpha subunit. In consequence, varying the age of the duplication from 0.8 to 4.2 Ga allows changes in the rate of evolution to be simulated with varying ΔT, ranging from 0.2 to 2.5 Ga. That is to say, it allows the rate of evolution at the point of duplication to be calculated if it occurred at any point in time before the MRCA of Cyanobacteria.

Molecular clocks were calculated with Phylobayes 3.3f using a log normal autocorrelated molecular clock model under the CAT+Γ non-parametric model of amino acid substitution and a uniform distribution of equilibrium frequencies. We preferred a CAT model instead of a CAT+GTR as the latter only outperforms the former on sequence alignments of over 1000 sites and it is also much less computationally expensive. Four discrete categories for the gamma distribution were used and four chains were executed in parallel through all experiments carried out in this study. The instant rates of evolution, which are the rates at each internal node in the tree, were retrieved from the output files of Phylobayes. These rates are calculated by the software as described by the developers elsewhere [115, 116] and are expressed as amino acid changes per site per unit of time.

As an additional comparison, we calculated rates of evolution for cyanobacterial FtsH subunits, which AAA ATPase domain is structurally similar to the catalytic head of ATP synthase [117]. Cyanobacterial FtsH are involved in membrane protein quality control, with some isoforms targeting specifically PSII subunits. It features a late duplication event leading to cyanobacterial FtsH1 and FtsH2 subunits (using the nomenclature of Shao, et al. [40]). It was shown previously that the duplication leading to FtsH1 and FtsH2 occurred after the divergence of *Gloeobacter* spp. Genes encoding plastid FtsH subunits are encoded in the nuclear genome of photosynthetic eukaryotes. Because not all the selected species had fully sequenced nuclear genomes, only those with available FtsH sequences at the time of this study were used. The tree topology shown in Supplementary Figure S14b was used as template for the calculation of the rate of evolution and was based on the topology presented by Shao, et al. [40] It was concluded that the Cyanobacteria-inherited closest paralog to FtsH1 and FtsH2 in photosynthetic eukaryotes was also acquired before their initial duplication. Therefore, from all FtsH paralogs in photosynthetic eukaryote genomes, those with greater sequence identity to cyanobacterial FtsH1/2 were used. Because this duplication is specific to Cyanobacteria, a few additional strains were included in this tree following well-established topologies [103, 105]. Calibrations were placed as indicated in Supplementary Figure S14b. To test the change in the rate of evolution at the time of duplication in comparison with CP43/CP47, node 19 was set to 1.6-1.8 Ga or 2.6-2.8 Ga and molecular clocks were run as described in the preceding paragraph.

Finally, we conducted a large molecular clock using the combined 897 CP43 and CBP sequences, including 40 eukaryotic CP43 sequences, to test whether using a more complex phylogeny would result in rates of evolution substantially different to those calculated with the method described above. Calibrations were assigned as illustrated in Supplementary

Figure S15. Cross-calibrations were used across paralogs constraining the origin of heterocystous Cyanobacteria. In this case, only the minimum constraint of 0.72 Ga was used with no maximum constraint to allow greater flexibility. Additional calibrations were assigned also across paralogs (point 20 in Supplementary Figure S15), this was considered as the node made by *Richelia intracellularis* and its closest sister sequence, as implemented in ref. [114] This strain is a specific endosymbiont of a diatom and its divergence was set to be no older than the earliest discussed age for diatoms [118]. The root equivalent to the MRCA of Cyanobacteria (divergence of *Gloeobacter* in CP43) was not calibrated. The root of the tree was varied: first it was given a maximum age of 4.52 Ga as recently implemented and justified by Betts, et al. [11] as the earliest plausible time in which the planet was inhabitable after the moon forming impact [119], and no minimum age was used. A second tree was executed with no constraint on the root and no root prior. A third root was implemented constrained to be between 2.3 Ga (the GOE) and 3.2 Ga. The latter date represents the age of the cyanobacteria-like well-preserved microbial mats of the Berberton Greenstone Belt in South Africa and neighbouring Eswatini [66]. Rates were obtained using the autocorrelated CAT+Γ model as described above. Because these root constraints did not have a strong effect in the overall estimated rates, we carried out an additional control applying an uncorrelated gamma clock model [120] with a root constrained at 4.52 Ga and no minimum age.

### 5.5. Molecular clock of RpoB and concatenated ribosomal proteins

The primary objective of this experiment was not to determine the absolute time of origin of Cyanobacteria, but to understand the spans of time between Cyanobacteria and their relatives. We also wanted to understand what rates of evolution are associated with those spans of time and how these change under different evolutionary scenarios. To do this, we applied a molecular clock to the phylogeny of RpoB sequences described above. We implemented 12 calibrations. The calibrations were assigned on the phylogeny as shown in Supplementary Figure S16 and listed in Table 2. A set of calibrations consisted of the earliest unambiguous evidence for Chroococcales Cyanobacteria of the Belcher group (point 21), the age of which has been recently revisited to 2.01 Ga [121]. This was assigned to the younger node from where Chroococcales strains branch out in the tree, with no maximum restrictions. The appearance of heterocystous Cyanobacteria were restricted from 0.72 Ga and 1.56 Ga as described above. No constraints on the node representing the MRCA of Cyanobacteria were used. However, for rigor, we also tested an alternative single calibration on the node representing the MRCA of Cyanobacteria with a maximum of 2.01 Ga and no minimum, and with no other calibrations in the clade. This considered a scenario in which crown group Cyanobacteria are younger than the Belcher fossils.

In addition to cyanobacterial calibrations, we also applied the often-used biomarker evidence for phototrophic Chlorobi and Chromatiaceae at 1.64 Ga [122], see for example refs. [9, 63] These were used as a minimum with no maximum constraints (node 30 and 31 respectively in Supplementary Figure S16).

A set of calibrations were chosen from well-described symbiotic relationships. Two of these are known in the phylum Margulisbacteria. The first one is *Termititenax,* these are specific ectosymbionts of spirochetes that live within oxymonad protists in the gut of diverse termites and cockroaches [43]. The two *Termititenax* sequences used in this analysis clustered together and therefore we calibrated this node to be between 396 Ma, some of the oldest fossil evidence of insects [123] and 140 Ma for *Mastotermes nepropadyom,* a Jurassic fossil termite [124] (node 22). This was done under the assumption that this symbiotic relationship may have started before the divergence of termites and cockroaches, as suggested in ref. [43] The second one is *Candidatus* Ruthmannia eludens, a cell-type specific endosymbiont of placozoans, early-branching metazoans. This symbiont has been detected in all haplotypes examined, regardless of geographical location or sampling time [82, 125], which suggest that this association may be as old as placozoans. Therefore, we used a minimum calibration of 542 Ma as the Cambrian explosion of animal diversity and 660 Ma, the earliest biomarker evidence for desmosponges [126] (node 23), which should antedate the divergence of placozoans. This calibration was assigned to the node separating *Candidatus* Ruthmannia eludens from its closest sister sequence.

Similar to Margulisbacteria, members of the clade Vampirovibrionia have been reported to form close associations with eukaryotes. The clade Gastranaerophilales is thought to be composed mostly of strains that inhabit the animal gut. Thus, we calibrated the node separating two strains isolated from human and koala faeces that clustered together with a maximum age of 201 Ma, representing the Jurassic split of marsupials and placental mammals [127] (node 24a). However, the koala and human sequences were embedded in the Gastranaerophilales clade within other sequences from the human gut. Because of this, we trialled changing this calibration to 55 Ma instead, the oldest primate fossil [128] and assuming that the retrieved sequences from the human gut had a common ancestor younger than the MRCA of primates. Alternatively, we tested moving this calibration to the ancestral nodes of the clade that included all the human gut sequences (node 24b). Gastranaerophilales is closely related to the order Vampirovibrionales, which include *Vampirovibrio chlorellavorus*. This strain is a predator of the eukaryotic green algae *Chlorella* [129], and therefore we trialled a calibration assuming that Gastranaerophilales and Vampirovibrionales radiated after the MRCA of eukaryotes (node 25). We thus assigned a maximum calibration to this node of 1.8 Ga representing the earliest described plausible eukaryote fossils [110] and no minimum age.

Another highly specific obligate symbiosis is that of the betaproteobacterium *Polynucleobacter necessarius* and ciliates of the genus *Euplotes* (Spirotrichea) [130]. *Polynucleobacter* has close free-living phototrophic relatives within the same genus [130]. We set the node separating the phototrophic and non-phototrophic *Polynucleobacter* (node 26) a maximum age of 444 Ma for the oldest fossil evidence of spirotrichs, as implemented in Parfrey et al. [131], and which predates the radiation of the genus *Euplotes* [132].

Another well-known association is that of the soil bacteria *Bradyrhizobium* and legumes. Thus we gave the node separating *Bradyrhizobium* spp. from its closest relative in the RpoB tree, *Xanthobacter autotrophicus*, a maximum age of 86 Ma for Rosids, which contain legumes [93] (node 27).

The Rickettsiales are Alphaproteobacteria that exists in very close association with eukaryotes [133]. An association that may reach to the lineage leading to the origin of mitochondria [134]. Therefore, we assumed that the divergence of Rickettsiales occurred before the MRCA of eukaryotes and gave this node a minimum age of 1.8 Ga [110] (node 28). Finally, the family Anaplasmataceae contains bacteria that exists in close association with insects as endosymbionts (e.g. *Wolbachia)* or as parasite vectors (e.g. *Anaplasma).* Therefore, we set a maximum constraint for the MRCA of *Wolbachia* and *Anaplasma* (node 29), excluding *Neorickettsia,* to be as old as the earliest evidence for insects about 395 Ma ago [123].

To constrain the age of the root, we first set a broad gamma prior with an average of 3.8 Ga and a standard deviation of 0.5 Ga. We found this to perform well and used it as benchmark to compare with a range of evolutionary models and the effects of key calibrations (Supplementary Figure S8). Alternatively, we applied a broad calibration on the root with a maximum of 4.52 Ga as described above and a minimum of 3.41 Ga, which is the earliest well-accepted evidence for photosynthesis [47]. This evidence was hypothesized to be anoxygenic in ref. [47] Therefore, Bacteria should be at least as old as the earliest evidence for photosynthesis under conventional evolutionary scenarios.

To further understand the effect that a distant outgroup would have on the estimated divergence time, and to measure the rate of evolution of RpoB at the divergence of Bacteria and Archaea, we repeated the clock including 112 diverse archaeal sequences, in addition to Thermotogae, MSV, and Cyanobacteria, but removing all other clades as a compromise between robustness and computing time. Calibrations were assigned on MSV and Cyanobacteria as described above. Relaxed molecular clocks were computed using an autocorrelated log normal clock and applying a CAT+Γ model. This was compared against a CAT+Γ and a CAT+GTR+Γ using birth-death priors and soft bounds on the calibrations allowing for 2.5% tail probability falling outside the minimum and maximum boundary, or 5% in the case of a single boundary. In addition, we also compared to an uncorrelated gamma model (Supplementary Figure S8).

We compared the RpoB molecular clock (CAT+Γ) with that of a clock executed using a dataset of concatenated ribosomal proteins of 2596 aligned sites, obtained from an independent study [38]. In total, 157 sequences were used including 56 cyanobacterial sequences, 14 sequences from Vampirovibrionia, 9 from Margulisbacteria, 9 from Thermotogae and 78 from Archaea. A set of calibrations were used including those two in Cyanobacteria (point 17 and 21), a calibration on the MRCA of Gastranaerophilales with a maximum 201 Ma (point 24b), and a root calibration between 4.52 and 3.41 Ga as described above.

### 5.6 Ancestral sequence reconstruction

Ancestral sequence reconstruction (ASR) of D1 and D2 amino acid sequences was carried out with a dataset collected on the 17^th^ of November 2017. Duplicates and partial sequences were removed, leaving 755 D1 and 248 D2 sequences. CD-HIT [135] was used to remove sequences with greater than 92% sequence identity to create a representative sample. The L and M RC subunits from 5 strains of Proteobacteria were used as outgroup. The final alignment did not include the atypical variant D1 sequence from *Gloeobacter kilaueensis* (NCBI accession AGY58976.1) as this showed an unstable phylogenetic position in this dataset. Maximum likelihood trees used as input for ASR were computed with PhyML using Smart Model Selection [86]. The LG substitution model with observed amino acid frequencies and four gamma rate categories exhibited the best log likelihood (LG+Γ+F) (Supplementary Table S7). We used the top four models for tree reconstruction (LG+Γ+F, LG+Γ+I+F, LG+Γ and LG+Γ+I) in addition to another tree computed using the WAG substitution model with observed amino acid frequencies and four gamma rate categories (WAG+Γ+F). These trees were used as input trees to calculate maximum likelihood ancestral states at each site for the node corresponding to the homodimeric reaction protein, D0. Three ASR programs were used for the reconstructions: FastML [136], Lazarus [137] (a set of Python scripts which wraps PAML [138]) and MEGA-7 [139]. The substitution model used by all three programs corresponded to the substitution model used for the specific input tree in PhyML and the branch lengths of the trees were fixed. In FastML, a maximum likelihood method of indel reconstruction using a probability cut-off of 0.7 was used. In MEGA-7, all sites were used for analysis with no branch swap filter. In Lazarus, the ‘—gapcorrect’ option was used to parsimoniously place indels.

### 5.7. Structural analysis

The following crystal structures were used in this work: the crystal structure of PSII from *Thermosynechococcus vulcanus,* PDB ID: 3wu2 [13]; the anoxygenic type II RC of *Thermochromatium tepidum,* PDB ID: 5y5s [140]; the homodimeric type I RC from *Heliobacterium modesticaldum,* 8v5k [141]; PSI from *Thermosynechococcus elongatus,* PDB ID: 1jb0 [142]; and the cryo-EM IsiA structure from *Synechocystis* sp. PCC 6803 [143]. Structures were visualised using Pymol^TM^ V. 1.8.2.2 (Schrodinger, LLC) and structural overlaps were carried out with the CEAlign plugin [144].

## 6. Acknowledgements

This work was done with the support of a Leverhulme Trust grant (RPG-2017-223 to T.C. and A.W.R.), a UKRI Future Leaders Fellowship (MR/T017546/1 to T.C.) a Royal Society University Research Fellowship (P.S.-B.), and the Biotechnology and Biological Sciences Research Council (BB/K002627/1 and BB/L011206/1 to A.W.R.). The authors are grateful to Dr. Travis J. Lawrence and Prof. Christoph Heubeck for feedback on a version of this manuscript. Computing resources were provided by the HPC facility at Imperial College London. The authors declare no conflict of interests.

