## Supplementary Figures and Text for "Time-resolved comparative molecular evolution of oxygenic photosynthesis"

**Content**

Suppementary Figure S1	Page 2

Suppementary Figure S2	Page 4

Suppementary Figure S3	Page 5

Suppementary Figure S4	Page 7

Suppementary Figure S5	Page 8

Suppementary Figure S6	Page 9

Suppementary Figure S7	Page 10

Suppementary Figure S8	Page 12

Suppementary Figure S9	Page 15

Suppementary Figure S10	Page 16

Suppementary Figure S11	Page 17

Suppementary Figure S12	Page 18

Suppementary Figure S13	Page 19

Suppementary Figure S14	Page 20

Suppementary Figure S15	Page 21

Suppementary Figure S16	Page 23

Suppementary Figure S17	Page 26

Supplementary Text S1	Page 27

Supplementary Text S2	Page 29

References	Page 29

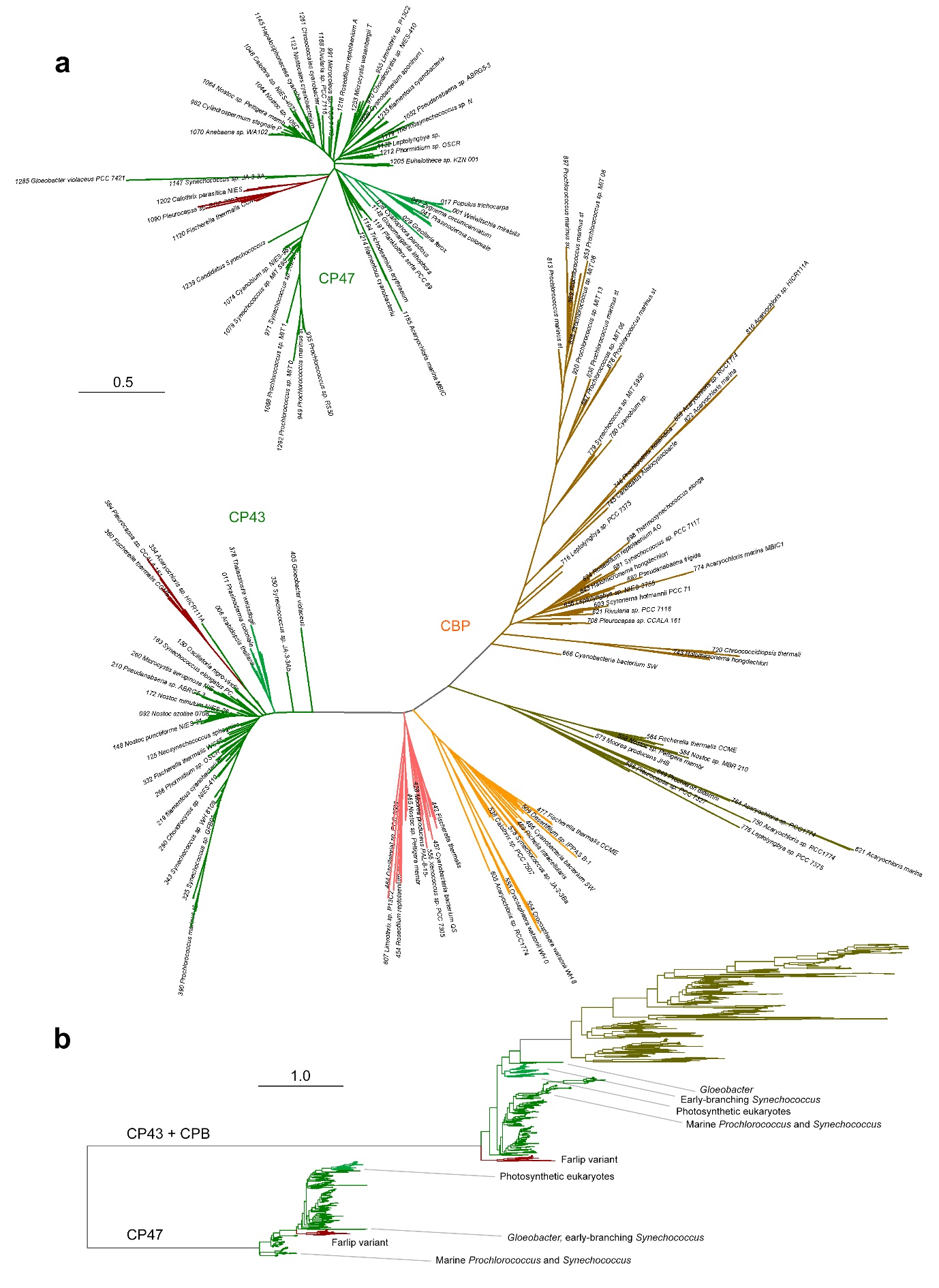

**Supplementary Figure S1.** Unrooted ML tree of CP43/CPB and CP47 subunits. **a** Unrooted trees of CP47 (top) and CP43/CPB (bottom). **b** A ML tree using an alignment that included CP43/CPB and CP47 sequences. The generated tree highlights the long distance between CP47 and CP43/CPB, however the faster evolving CPB sequences in combination with the long branches that separate CP43 and CP47 produces a phylogeny with strong long-branch attraction artefacts and an inverted CP43 topology regardless of model selection.

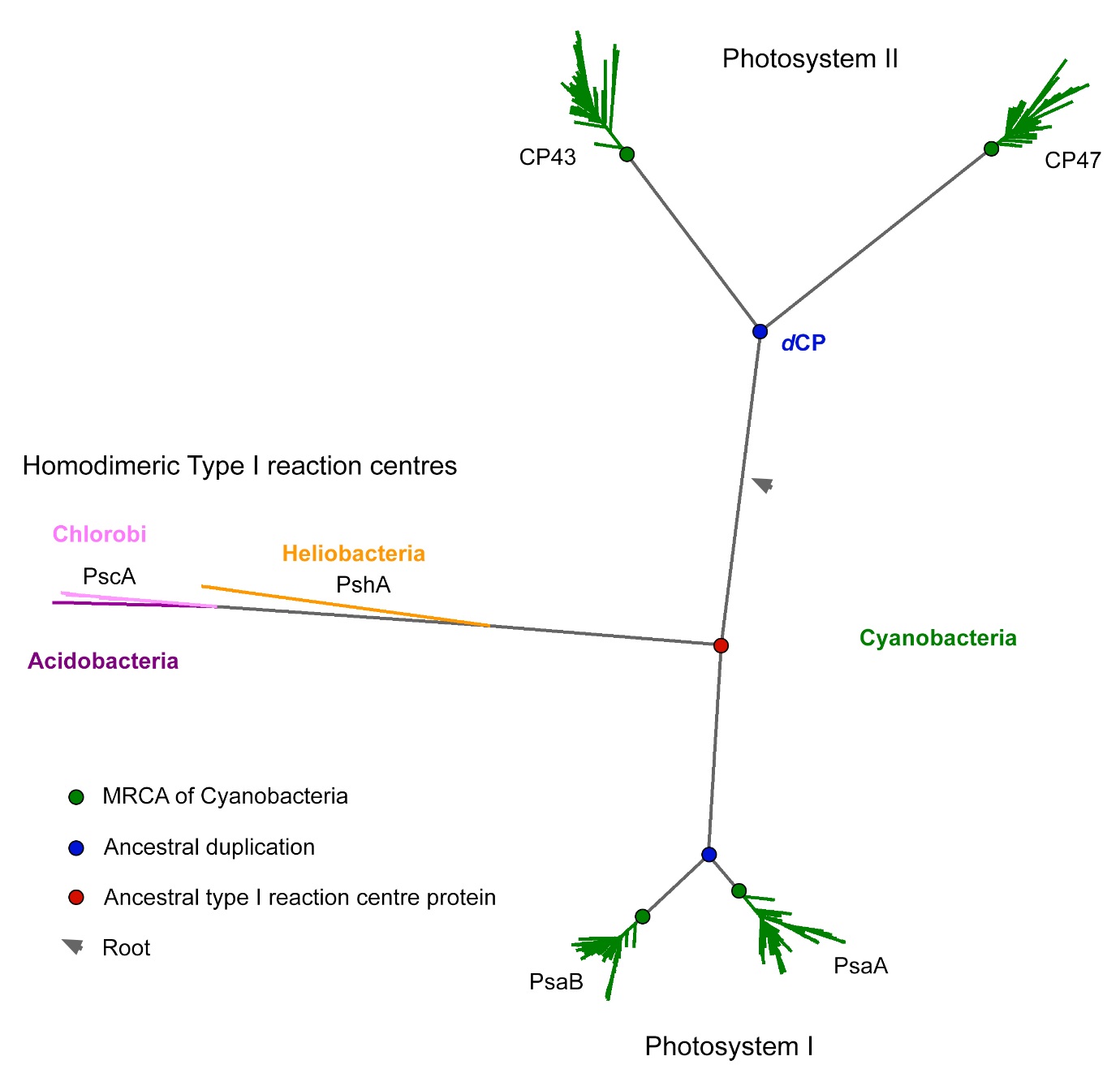

**Supplementary Figure S2.** ML tree of type I reaction centre proteins including CP43 and CP47. This tree highlights the long distance between the PSII antenna proteins and type I reaction centres. Type I reaction centre proteins are more likely to share a common ancestor to the exclusion of CP43 and CP47. This is supported by the observation that type I reaction centre proteins share greater sequence and structural identity (see also Supplementary Table S1). If that is the case, the root would therefore be placed at the longest branch (arrow). This placement is consistent with the phylogenetic relationships of reaction centre proteins showing that type I and type II reaction centres make separate monophyletic lineages [1, 2] and as illustrated in the next supplementary figure. The blue dot represent the ancestral duplications that are specific to the evolution of oxygenic photosynthesis. Green dots represent the MRCA of Cyanobacteria, which inherited well-defined CP43/CP47; and PsaA and PsaB, the core subunit of photosystem I (PSI). Therefore, the duplications of the ancestral core photosystem proteins of oxygenic photosynthesis predate the MRCA of Cyanobacteria by a set amount of time.

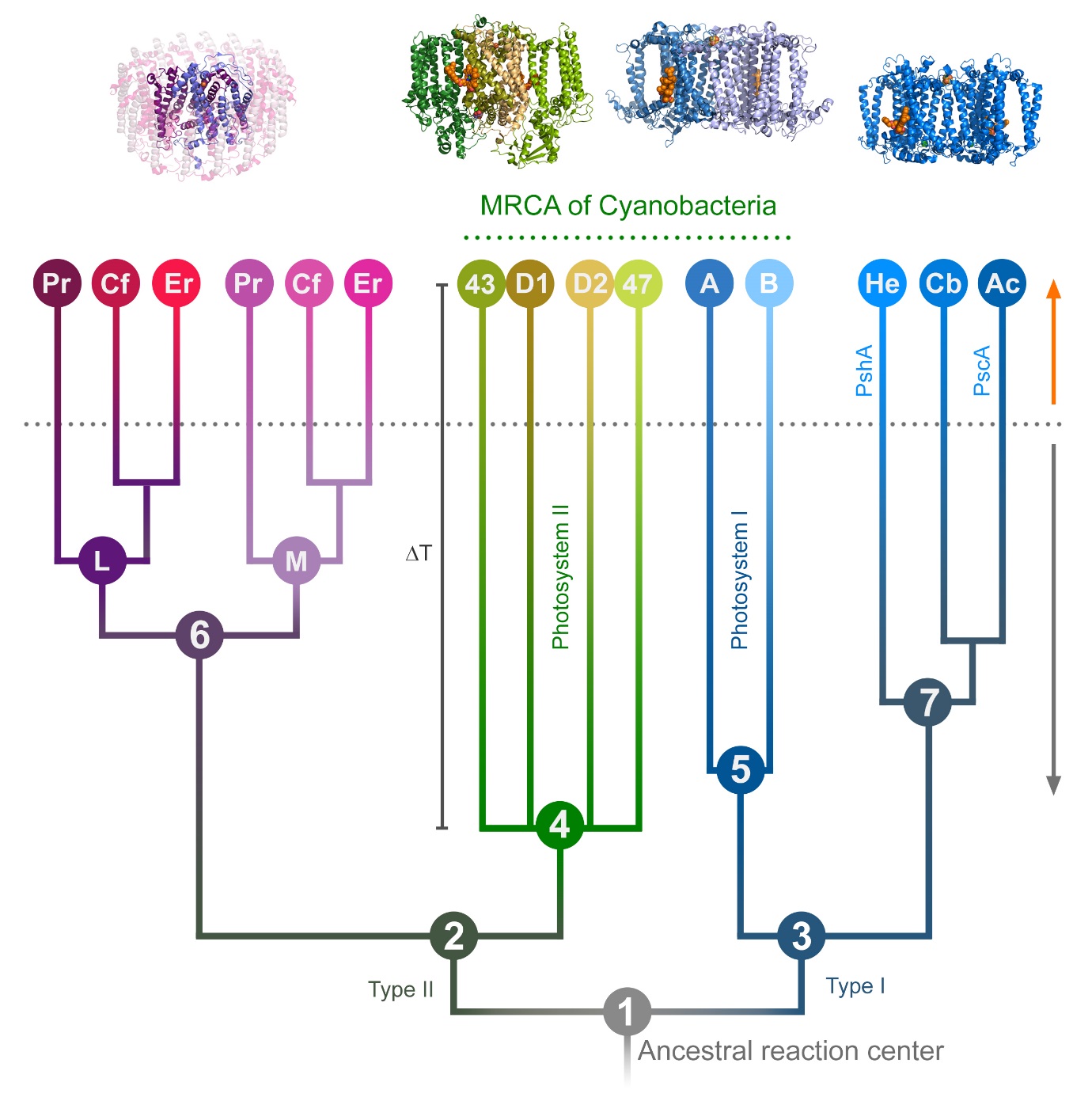

**Supplementary Figure S3.** Schematic representation of the evolution of reaction centre proteins. The evolutionary relationships of reaction centre proteins have been explained in detail before [1-3], these are based on phylogenetic analysis in combination with the comparative structural biology of the photosystems. All reaction centre proteins have a common origin (**1**). L, M, D1, and D2, the type II reaction centre proteins, make a monophyletic clade (**2**). PsaA (A), PsaB (B), PshA, and PscA, the type I reaction centre proteins, make a distinct monophyletic clade (**3**). Therefore, the earliest diversification event in the evolution of reaction centre proteins is the divergence of type II and type I reaction centre proteins. PSII retains antenna subunits (CP43/CP47) homologous to the antenna domain found in type I reaction centre proteins, but absent in anoxygenic type II reaction centres made up of L and M. The long distance between CP43/CP47 and type I reaction centres, as seen in the preceding figure, is consistent with these proteins branching out together with D1/D2 before the diversification of type I reaction centre proteins (events **1**, **2** and **4**). Analysis of the rate of evolution of type II [4] and type I [5] reaction centres suggests that the core duplications leading to the heterodimeric PSII (**4**) and heterodimeric PSI (**5**) of oxygenic photosynthesis antedate the duplication leading to the heterodimeric core of anoxygenic type II reaction centres (**6**), and the split leading to PshA (Firmicutes) and PscA (Chlorobi, Acidobacteria) (**7**), respectively. Overall, the evolutionary relationships of reaction centre proteins indicate that their early stages of diversification (**1** to **7**, grey arrow) likely antedated the diversification events that lead to the known diversity of phototrophs (dotted line, orange arrow) regardless of whether these events happened through vertical or horizontal inheritance. Proteobacteria (Pr), Chloroflexi (Cf), *Candidatus* Eremiobacterota (Er), Heliobacteria (He), Chlorobi (Cb), Acidobacteria (Ac).

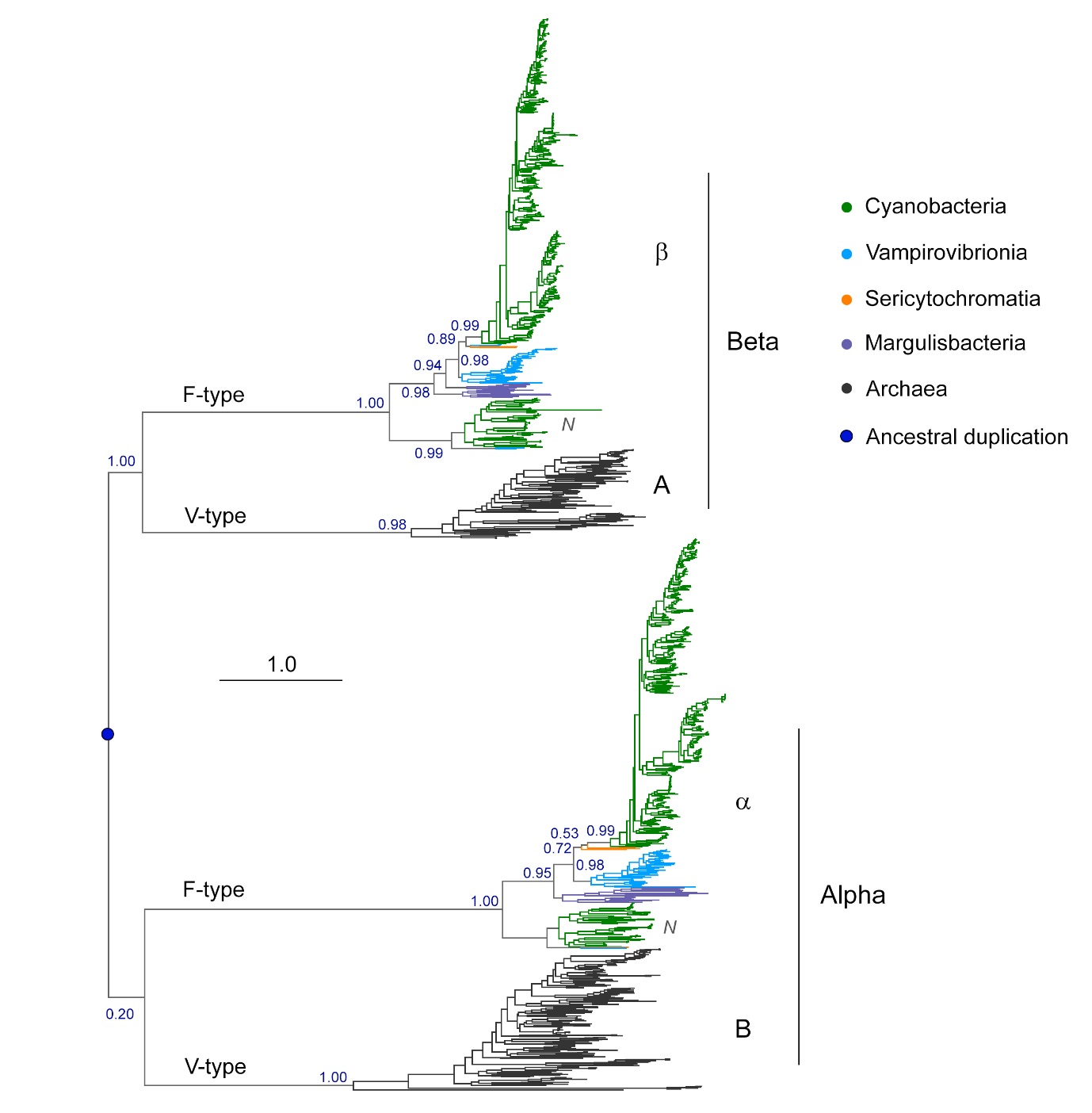

**Supplementary Figure S4.** ML tree of ATP synthase catalytic head subunits with emphasis on Cyanobacteria and their non-photosynthetic closest relatives. The evolution of ATP synthase is complex [6]. Overall, F-type is considered of bacterial origin and V-type of archaeal origin. Interdomain events of horizontal gene transfer are known and some strains of Margulisbacteria have been reported to have acquired V-type ATP synthase [7], for example. These are not included here. Alpha and Beta subunits share distant homology with other proteins containing P-loop NTPase domains. It is generally considered that the ancestral duplication that allowed the heterohexamerization of the catalytic head (F_1_, V_1_) is a very ancient event that occurred before the diversification of most forms of life and may even predate the LUCA. For simplicity, we refer in this work to the A and β subunit of ATP synthase together as Beta, and to B and α subunit as Alpha. *N* denotes Na^+^-translocating N-type ATPase of the bacterial F-type. Scale bar, number of substitutions per site.

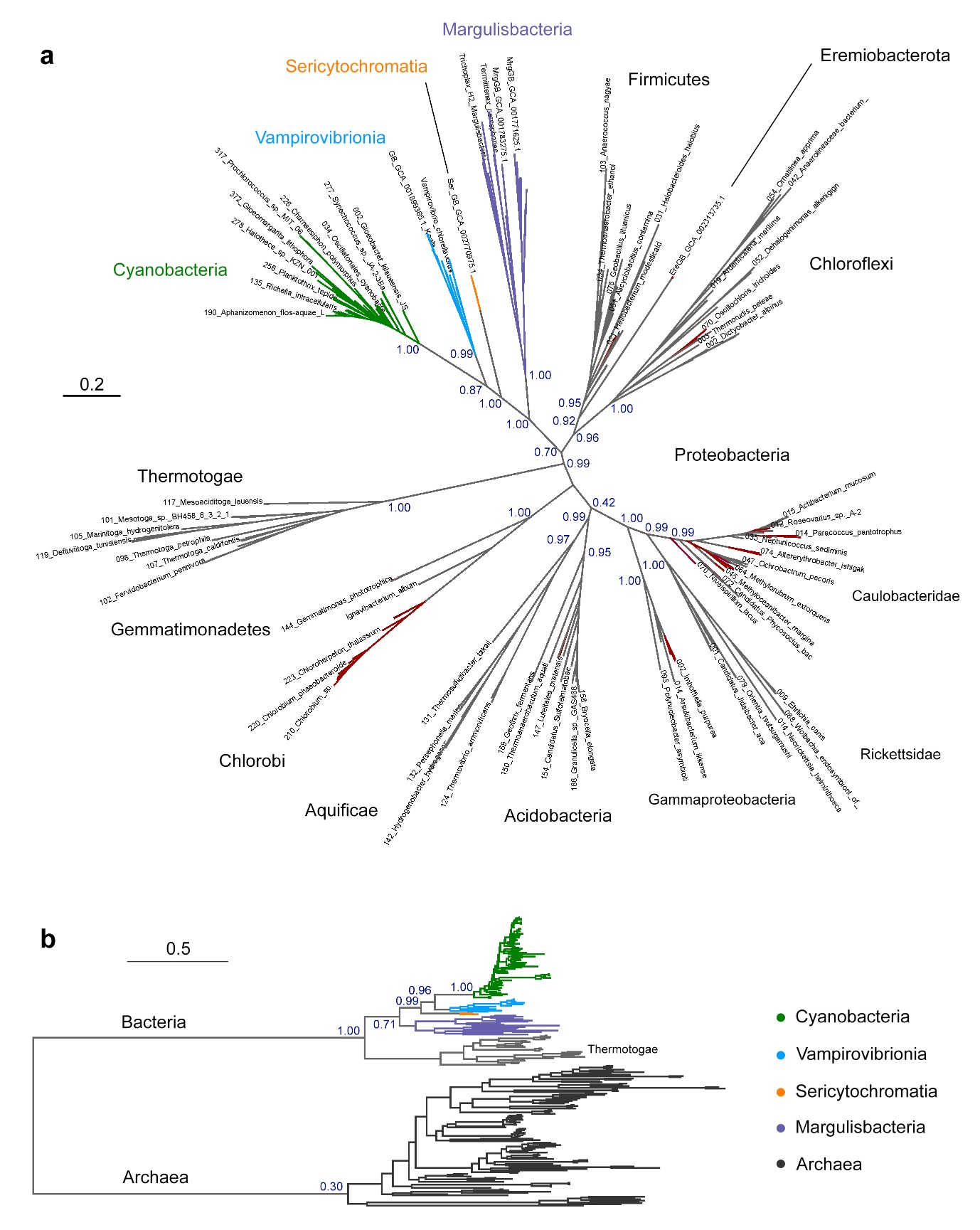

**Supplementary Figure S5**. ML trees of RNA polymerase subunit β (RpoB). **a** A tree of bacterial RpoB with emphasis on clades known to contain phototrophic representatives, Cyanobacteria and their closest non-photosynthetic relatives. Red branches denote phototrophic strains or clades. **b** A ML tree of selected bacterial RpoB and a diverse range of archaeal homologous sequences. Scale bars, number of substitutions per site.

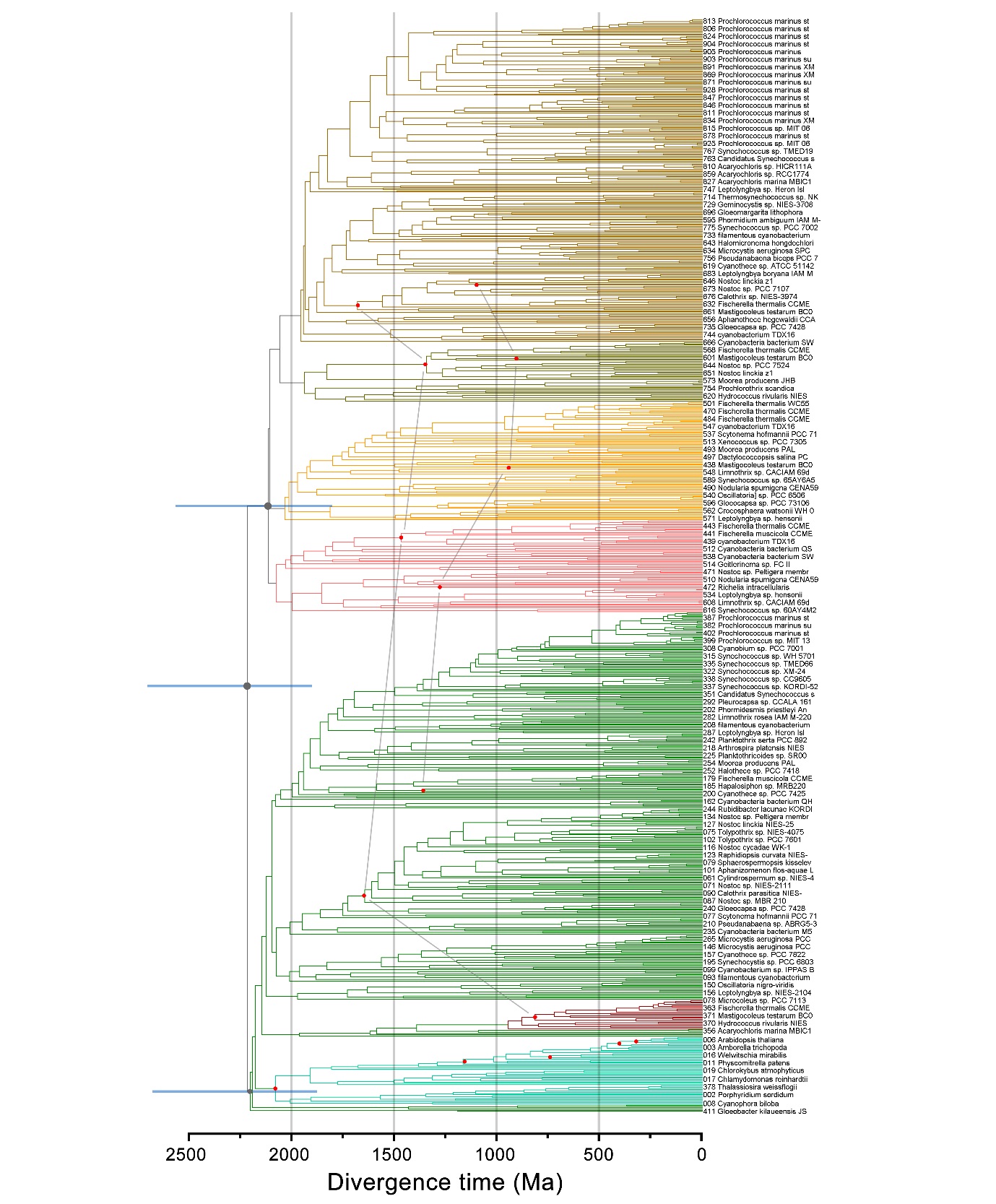

**Supplementary Figure S6.** Bayesian relaxed molecular clock of CP43 and CBP. This clock was calculated using a log normal autocorrelated molecular clock with a CAT+Γ non-parametric model of amino acid substitutions as described in the Materials and Methods. The blue bars represent 95% confidence intervals at the deepest nodes. Red dots mark calibrated nodes and those connected by grey lines represent cross-calibrations across paralogues. The variation in the mean ages of the calibrated nodes is the result of the very broad constraints used in combination with the inherent variation of the rates of evolution within clades. The tree is colour-coded as seen in Figure 1 and Supplementary Figure 1a.

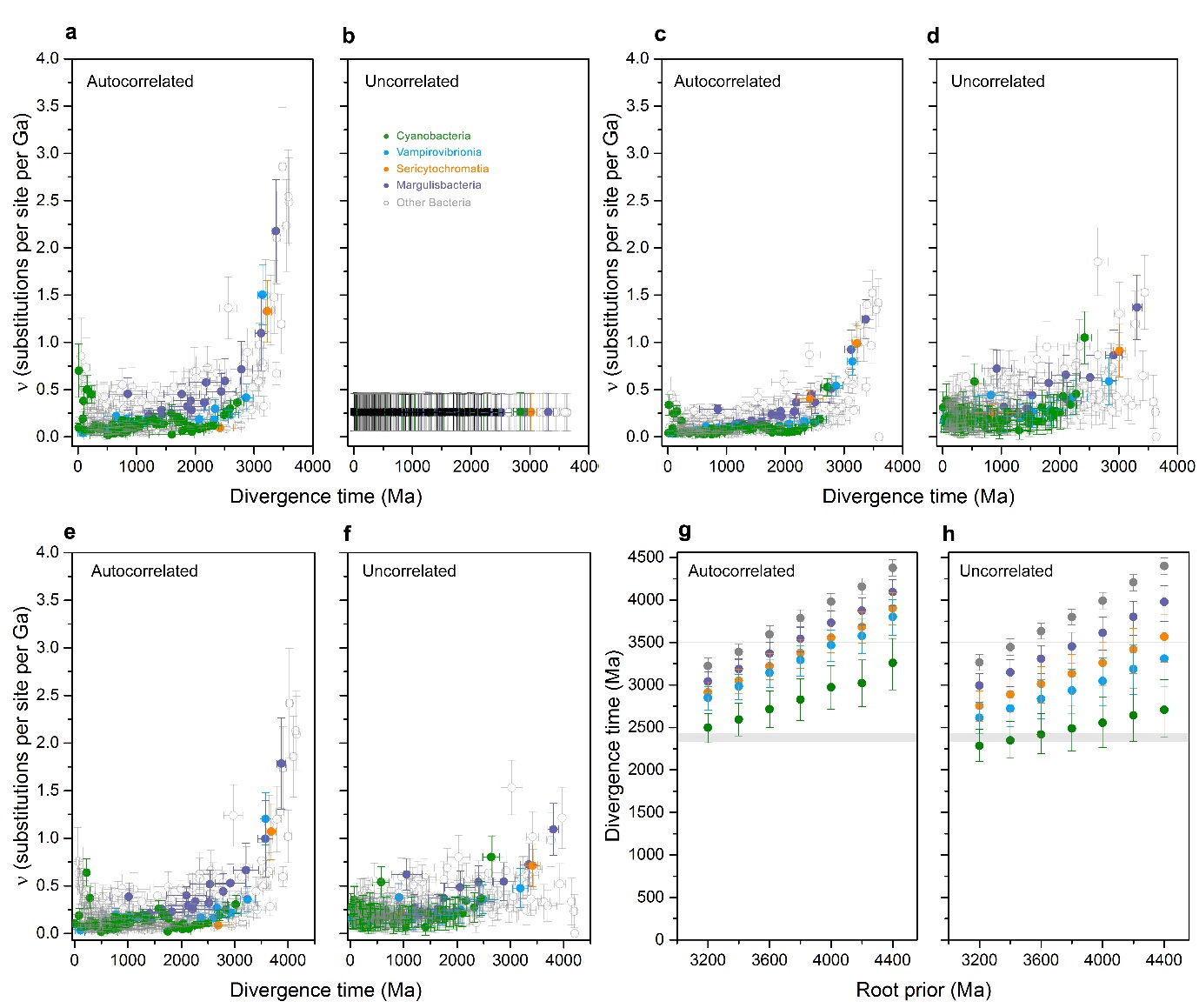

**Supplementary Figure S7.** Comparison of the rates of evolution determined using an autocorrelated and uncorrelated model of rate change for the bacterial RpoB sequence dataset. Conceptually, autocorrelated models assume that rates of evolution can vary across clades, but closely related clades are more likely to evolve at similar rates. Uncorrelated models assume that each branch of the tree can evolve at a rate that is independent from any other [8]. **a** Rates at internal node (instant rates) as a function of divergence time computed with a log normal autocorrelated relaxed clock. The root age was fixed at 3.6 Ga using a gamma root prior with a narrow standard deviation of 0.05 Ga. The tree was calibrated as described in the Materials and Methods. **b** Rates at internal nodes (instant rates) computed with an uncorrelated gamma model and a root prior of 3.6 ± 0.05 Ga. This model assumes that the rate assigned at every internal node of the tree is equal to the total average rate. **c** Average rate of evolution across internal branches of the same tree used to extract rates in panel **a**. The root has no length and therefore its rate is 0. **d** Average rate of evolution across internal branches of the same tree used to extract rates in panel **b**. A similar trend of decreasing rates through the Archean is observed, but a greater scatter is seen in comparison with an autocorrelated model. **e** Instant rates from an autocorrelated relaxed clock identical to **a** but using a root prior of 4.2 ± 0.05 Ga. **f** Average internal branch rates from an uncorrelated clock identical to **d**, but using a root prior of 4.2 ± 0.05 Ga. Panels **c** to **f** show that high rates of evolution are required during the Archean regardless of when exactly the domain Bacteria originated. The younger the estimated age of Bacteria, the faster the rates required at the initial stages of diversification. Panels **g** and **h** compare the estimated divergence times of Thermotogae (grey), Margulisbacteria (violet), Sericytochromatia (orange), Vampirovibrionia (blue), and Cyanobacteria (green) for the two models using identical calibrations, but varying the root prior from 3.2 to 4.4 ± 0.05 Ga. The estimated ages are dependent on the age of the root, however, the span of time between ancestral nodes remains somewhat constant. There is a greater spread of dates generated with the uncorrelated gamma model. We attribute this difference to the assumption that all internal nodes evolve at the same rate, which is not necessarily reflective of true evolutionary processes. Error bars on the nodes represent standard error on mean values.

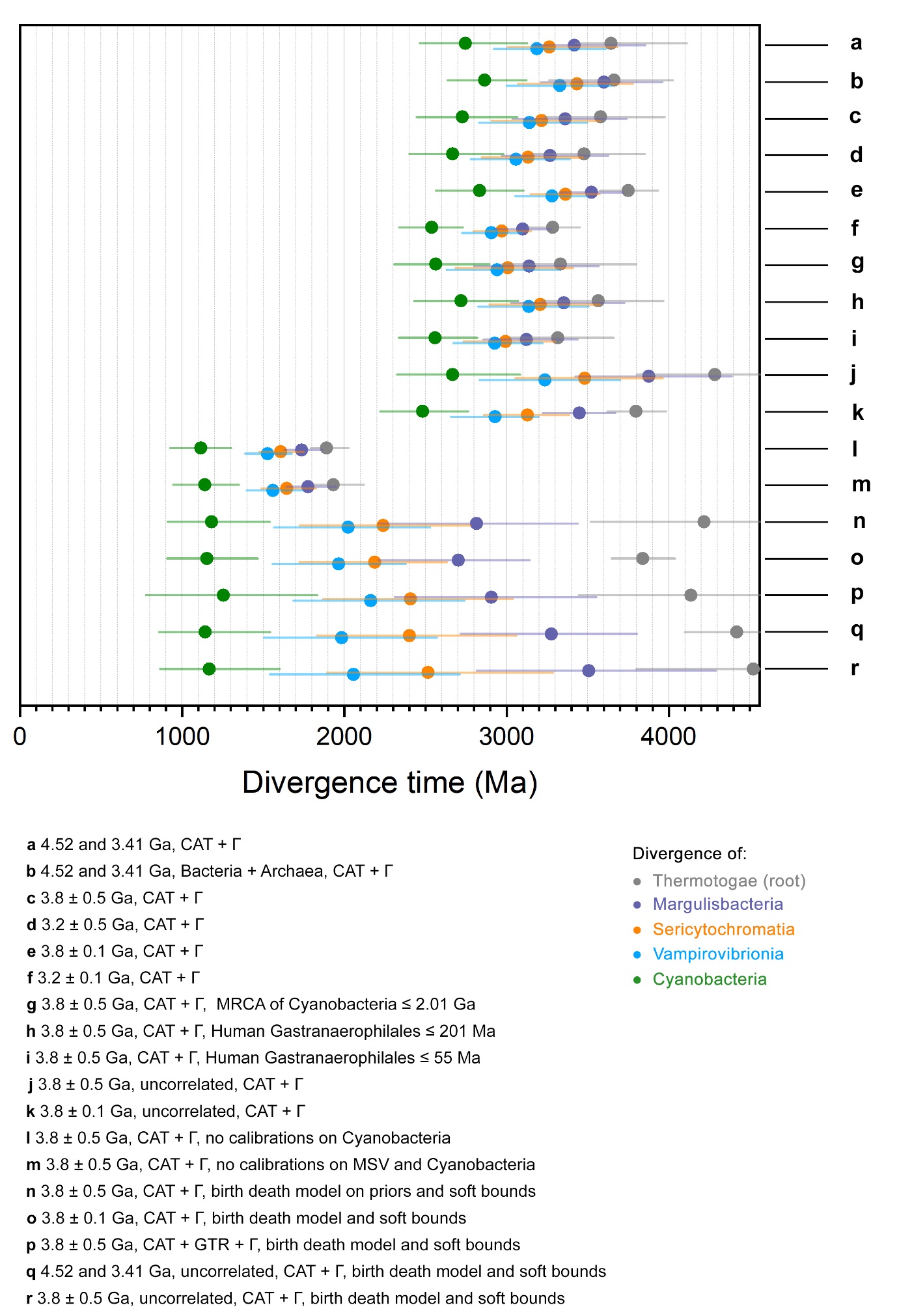

**Supplementary Figure S8.** Molecular clock **s**ensitivity analysis. Each row represents an independently performed molecular clock using various models as listed from **a** to **p** on the RpoB dataset. Each coloured dot represents the estimated divergence time for each respective node as indicated in the figure. Error bars represent 95% confidence intervals. Model **a** was calculated using a calibration on the root with a maximum of 4.52 Ga and a minimum of 3.41 Ga and applying a log normal autocorrelated clock with CAT+Γ. Model **b** was calculated under similar parameters as **a** but used the dataset that included sequences from Archaea. All other models used the sequence dataset that included only bacterial sequences. Model **c**, instead of a calibration on the root, used a gamma root prior with average of 3.8 Ga and a standard deviation (s.d.) of 0.5 Ga. Models **a** to **c** generated largely consistent results indicating that the root constraints are sufficiently broad to allow divergence times to converge towards similar ages.

Models **c** to **e** varied the gamma root prior and its s.d. while maintaining all calibrations and settings intact, to understand how the rates of evolution and ages would vary in connection to a younger or an older root. In model **c**, the prior was set to 3.8 ± 0.5 Ga, which is a relatively flexible constraint. This produced a posterior mean age on the root of 3.57 Ga (95% CI: 3.23 – 3.97 Ga). In model **d**, the prior average was changed to 3.2 Ga, while maintaining the broad s.d. of 0.5 Ga. This produced a posterior age of 3.47 Ga (95% CI: 3.16 – 3.85 Ga). Thus, when a flexible constraint on the root is used, the estimated mean age on the root converges towards 3.5-3.6 Ga, as in model **a** or **b**. Now, if the s.d. is made more restrictive (0.1 Ga), then the posterior age would be fixed within the set range. This allowed us to simulate other evolutionary scenarios. For example, model **e**, with a root prior of 3.2 ± 0.1 Ga, generated a posterior age of 3.28 Ga (95% CI: 3.13 – 3.45 Ga). Model **f** with a root prior of 3.8 ± 0.1 Ga, generated a posterior age of 3.75 Ga (95% CI: 3.57 – 3.93 Ga). It is unclear when exactly the domain Bacteria started to diversify. Therefore, model **e** would assume that Bacteria started to radiate late relative to **f**. We found that the span of time between the MRCA of Cyanobacteria and the root was found to be in the range of 0.75 for the most restrictive and young prior (**e**) to 0.91 Ga for the most restrictive and old prior (**f**), while the span of time between Cyanobacteria and Vampirovibrionia varied from 0.36 Ga (**e**) to 0.44 Ga (**f**). The latter being the same as in model **a** and **b** (see Figure 5 and Supplementary Figure S9a).

Model **g** varied instead a calibration point. In this case, the MRCA of Cyanobacteria was restricted with a maximum age of 2.01 Ga with no minimum age; and did not include the other two cyanobacterial calibrations described in the Materials and Methods (node 17 and 21). However, the estimated mean age for this node was 2.56 Ga (95% CI: 2.30 – 2.89 Ga). This change in calibration resulted in ages that were overall about 0.2 – 0.3 Ga younger for the deepest nodes relative to our benchmarking models, which did not include a maximum age for the MRCA of Cyanobacteria (e.g. **a**, **c**). This effect is a consequence of overall faster calculated rates. The span of time between Cyanobacteria and Vampirovibrionia was found to be 0.37 Ga and that between the root node and Cyanobacteria was found to be 0.76 Ga. A similar effect was observed when Vampirovibrionia are restricted to faster evolutionary rates instead (**h** and **i**), but maintaining the cyanobacterial calibrations (node 17 and 21). For example, in model **i**, the span of time between Vampirovibrionia and Cyanobacteria was calculated to be 0.36 Ga and between the root node and Cyanobacteria 0.75 Ga. The effect was less strong for model **h**, but followed a similar trend.

Uncorrelated gamma models (**j** and **k**) resulted in spread-out divergence time estimates (see also the preceding figure). If it is considered that this model is a better representation of the changes in the rates of evolution as a function of time, then this would imply a less explosive diversification of bacterial clades in contrast to what has been suggested in other independent analyses [9, 10]. Removal of all calibrations on Cyanobacteria and MSV resulted in collapsed age estimates (**l** and **m**). Super-relaxed clock models that implemented soft bounds (**n** to **r**) resulted on a very large spread of ages that appear unrealistic, regardless of whether an autocorrelated and uncorrelated rate model was used, by mimic the results in ref. [11] that used a similar algorithm, although this dataset did not include an Archaeal outgroup. We attribute this effect to the non-parametric rate smoothing method employed, which attempts to minimize differences in substitution rates among lineages (birth and death, soft bounds) [8]. It should be noted that soft bounds were originally developed for datasets featuring largely homogenous rates of protein evolution, such as highly-conserved mitochondrial proteins in primates, showing variation in sequence identity of just below 2%, and were not anticipated to be used on molecular clocks that contained representatives from all domains of life [12].

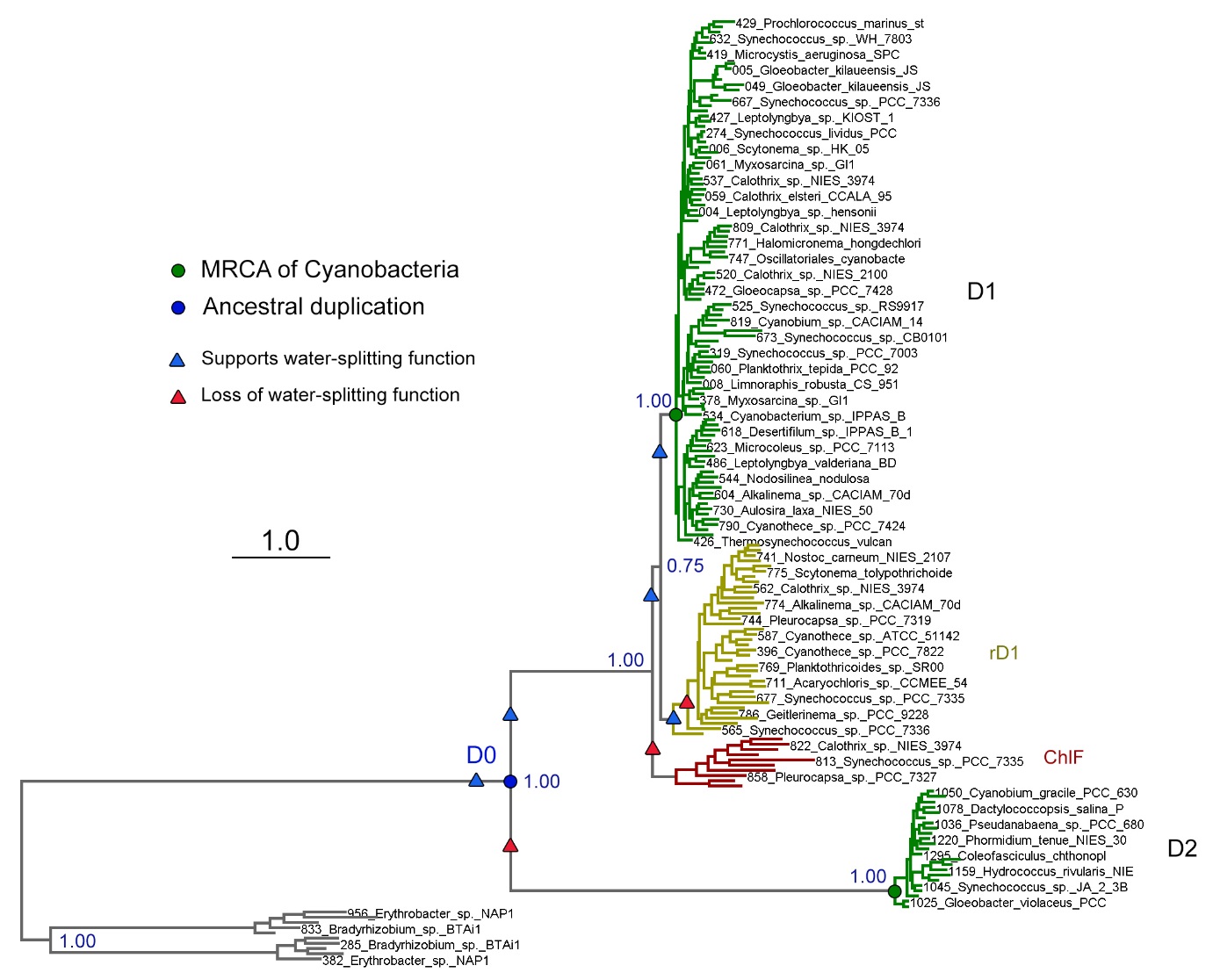

**Supplementary Figure S9.** ML tree of D1 and D2 sequences used for Ancestral Sequence Reconstruction and as described in the Materials and Methods. Green dots mark the standard D1 and D2 sequences inherited by the MRCA of Cyanobacteria [4]. Coloured triangles mark predicted capacity for supporting water oxidation as assessed by the presence or absence of ligands to the Mn_4_CaO_5_ cluster and overall similarity to D1. ChlF and rD1 are atypical D1 forms, characterized by lack of some, but not all, ligands to the Mn_4_CaO_5_ cluster as described in Cardona et al., before [13]. ChlF denotes the “super-rogue” D1 (Group 1 D1) or chlorophyll *f* synthase [14]. The clade denoted rD1 (“rogue” D1) is more widely distributed than ChlF. It has been proposed previously that this D1 has the role of “switching off” PSII in the dark [15] or under other conditions. We found that the ancestrally reconstructed sequence to all rD1 had a full set of ligands to the Mn_4_CaO_5_ cluster, with glutamate at position equivalent to 170 instead of aspartate. Furthermore, the earliest branching sequence in this clade was found to be that of *Halothece* sp. PCC 7418 (WP_015224801.1), which had a full set of ligands with E170 and reproduced the analysis in ref. [14], but this was not noted then. The ancestral sequence to ChlF was found to lack only a single ligand to the cluster, H332 and also featured E170. This is consistent with a scenario in which all atypical D1 sequences emerged from a D1 that was capable of water oxidation. All reconstructed sequences are freely available in the repository as indicated in the Materials and Methods or on request to the corresponding author.

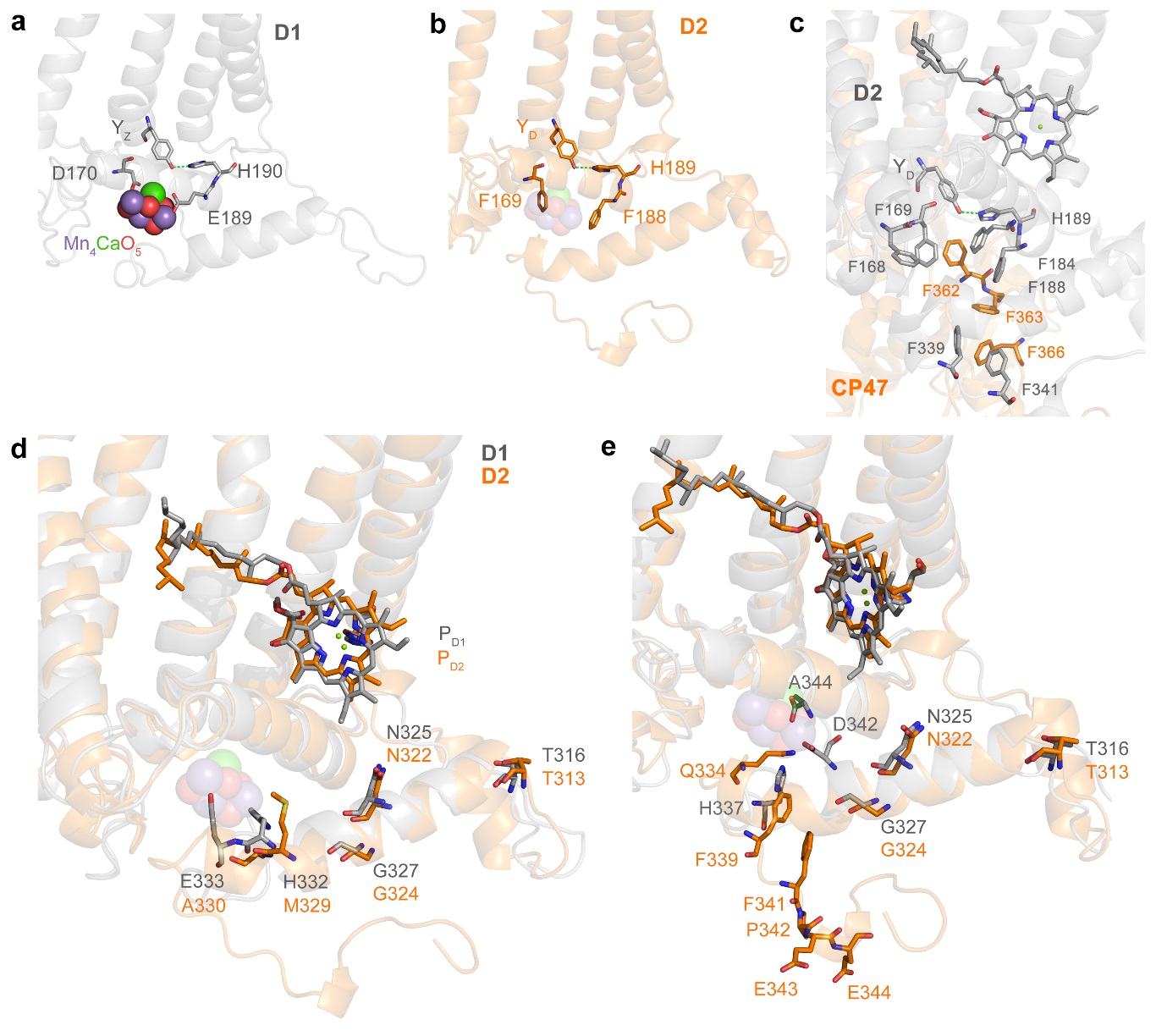
**Supplementary Figure S10.** Comparison of the electron donor site of D1 and D2. **a** The Mn_4_CaO_5_ cluster and associated redox Y_Z_-H190 pair showing the inner ligands D170 and E189. **b** The electron donor site of D2 showing the redox active Y_D_-H189 pair and the phenylalanine residues that occupy positions homologous to D1-D170 and E189. The equivalent position of the Mn_4_CaO_5_ cluster is shown transparently from an overlap with D1. **c** Entire patch of phenylalanine residues blocking access to Y_D_-H189. **d** Overlap of D1 and D2. Residues occupying homologous positions are shown as sticks and assuming no indels after conserved residues D1-G327 and D2-G324. **e** Overlap of D1 and D2 focusing on the C-terminal residues. Assuming no indels, ligand D1-H337 would be equivalent to D2-Q334, D1-D342 to D2-F339, and D1-A344 to D2-F341. The codons of the *psbD* gene (D2) that encodes P342, E434, and E344 overlap with the Shine-Dalgarno ribosomal binding site of *psbC* (CP43).

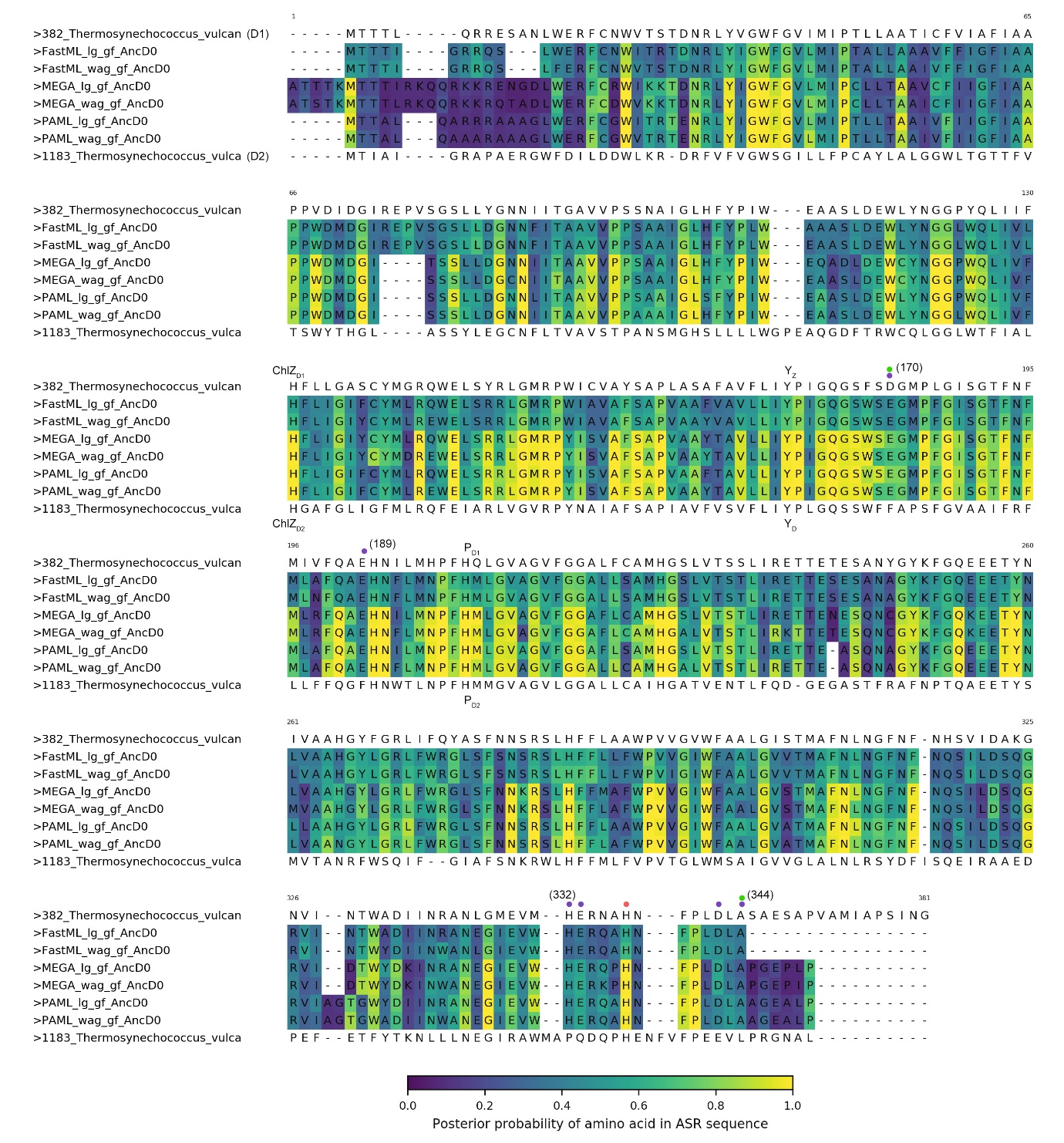

**Supplementary Figure S11.** Posterior probabilities (PP) of selected ancestrally reconstructed D0 sequences. The top and bottom sequences represent the D1 (PsbA1) and D2 subunits of *Thermosynechococcus vulcanus*. The colour at each site denotes the PP for the most likely residue as shown in the colour map at the bottom of the alignment. Ligands to the Mn_4_CaO_5_ cluster are highlighted with coloured dots: purple for Mn, green for Ca, and red for O. Y_Z_ and Y_D_ denote the redox active tyrosine residues; P_D1_ and P_D2_ denote the ligands to the chlorophylls equivalent to the “special pair”; ChlZ_D1_ and ChlZ_D2_ denote the ligands to these antenna chlorophylls.

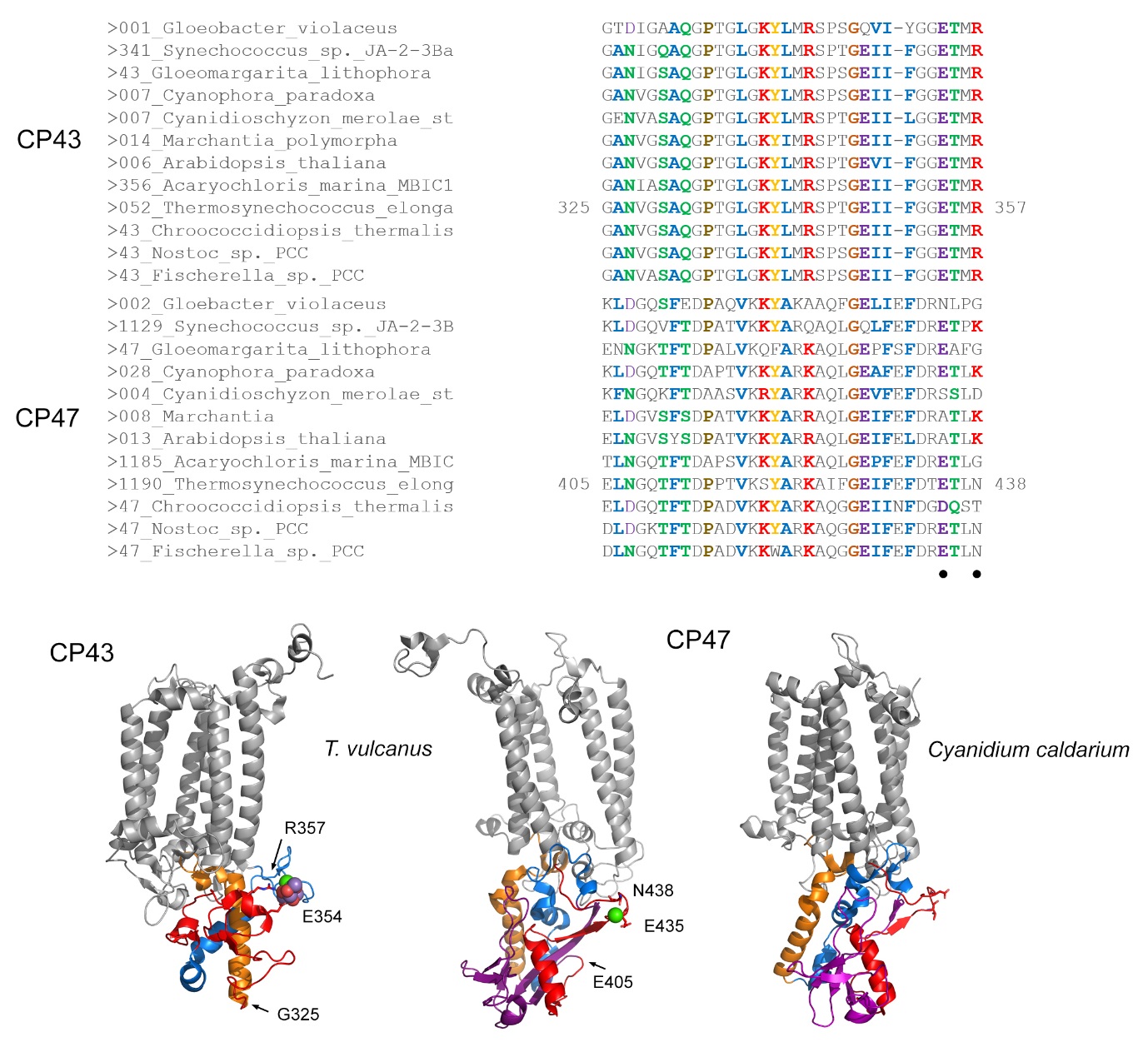

**Supplementary Figure S12.** Comparison of the region denoted as EF_3_ in the extrinsic domain of CP43 and CP47. The sequence alignments compare the protein fold highlighted in red ribbons on the structures below and as indicated with arrows. Sequence identity is detected indicating a swap of position in one subunit relative to the other. A Ca is bound in CP47 to E435 and N438, which are predicted to be at homologous positions to the Mn_4_CaO_5_ cluster ligands CP43-E354 and R357. This Ca site is not observed in the 2.77 Å crystal structure of the red algae *Cyanidium caldarium* [16].

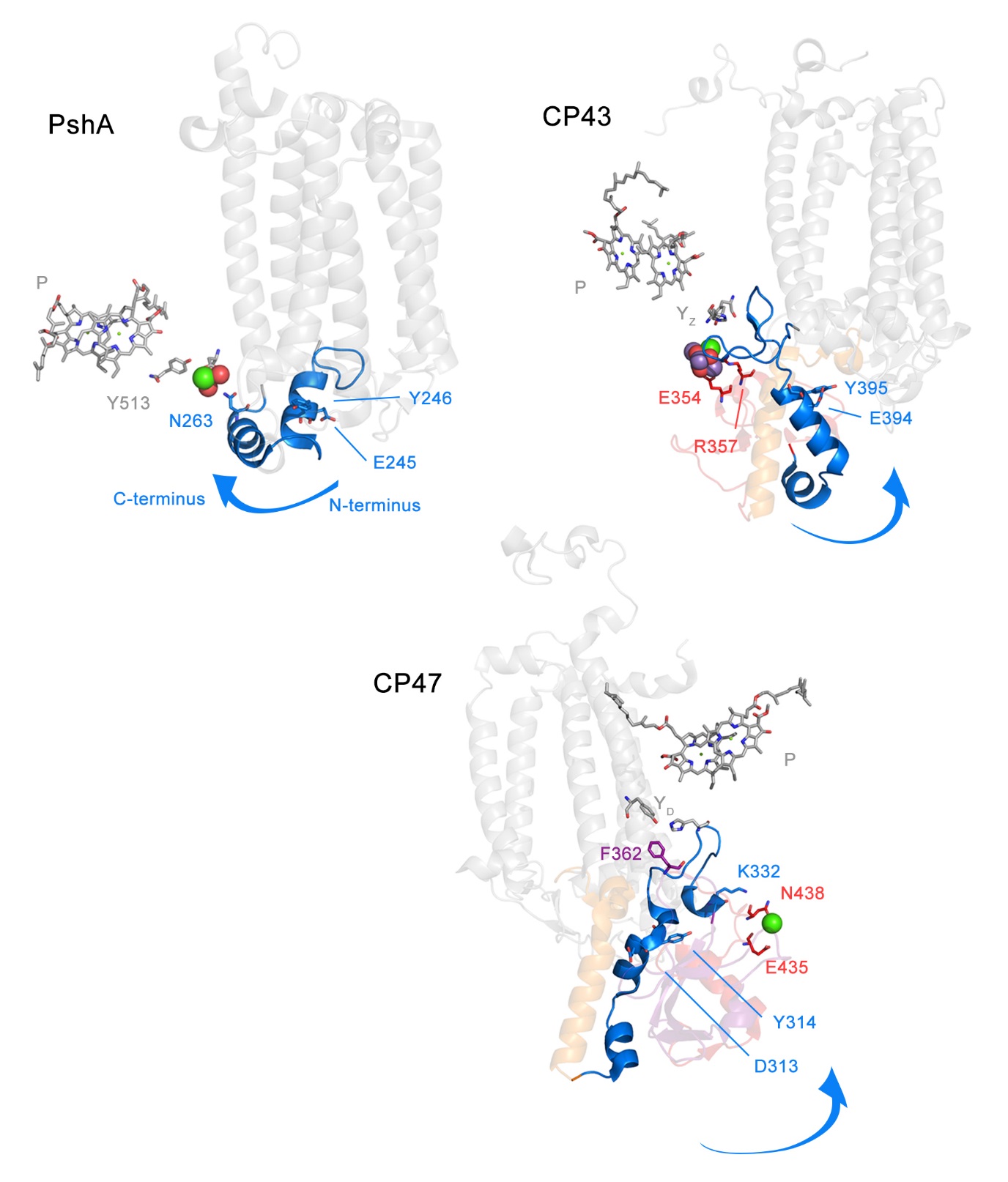

**Supplementary Figure S13.** Structural comparison of region denoted as EF_1_ in the extrinsic domain of heliobacterial PshA, CP43 and CP47. The fold is characterized by two small alpha helix and although there is little sequence conservation between PshA and the cyanobacterial sequences, the fold contain a potentially conserved and characteristic EY or DY pair. The blue arrow indicates the direction of the fold from N- to C-terminus.

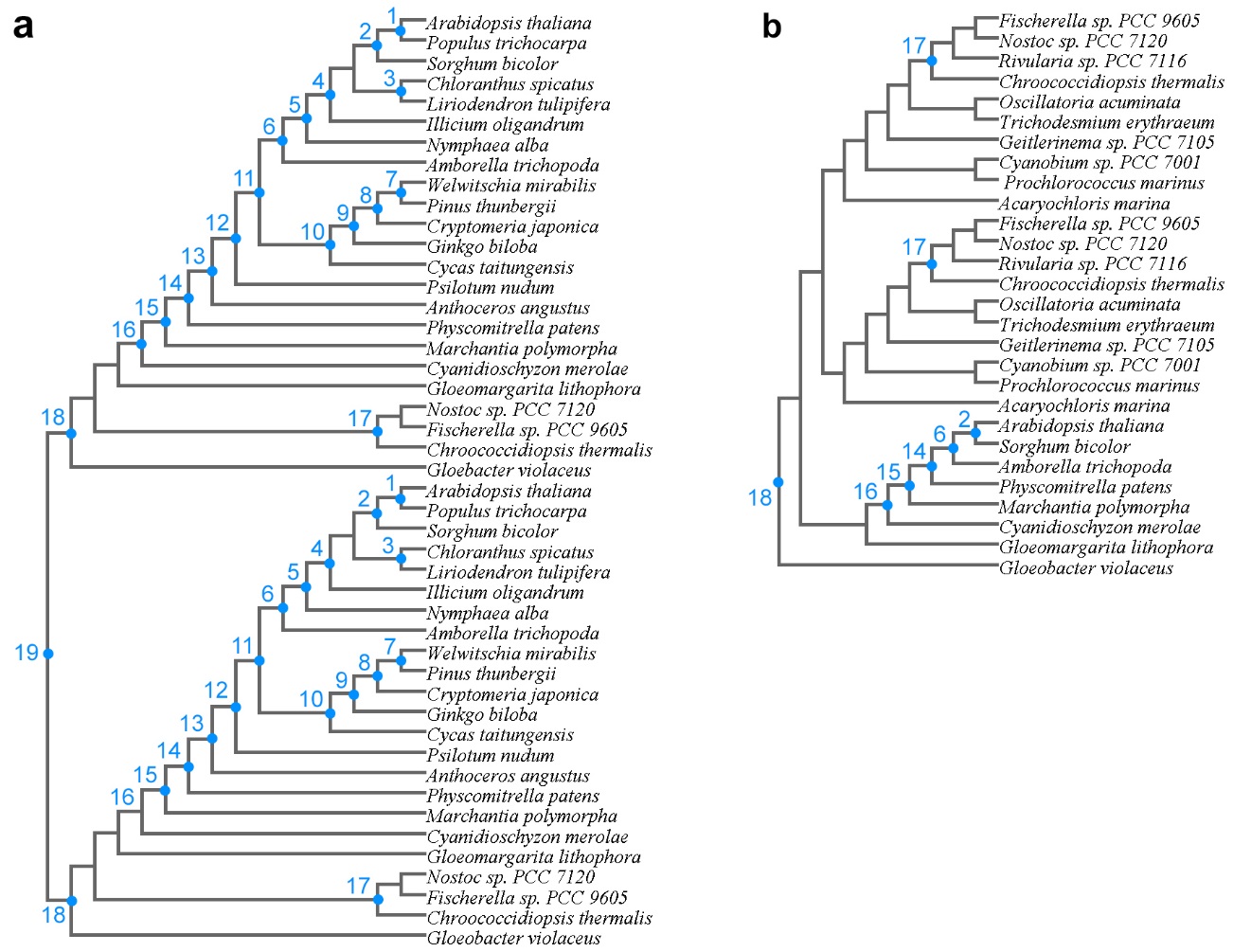

**Supplementary Figure S14.** Calibration points for the determination of rates of evolution. **a** Topology used for of CP43/CP47 and Alpha/Beta. **b** Topology used for FtsH subunits. Calibrated nodes are labelled, ages were set as described in the Materials and Methods, and listed in Table 2 using the respective labels.

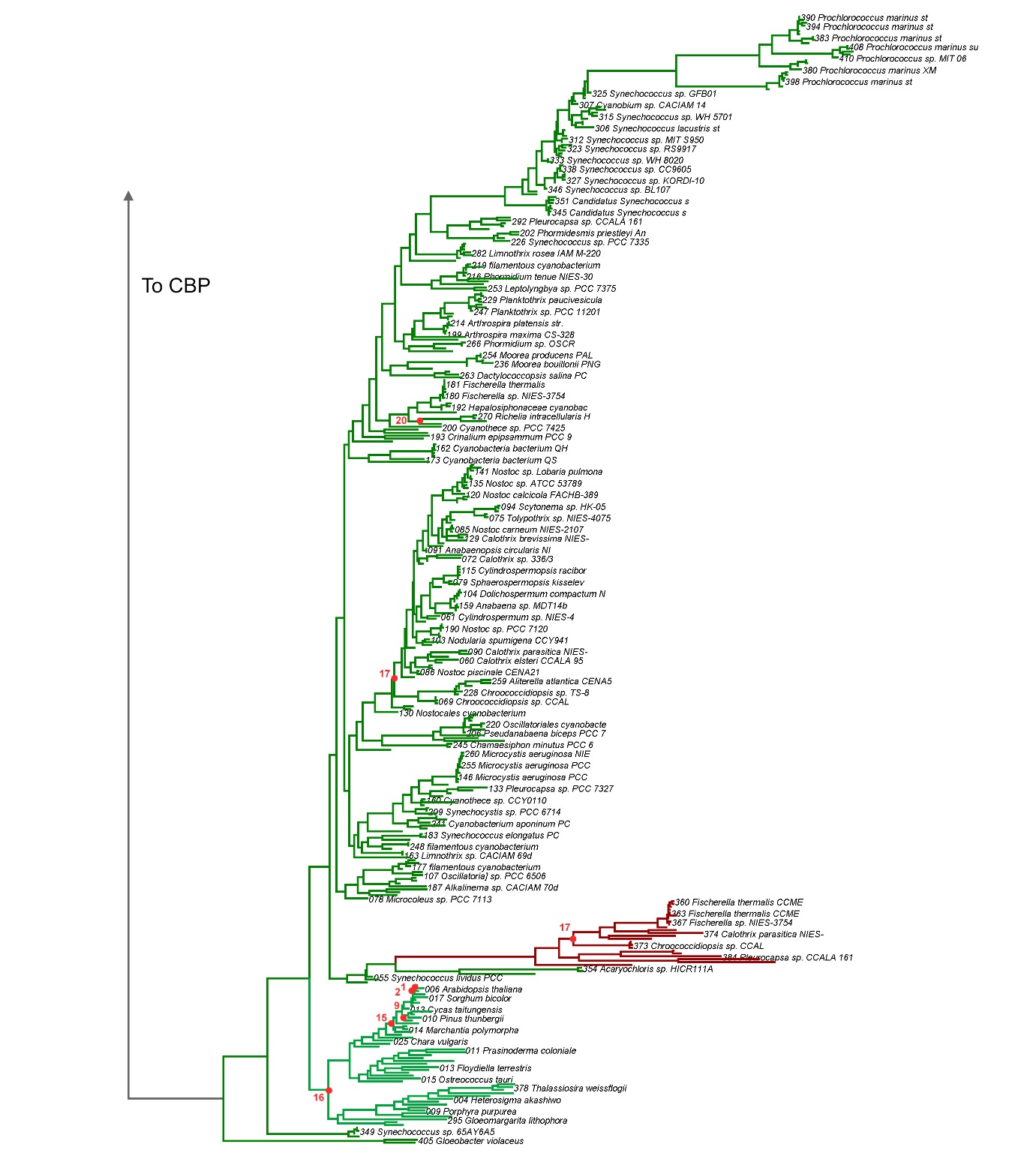

**Supplementary Figure S15 (Part 1).** Calibrations used to determine rates of evolution in the large phylogeny of CP43 and CBP. This part of the tree shows CP43 subunits. Calibrated nodes are labelled as listed in Table 2.

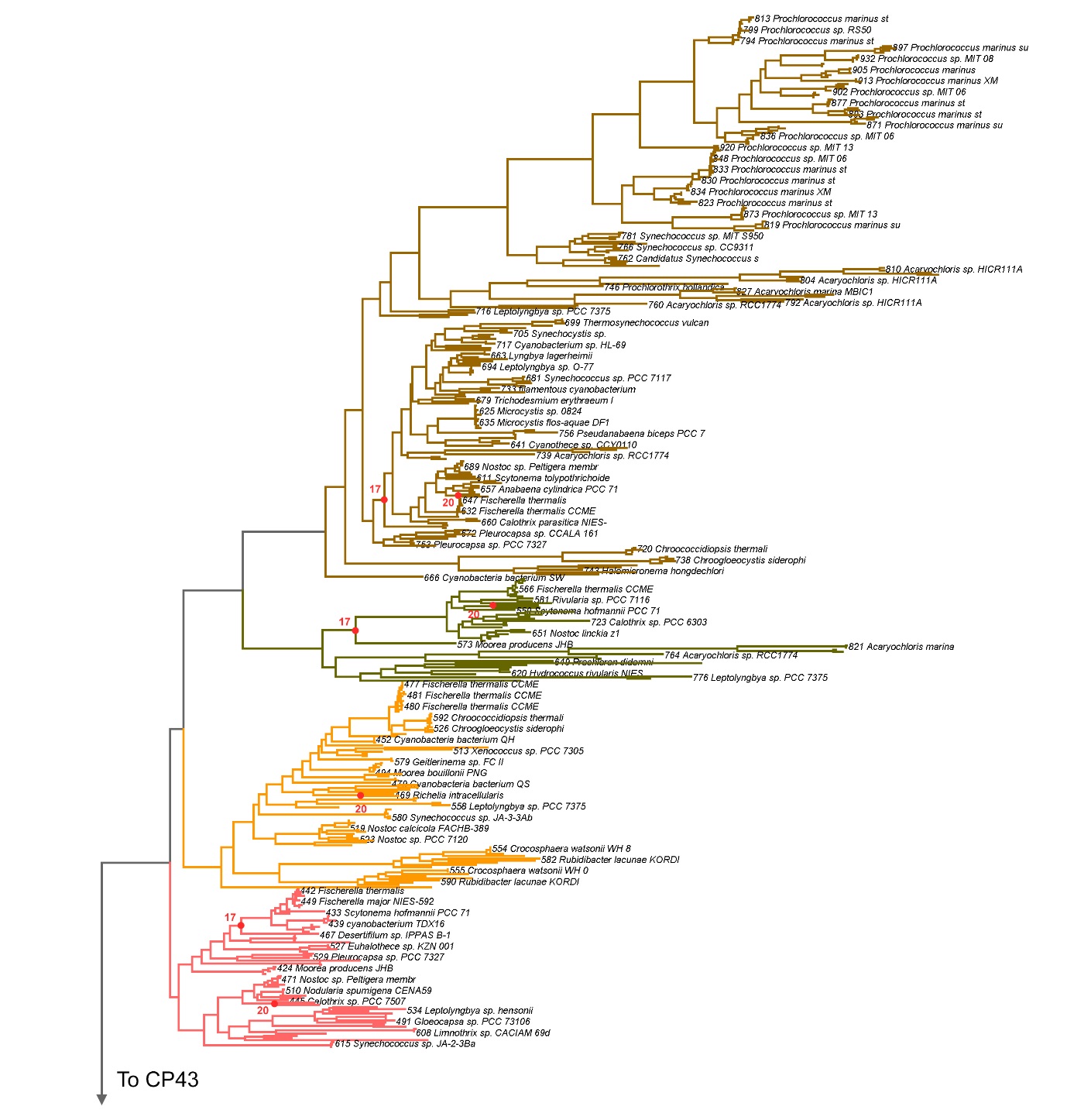

**Supplementary Figure S15 (Part 2).** Calibrations used to determine rates of evolution in the large phylogeny of CP43 and CBP. This part of the tree shows CBP subunits. Calibrated nodes are labelled as listed in Table 2.

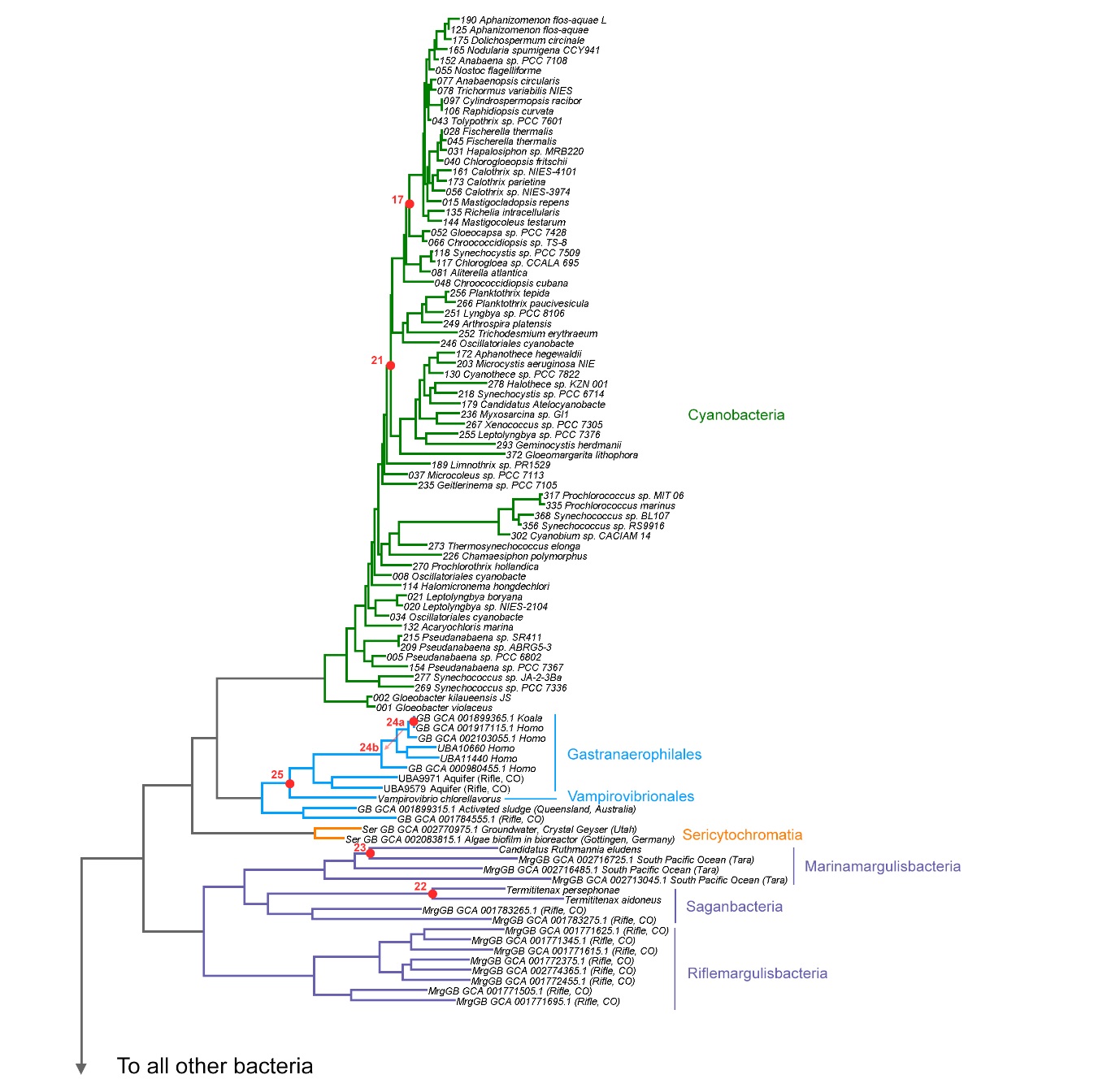

**Supplementary Figure S16 (Part 1).** Calibrations implemented on the phylogeny of bacterial RpoB subunits. This part of the tree shows Cyanobacteria, Margulisbacteria, Sericytochromatia, and Vampirovibrionia. Calibrated nodes are labelled as listed in Table 2 and justified in the Materials and Methods.

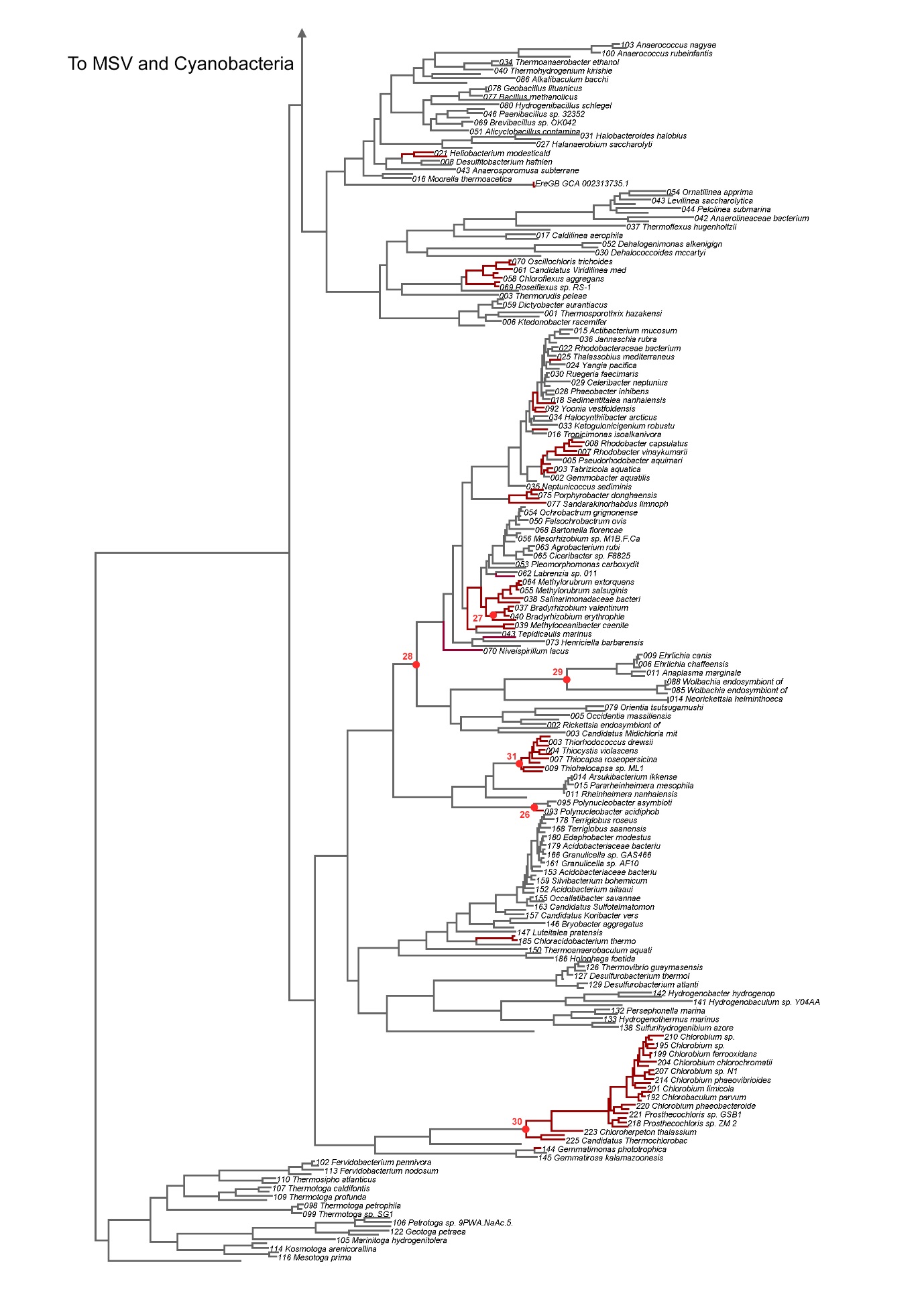

**Supplementary Figure S16 (Part 2).** Calibrations implemented on the phylogeny of bacteria RpoB subunits. This part of the tree shows all other bacterial sequences in the same dataset. Red branches mark phototrophic clades and strains. Calibrated nodes are labelled as listed in Table 2 and justified in the Materials and Methods.

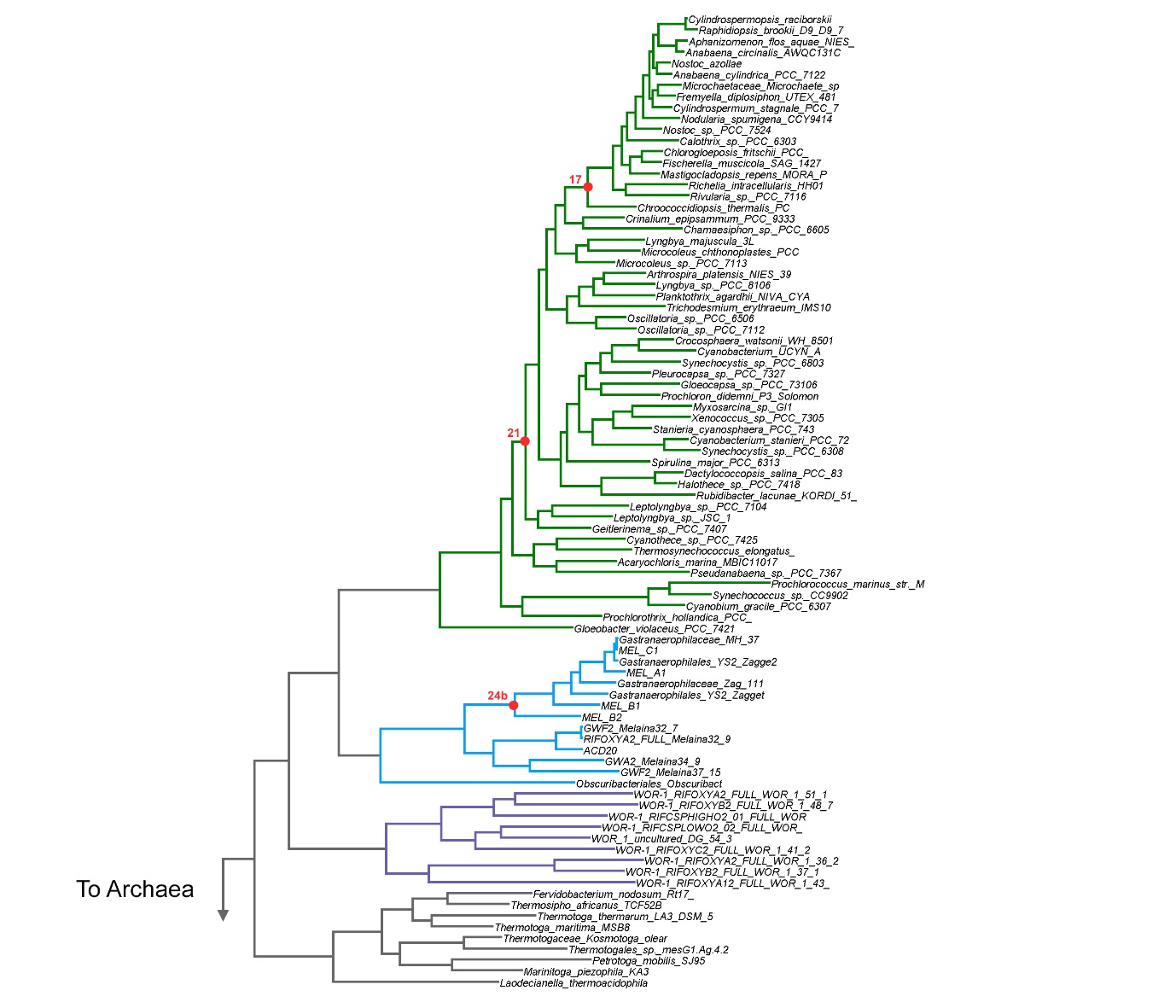

**Supplementary Figure S17.** Calibrations implemented on the phylogeny of concatenated ribosomal protein sequences. This part of the tree shows the calibrated bacterial sequences. Calibrated nodes are labelled as listed in Table 2 and justified in the Materials and Methods.

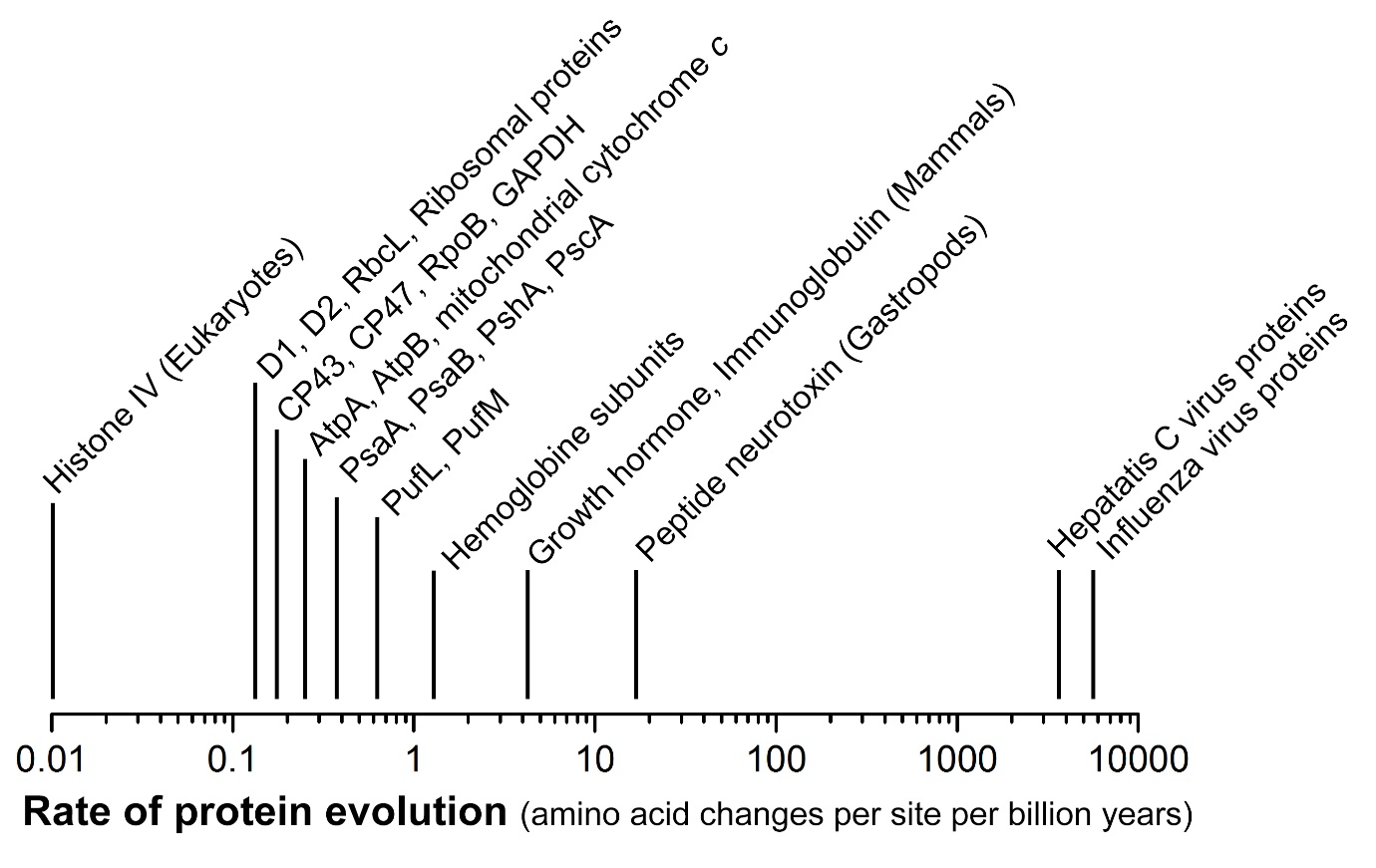

**Supplementary Text S1. Rates of protein evolution in context**

Proteins display rates of amino acid substitutions that spans nearly six orders of magnitude. Take two identical sequences: if one of them evolves at a rate of 1 amino acid substitution per site per billion years and the other does not change, and assuming that all sites have a similar chance of mutating, it would take 1 Ga for each site to have changed once in the evolving sequence. This means that it would take under 1 Ga for these two hypothetical sequences two lose all sequence identity. Alternatively, if both sequence are evolving independently from each other at *that* same rate, it would take less than half that time, 500 Ma, for both sequences to lose all sequence identity, because both are accumulating change [17].

If two proteins are evolving at 10 amino acid substitutions per site per Ga, assuming the same hypothetical conditions in which all sites and all substitutions are equally probable, it would take less than 50 Ma for the two identical sequences to lose all trace of identity. On the other hand, if the two sequences were evolving at 0.1 amino acid substitutions per site per Ga, it would take 5 Ga for them to lose all sequence identity. That means that after a period of 2.5 Ga of evolution, these two slowly evolving sequences would still retain about 50% sequence identity. Therefore, most highly-conserved proteins across the tree of life, that are predicted to be very ancient, tracing back to the LUCA or the earliest forms of life, such as ATP synthase, RNA polymerase, or the ribosome; proteins that have retained substantial sequence identity over a multibillion year history, evolve at rates between well below 1, and above 0.1 amino substitution per site per Ga. In this way, for example, comparing the protein sequence of RNA polymerase subunit B (RpoB) from two very distantly related bacteria such as *Thermotoga* (Thermotogae) and *Thermosynechococcus* (Cyanobacteria) the level of sequence identity is about 40%, including gap and insertions. If we consider that the phyla containing these two clades diverged over 2.5 Ga ago, it can be expected that these sequences have been evolving at rates on average somewhat above 0.1 amino acid changes per site per Ga, but certainly well below 1.

In contrast, fast evolving proteins, like those found in viruses, that accumulate change in just a few years, display rates of evolution of several thousand changes per site per billion years. At the other extreme, one of the slowest evolving protein known is histone subunit H4, a protein of about 100 residues that has barely changed at all within the past billion years, showing just 2 amino acid changes between *Arabidopsis* and *Homo sapiens*, for example.

Molecular clocks will use a protein sequence alignment and some known time constraints (calibrations) to estimate and extrapolate the rates of protein evolution in amino acid changes per site per unit of time, across a phylogenetic tree, much in the same way that we have considered in the preceding paragraphs. Thus, knowing the *distance* between two sequences (number of amino acid changes per site), and the rate of protein evolution (the *speed* of sequence change), it is possible to estimate the divergence *time*. Modern molecular clocks with the implementation of Maximum Likelihood or Bayesian Inference methods can account for the uncertainty generated by sparse calibration points, and the fact that in reality different clades, different sequences, and different parts in a sequence evolve at different rates.

In the image above, the rates of evolution for D1, D2 , PufL and PufM were taken from ref. [4]. Those for PsaA, PsaB, PshA, PscA from ref. [5]. For CP43, CP47, AtpA, AtpB, RpoB and ribosomal proteins were taken from this study. For histone IV, GAPDH, cytochrome *c*, hemoglobine, growth hormone and immunoglobuline and other not shown here, see ref. [18]. For gastropod neurotoxins and viral proteins, see also [4] and references therein.

**Supplementary Text S2. Over- and underestimation of the rates of evolution**

It is understood that these very ancient systems (e.g. ATP synthase, RNA polymerase, ribosomes) have likely reached substitutional saturation at their longest distance. In consequence, the distance between Alpha and Beta, and between Archaea and Bacteria, is thought to be largely underestimated [19, 20]. This seems to be also true for the distance between CP43 and CP47, and between D1 and D2 [4]. Therefore, the rates of amino acid substitutions at the oldest node are more likely to be underestimated (slower than they should be) rather than overestimated, and therefore, ΔT is more likely to be of larger magnitude than otherwise. Furthermore, we suspect that the rates of evolution of Margulisbacteria and Vampirovibrionia might still be underestimated (slower than real rates), while that of Cyanobacterial overestimated, due to the relaxed nature of molecular clocks, which tends to smooth rates out [8]. For example, Gastranaerophilales, were timed to have originated long before the emergence of animals for both RpoB (95% CI: 1.91 - 2.74 Ga) and the ribosomal proteins dataset (95% CI: 0.89 - 1.41 Ga). It is not implausible that strains of Margulisbacteria and Vampirovibrionia have experienced faster rates of evolution than Cyanobacteria as many members of this groups have complex symbiotic lifestyles with other bacteria, eukaryotes, or both [21, 22]. Some of these strains have experienced genome size reductions, which is usually correlated with accelerated rates of change [23]. This dichotomy in the rates was also detected for Rickettsidae (0.40 ± 0.11 δ Ga^-1^) when compared to Caulobacteridae (0.17 ± 0.05 δ Ga^-1^) of the Alphaproteobacteria. We also noted the same likely underestimation for many strains of Rickettsidae, which appeared to be much older than their symbiotic associations would suggest despite the provided constraints. If this is indeed the case, and the clocks underestimate the rates of the faster evolving clades, then it is likely that the events leading to the divergence of MSV occurred more rapidly than anticipated from this and previous molecular clocks. To overcome this limitation accurate measurements of the rates of protein or genome evolution of a representative number of clades across the trees must be obtained to complement, validate or entirely supersede, calibrations on deep-time molecular clocks of prokaryotes.
